# Genetic, epigenetic, and environmental mechanisms govern allele-specific gene expression

**DOI:** 10.1101/2021.09.09.459642

**Authors:** Celine L St. Pierre, Juan F Macias-Velasco, Jessica P Wayhart, Li Yin, Clay F Semenkovich, Heather A Lawson

**Author notes:** Corresponding author 660 South Euclid Ave, Campus Box 8232, Saint Louis, MO, 63110.

## Abstract

Allele-specific expression (**ASE**) is a phenomenon where one allele is preferentially expressed over the other. Genetic and epigenetic factors cause ASE by altering the final allelic composition of a gene’s product, leading to expression imbalances that can have functional consequences on phenotypes. Environmental signals also impact allele-specific gene regulation, but how they contribute to this crosstalk remains understudied. Here, we explored how allelic genotype, parent-of-origin, tissue type, sex, and dietary fat simultaneously influence ASE biases in a F_1_ reciprocal cross mouse model. Male and female mice from a F_1_ reciprocal cross of the LG/J and SM/J strains were fed a high fat or low fat diet. We harnessed strain-specific variants to distinguish between two classes of ASE: parent-of-origin dependent (unequal expression based on an allele’s parental origin) and sequence dependent (unequal expression based on an allele’s nucleotide identity). We present a comprehensive map of ASE patterns in 2,853 genes across three metabolically-relevant tissues and nine environmental contexts. We found that both ASE classes are highly dependent on tissue type and environmental context. They vary across metabolic tissues, between males and females, and in response to dietary fat levels. Surprisingly, we found 45 genes with inconsistent ASE biases that switched direction across tissues and/or contexts (e.g. SM/J biased in one cohort, LG/J biased in another). We also integrated ASE and QTL data from populations at various degrees of intercrossing the LG/J and SM/J strains. ASE genes in these tissues are often enriched in QTLs for metabolic and musculoskeletal traits, highlighting how this orthogonal approach can prioritize candidate genes for functional validation. Together, our results provide novel insights into how genetic, epigenetic, and environmental mechanisms govern allele-specific gene regulation, which is an essential step towards deciphering the genotype to phenotype map.

## INTRODUCTION

Deciphering the genotype to phenotype map remains a fundamental quest in biology. Gene expression is a promising focal point, as it is an intermediate step between DNA sequence and gross phenotype. Gene expression itself is a complex trait that is regulated by genetic, epigenetic, and environmental factors (Pastinen 2010). Recent efforts to characterize how genes are regulated have often focused on mapping expression quantitative trait loci (**eQTLs**), which identify genetic variants linked to changes in gene expression at the population level (Cookson et al. 2009). However, these studies can only interrogate total gene expression and assume that genes are biallelically expressed (i.e. both alleles are equally expressed), which may mask underlying regulatory mechanisms. Furthermore, epigenetic changes and gene-by-environment interactions can alter gene expression patterns in a dynamic and tissue-specific manner without modifying the underlying nucleotide sequence; these effects are missed in a typical sequence-based method. An allele-specific approach is required to directly measure how *cis*-regulatory variation impacts gene expression and to tease it apart from *trans*-acting factors that affect both chromosomes (Bonasio et al. 2010).

Allele-specific expression (**ASE**) is a phenomenon where a gene’s expression diverges from biallelic. In diploid organisms, one allele is preferentially expressed over the other allele. Previous findings estimate that 30-56% of genes show evidence of allelic imbalance, indicating allele-specific effects have widespread impacts on gene regulation (Ge et al. 2009; Castel et al. 2019; Keane et al. 2011). Depending on a gene’s function, these expression imbalances can lead to phenotypic variation with functional consequences. To detect ASE, single nucleotide polymorphisms (**SNPs**) and other genetic variants are used to distinguish between alleles in heterozygotes and map RNA-sequencing reads to their chromosome of origin (Heap et al. 2010). Allele-specific analyses are a powerful way to exploit a within-sample control (the other allele) to gauge how genetic and epigenetic variation in *cis*-regulatory elements shapes gene expression (Pastinen 2010).

ASE can be divided into two classes: sequence dependent or parent-of-origin dependent (**Figure 1**). Sequence dependent ASE refers to cases where the two alleles are differentially expressed based on their haplotype or nucleotide identity. These patterns are thought to be largely driven by *cis*-acting genetic variants in coding and noncoding regions, such as a premature stop codon that truncates one allele’s transcript or a variant in a promoter region that prevents transcription factors from binding (Keane et al. 2011; Rivas et al. 2015). They can also occur further away from the gene yet still impact its expression, such as motif variations for long-range enhancers or in the sequence context of DNA methyltransferase substrates (Cavalli et al. 2016; Wienholz et al. 2010). Stochastic transcriptional bursts can also cause transient fluctuations in which allele is expressed, resulting in random monoallelic expression that is inconsistent across cells or organisms (Reinius and Sandberg 2015; Deng et al. 2014).

**Figure 1:**
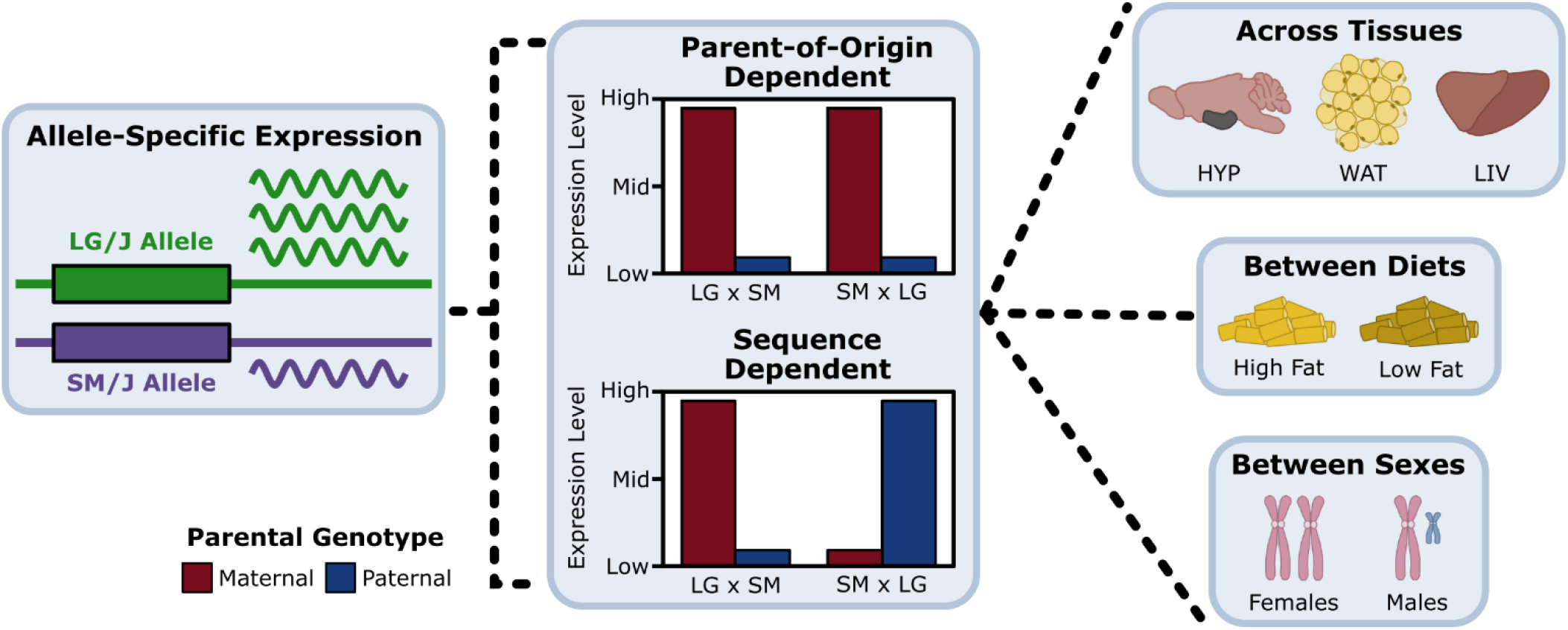
Evaluating parent-of-origin and sequence dependent allele-specific expression across tissues and environmental contexts. We partitioned allele-specific expression (ASE) into its parent-of-origin and sequence effects in a F_1_ reciprocal cross. An example of parent-of-origin dependent ASE is when the maternal allele (red) is preferentially expressed over the paternal allele (blue), regardless of which haplotype contributed it. An example of sequence dependent ASE is when the LG/J allele is preferentially expressed over the SM/J allele, regardless of which parent contributed it. Once we identified significant ASE genes, we compared their expression patterns across metabolic tissues (HYP, WAT, LIV), in response to different diets (high fat, low fat), and between sexes (females, males).

In contrast, parent-of-origin dependent ASE refers to cases where the alleles are differentially expressed based on which parent contributed it, regardless of the underlying sequence. These patterns fall under the umbrella of parent-of-origin effects (**POEs**), a broader class of epigenetic phenomena that manifest as phenotypic differences according to maternal or paternal inheritance (Lawson et al. 2013). The best characterized POE mechanism is genomic imprinting, an extreme case of parent-of-origin dependent ASE where one parent’s allele is completely silenced via selective epigenetic marks, such as DNA methylation and histone modifications (Barlow and Bartolomei 2014; Inoue et al. 2017; Umlauf et al. 2004). Known imprinted genes comprise ∼1% of the human and mouse genomes, yet play important roles in development, metabolism, cognition, and other complex traits (Reik and Walter 2001). Partial imprinting is another, more subtle, form of parent-of-origin dependent ASE that has been documented but is less understood (Wolf et al. 2008; Morcos et al. 2011). Random monoallelic expression can also produce transient and non-heritable parental biases in individual cells through different mechanisms than with imprinting (Gimelbrant et al. 2007; Morcos et al. 2011). Other sources of POEs include maternal effects and gene-specific trinucleotide expansions, but these do not involve ASE (Hager et al. 2008).

Comprehensive atlases of how both classes of ASE vary between tissues and developmental stages have been generated in human and mouse models (Leung et al. 2015; Castel et al. 2019; Babak et al. 2015; Andergassen et al. 2017). These studies reveal that both parent-of-origin and sequence dependent ASE patterns are not always consistent between tissues, indicating that tissue-specific genetic and epigenetic features can mediate allelic imbalances. Additionally, *trans*-acting environmental factors have been shown to interact with *cis*-regulatory variants to modulate the magnitude of ASE effects in human and rice models (Buil et al. 2015; Knowles et al. 2017; Moyerbrailean et al. 2016; Shao et al. 2019). Imprinted genes are also known to be responsive to environmental exposures during fetal development, such as teratogenic agents and maternal nutrition (Kappil et al. 2015). Together, these findings suggest that a complicated crosstalk among genetic variants, epigenetic changes, and environmental signals underlies allele-specific gene regulation, but this model needs to be further investigated.

Here, we explored how allelic genotype, parent-of-origin, tissue type, sex, and dietary fat simultaneously work together to influence allele-specific expression patterns in a F_1_ reciprocal cross of the LG/J and SM/J inbred mouse strains. These strains are uniquely suited for gene-by-environment studies due to their divergent genetic backgrounds and variable responses to dietary nutrition (Nikolskiy et al. 2015; Ehrich et al. 2003; Carson and Lawson 2020; Miranda et al. 2019). These strains are also the progenitors of the longest-running mouse advanced intercross line (**AIL**). An AIL is a multigenerational outbred population produced by intercrossing two inbred strains beyond the F_2_ generation while minimizing relatedness between mates. At each generation, meiotic recombination occurs and further degrades the linkage disequilibrium between adjacent markers, resulting in finer mapping resolutions of quantitative trait loci (**QTL**) (Darvasi and Soller 1995). Since its inception, the LG/J x SM/J AIL has been extensively used to map QTLs related to metabolic, musculoskeletal, behavioral, and physiological traits (Ehrich et al. 2005; Cheverud et al. 2011; Lawson et al. 2010, 2011b, 2011a; Lionikas et al. 2010; Parker et al. 2011; Carroll et al. 2017; Gonzales et al. 2018; Cordero et al. 2019; Zhou et al. 2020). To probe whether allele-specific expression imbalances contribute to complex trait etiology, we integrated our ASE results with these AIL QTL mapping studies and highlight how this orthogonal approach can prioritize potential candidate genes. Untangling the genetic, epigenetic, and environmental mechanisms that govern allele-specific gene regulation is crucial to improving our ability to predict phenotypes from genotypes.

## RESULTS

### Allele-specific expression can be decomposed into parent-of-origin and sequence effects

We measured ASE in a F_1_ reciprocal cross of the LG/J and SM/J inbred mouse strains. Briefly, LG/J mothers were mated with SM/J fathers and vice versa, resulting in F_1_ offspring who are genetically equivalent but differ in the allelic direction of inheritance. Male and female F_1_ mice were fed either a high fat or low fat diet. We obtained RNA-Seq data from three metabolically-relevant tissues: hypothalamus (**HYP**), white adipose (**WAT**), and liver (**LIV**). To explore how dietary fat and/or sex impact ASE patterns, we analyzed nine environmental contexts per tissue: high fat-fed diet (**H**), low fat-fed diet (**L**), females (**F**), males (**M**), high fat-fed females (**HF**), high fat-fed males (**HM**), low fat-fed females (**LF**), low fat-fed males (**LM**), and all contexts collapsed (**All**). Together, these 27 “tissue-by-context” analyses (3 tissues x 9 contexts) allowed us to probe how tissue type and environmental signals influence ASE (**Figure 1**).

The LG/J and SM/J genomes differ by >5 million SNPs and indels (Nikolskiy et al. 2015). To be informative for ASE, a strain-specific variant must be located within a gene’s exome. In this F_1_ model, informative variants are located in 55% of expressed genes in our three tissues and 47% of all genes in the mm10 reference genome (**Supplemental Figure S1**). We harnessed those strain-specific variants to map sequencing reads to their chromosome of origin. Overall, 9,016 protein-coding genes and noncoding RNAs had detectable ASE in at least one tissue-by-context analysis (∼37% of all expressed genes, **Supplemental Figure S2**). Next, we identified 2,853 genes with significant ASE biases (∼6%) and classified them into two patterns: parent-of-origin dependent (unequal expression based on the allele’s parental origin) and sequence dependent (unequal expression based on the allele’s nucleotide identity) (**Figure 1, Supplemental Table S1**).

### Parent-of-origin and sequence dependent ASE patterns are prevalent and distinct

Across our 27 tissue-by-context analyses, we identified 271 genes with significant parent-of-origin dependent ASE (**Supplemental Table S2 – S4**). HYP had the greatest number of significant genes (n = 229), followed by WAT (n = 35), then LIV (n = 27). 14 genes were expressed in multiple tissues, but the majority were tissue-specific (**Figure 2A**). In HYP, the most genes were detected in the “All” context. However, in WAT and LIV, more genes were only detected in a specific diet, sex, and/or diet-by-sex context; those expression biases were missed when contexts were collapsed (**Figure 2B**). 214 genes (79%) were paternally biased, 55 genes (20%) were maternally biased, and 2 genes (1%) switched their expression bias direction across the cohorts (**Figure 2C**). This heavy skew is driven by a 672 kb cluster of 171 paternally-biased small nucleolar RNAs (snoRNAs) located within the Prader-Willi/Angelman syndrome (**PWS/AS**) orthologous domain on mouse chromosome 7 that are only expressed in HYP (**Supplemental Figure S3**). Parent-of-origin dependent ASE genes also included protein-coding genes, microRNAs, long noncoding RNAs, long interspersed noncoding RNAs, and pseudogenes (**Figure 2D**).

**Figure 2:**
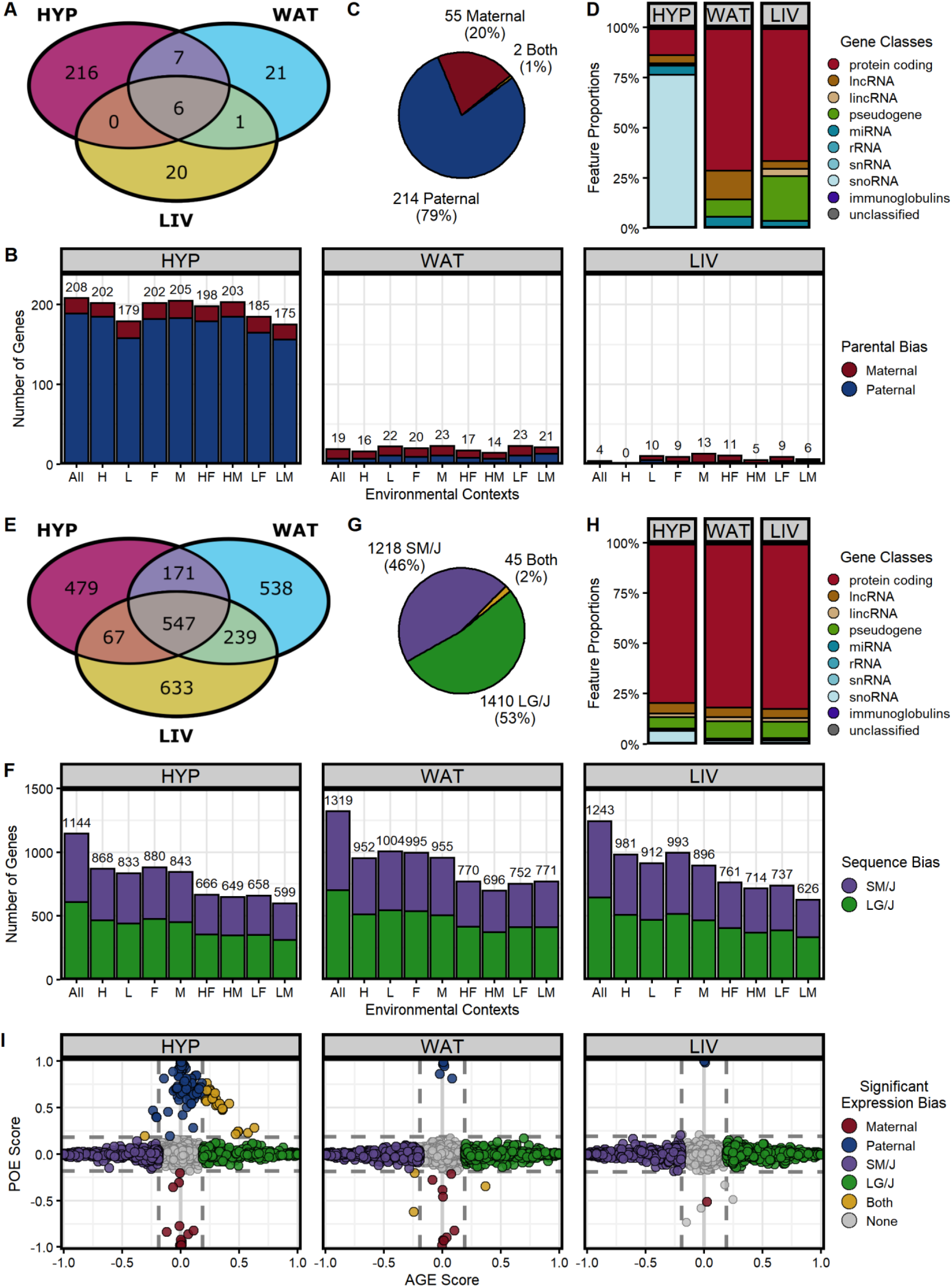
Parent-of-origin and sequence dependent allele-specific expression patterns are prevalent and distinct. **(A)** Venn diagram of the total parent-of-origin dependent ASE genes across HYP, LIV, and WAT (all contexts collapsed). **(B)** Number of genes with significant parental biases in each tissue-by-context analysis (maternal = red, paternal = blue): all contexts collapsed (All), high fat diet (H), low fat diet (L), females (F), males (M), high fat-fed females (HF), high fat-fed males (HM), low fat-fed females (LF), and low fat-fed males (LM). **(C)** Summary of expression bias directions across all analyses: paternally biased (blue), maternally biased (red), and genes that switch bias direction depending on the cohort (yellow). **(D)** Gene class proportions of significant parent-of-origin dependent ASE genes in each tissue. **(E)** Venn diagram of the total sequence dependent ASE genes across HYP, LIV, and WAT (all contexts collapsed). **(F)** Number of genes with significant sequence biases in each tissue-by-context analysis (SM/J = purple, LG/J = green). **(G)** Summary of expression bias directions across all analyses: SM/J biased (purple), LG/J biased (green), and genes that switch bias direction depending on the cohort (yellow). **(H)** Gene class proportions of significant sequence dependent ASE genes in each tissue. **(I)** Parent-of-Origin Effect (POE) versus Allelic Genotype Effect (AGE) scores in the “All” context of each tissue. Genes with significant parental biases (red = maternal, blue = paternal) have extreme POE scores, but weak AGE scores. Conversely, genes with significant sequence biases (purple = SM/J, green = LG/J) have extreme AGE scores, but weak POE scores. Some genes have both parental and sequence biases (yellow). Most genes have no bias (gray). Dashed lines indicate effect score thresholds.

We also identified 2,673 genes with significant sequence dependent ASE across our 27 tissue-by-context analyses (**Supplemental Table S2 – S4**). WAT had the greatest number of significant genes (n = 1,495), followed by LIV (n = 1,486), then HYP (n = 1,264). While some genes’ biases were tissue-specific, 1,657 genes (62%) were biased in multiple tissues (**Figure 2E**). In each tissue, the most genes were detected in the “All” context, then the diet- or sex-specific contexts, and finally the diet-by-sex-specific contexts, likely reflecting the sample sizes and power available in each cohort (**Figure 2F**). 1,218 genes (46%) were SM/J biased, 1,410 genes (53%) were LG/J biased, and 45 genes (2%) switched their expression bias direction across the cohorts (**Figure 2G**). Sequence dependent ASE genes were predominantly classified as protein-coding genes (∼80%) in each tissue, but also included pseudogenes, immunoglobulins, and various non-coding RNAs (long non-coding, long interspersed non-coding, micro, ribosomal, small interfering, and small nucleolar) (**Figure 2H**).

Parent-of-origin and sequence dependent ASE patterns were often mutually exclusive. In all three tissues, genes with extreme parental biases typically had weak sequence biases and vice versa (**Figure 2I, Supplemental Figure S4)**. However, 61 genes in HYP and 3 genes in WAT showed both parental and sequence biases, likely due to epigenetic regulatory mechanisms affected by haplotype variation/allelic identity. Both ASE patterns occurred genome-wide (**Supplemental Figure S5**). Parent-of-origin dependent ASE genes tended to cluster in well-known imprinted domains, such as the *Ube3a/Snrpn, Meg3, Peg3/Usp29*, and *H13/Mcts2* domains (**Supplemental Figures S6**). Sequence dependent ASE genes were spread more diffusely across chromosomes, but occasionally clustered in regions potentially controlled by the same regulatory element (**Supplemental Figures S7**).

### Parent-of-origin dependent ASE recapitulates canonical imprinting patterns

To evaluate whether parent-of-origin dependent ASE is influenced by tissue or environmental context (diet and/or sex), we characterized the expression profiles of the 271 parentally biased genes across our 27 tissue-by-context analyses (3 tissues x 9 diet-by-sex contexts). For each analysis, we quantified the direction and magnitude of each gene’s parental expression bias by calculating a Parent-of-Origin Effect (**POE**) score from the mean allelic bias of each F_1_ reciprocal cross. POE scores range from -1 (completely maternally expressed) to +1 (completely paternally expressed); a score of 0 indicates biallelic expression (see Methods). In each tissue-by-context analysis, a gene could be expressed in one of three ways: significant parental bias, biallelic (expressed but with no allele-specific bias), or not expressed. We sorted the 271 genes with significant parent-of-origin dependent ASE into three expression profiles: tissue-independent, tissue-dependent, and context-dependent (**Figure 3A-3D, Supplemental Figure S3**).

**Figure 3:**
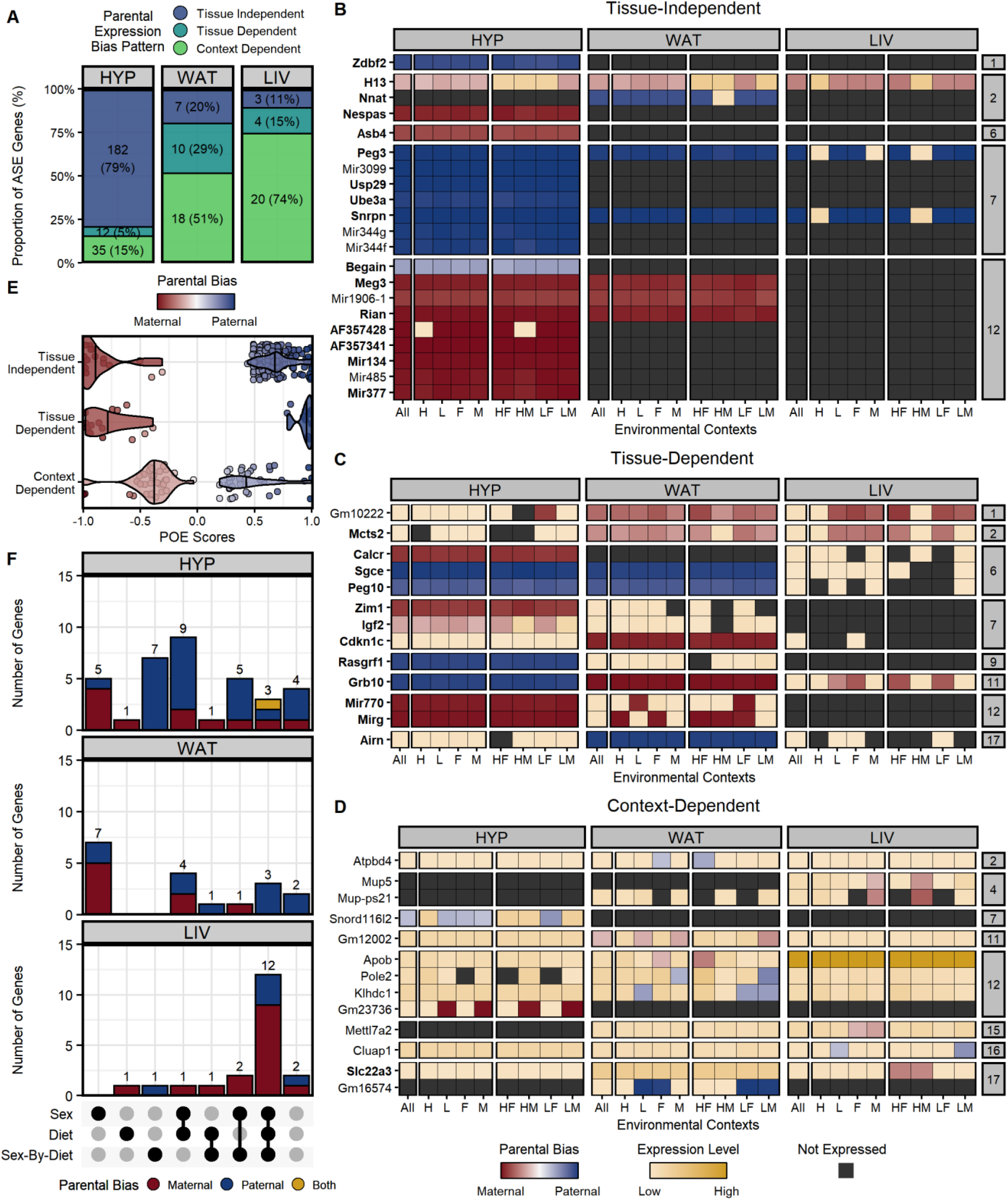
Atlas of parent-of-origin dependent ASE patterns across tissues and environmental contexts. **(A)** Number and proportion of ASE genes per tissue with each parental expression bias profile: tissue-independent (dark blue), tissue-dependent (teal), and context-dependent (green). Heatmaps of ASE profiles across tissues and contexts: **(B)** tissue-independent (parental bias wherever expressed), **(C)** tissue-dependent (parental bias in some tissues, no bias in others), and **(D)** context-dependent (parental bias only in certain diet-by-sex contexts). A subset of the 271 genes with parent-of-origin dependent ASE is shown, including those validated with pyrosequencing (see **Supplemental Figure S3** for full heatmap). Genes are color-coded by their expression pattern in each tissue-by-context analysis. Shades of red and blue indicate their degree of maternal or paternal bias, respectively (POE scores). Where genes are not significantly biased, shades of yellow indicate their biallelic gene expression levels (log-transformed total counts). Black indicates genes are not expressed in that context/tissue. Bolded genes are canonically imprinted. The y-axis is grouped and sorted by chromosomal position. Each supercolumn denotes a tissue: hypothalamus (HYP), white adipose (WAT), and liver (LIV). Each subcolumn denotes an environmental context: all contexts collapsed (All), high fat diet (H), low fat diet (L), females (F), males (M), high fat-fed females (HF), high fat-fed males (HM), low fat-fed females (LF), and low fat-fed males (LM). **(E)** Parent-of-Origin Effect (POE) score distributions for each parental expression bias profile. Vertical lines indicate that profile’s mean POE score. Dots represent individual ASE genes. **(F)** UpSet plots for each tissue summarizing the set intersections of context-dependent genes with significant sex, diet, and/or sex-by-diet effects. Bar height and color indicate the number of genes with each parental bias direction: maternal biased (red), paternal biased (blue), and those that switch expression bias direction depending on the cohort (yellow).

We identified 183 tissue-independent genes (67%), defined as a consistent parental bias in every tissue they were expressed. Within a given tissue, they were parentally biased in most environmental contexts (**Figure 3B**). Next, we identified 15 genes (6%) as tissue-dependent. These genes showed a parental bias across most contexts in one or two tissues, but were biallelically expressed (no bias) across contexts in the other tissue(s) (**Figure 3C**). Here, we distinguish between tissue-specific gene expression (not expressed in certain tissues) and tissue-dependent ASE (biased expression only in certain tissues).

All 198 tissue-independent/dependent genes were related to genomic imprinting, the best characterized mechanism of parent-of-origin dependent ASE. 29 genes were canonically imprinted (bolded in **Figure 3**, defined in Methods) and the other 169 genes were various non-coding RNAs located within well-known imprinted domains. For example, 156 genes were part of a cluster of 171 paternally-biased snoRNAs in the PWS/AS domain on mouse chromosome 7 (**Supplemental Figure S3**). As expected with imprinting, these genes were extremely biased towards one parent’s allele; their mean POE scores were -0.82 and 0.71 for maternally and paternally biased genes, respectively (**Figure 3E**). HYP had the highest proportion of these tissue-independent/dependent genes among our three adult tissues (85%), even if the HYP-specific snoRNA cluster in the PWS/AS domain is excluded (52%). WAT had the second highest proportion (49%), but very few of these genes were expressed in LIV (26%) (**Figure 3A**). These findings are consistent with the previously-reported dichotomy of imprinting levels between neural and non-neural adult tissues (Babak et al. 2015) as well as imprinting’s role in development and cognition (Barlow and Bartolomei 2014). Furthermore, our data replicated 25 of the 96 genes previously found to be imprinted in these tissues (Babak et al. 2015), comprising nearly all bolded genes in **Figures 3B - 3D**. We could not compare the imprinting status of 55 genes because they could not be probed for ASE in this F_1_ model (34) or did not appear in the mm10 reference assembly (21). 5 genes had significant sequence biases instead and the remaining 11 were biallelically expressed in our datasets, potentially due to methodology differences between the studies.

We validated the expression profiles of two canonically imprinted genes (*Peg3* and *Grb10*) by pyrosequencing (**Figure 3B-3C, Supplemental Figure S8**). *Peg3* (paternally expressed gene 3) had a significant paternal bias in all three tissues, regardless of context. Its locus has 60 variants between the LG/J and SM/J backgrounds, though none are predicted to be functional. *Peg3* functions as a DNA-binding transcriptional repressor to control fetal growth rates, maternal caring behaviors, and tumor growth. It is only expressed from the paternal allele in most tissues, especially the placenta and brain (Thiaville et al. 2013; He and Kim 2014). *Grb10* (growth factor receptor bound protein 10) had a tissue-dependent pattern of significant paternal bias in HYP but a maternal bias in WAT and LIV. Its locus has 154 variants between the LG/J and SM/J backgrounds, but none are predicted to be functional. *Grb10* encodes an adapter protein that interacts with receptor tyrosine kinases to impact insulin signaling and growth hormone pathways (He et al. 1998). It has a documented pattern of maternal expression in most adult mouse tissues, but paternal expression in the brain (Plasschaert and Bartolomei 2015).

### Dietary environment and sex influence parent-of-origin dependent ASE in a partially imprinted manner

Finally, we classified 73 genes (27%) as context-dependent, meaning they had a parental bias only in certain environmental contexts within a tissue, but biallelic expression (no bias) in other contexts and/or tissues (**Figure 3D**). Only three of these genes were canonically imprinted and 18 genes were non-coding RNAs located in known imprinted domains (including 14 snoRNAs in the PWS/AS domain). The remaining 52 genes had no clear connection to genomic imprinting, yet they showed significant parent-of-origin dependent ASE in certain context(s). These context-dependent genes had more subtle allelic biases than the tissue-independent/dependent genes; their mean POE scores were -0.39 and 0.39 for maternally and paternally biased genes, respectively (**Figure 3E**). These patterns are consistent with partial imprinting, where the two parental alleles are differentially expressed in a less extreme manner than the uniparental expression associated with genomic imprinting (Wolf et al. 2008; Morcos et al. 2011). Interestingly, LIV had the highest proportion of context-dependent genes among its total parent-of-origin dependent ASE genes (74%) while HYP had the lowest proportion (15%) (**Figure 3A**). LIV is also a tissue where more parent-of-origin dependent ASE genes were detected in the diet- and/or sex-specific contexts than the “All” context (**Figure 2B**). LIV is incredibly responsive to environmental factors (especially diet), given its central roles in digestion and detoxification (Trefts et al. 2017). Taken together, these findings suggest a mechanism of parent-of-origin dependent ASE outside of traditional imprinting that is sensitive to environmental perturbations.

To further explore how intrinsic (sex) and/or extrinsic (dietary fat) environments can alter parental ASE biases, we calculated individualized POE scores for each context-dependent gene and modeled how they vary across diet, sex, and diet-by-sex contexts (**Supplemental Tables S5 and S6**). These categories were not mutually exclusive; across all three tissues, most context-dependent genes were significant for more than one effect. Nonetheless, when we intersected these significant gene lists, we found that each tissue showed a distinct pattern of context-dependent parental ASE biases (**Figure 3F, Supplemental Figure S9**). For example, LIV had similar proportions of significant sex, diet, and diet-by-sex effects (each 75 – 80% of its context-dependent genes); 60% of its genes were significant for all three effects. Most of WAT’s genes had a significant sex effect (83%), but diet or diet-by-sex effects were much less common (44% and 28%, respectively). Finally, 63% of HYP’s genes had a significant sex effect, while 40 – 45% had significant diet or diet-by-sex effects.

We validated the expression profiles of two context-dependent genes (*Apob* and *Slc22a3*) by pyrosequencing (**Figure 3D, Supplemental Figure S10**). *Apob* (apolipoprotein B) had a significant diet effect in WAT, reflected in significant maternal biases in the HF and F contexts. It also had a maternal bias in the HM context, but low sample sizes excluded it from further analysis. *Apob* was biallelically expressed in the remaining contexts in WAT and all contexts in LIV and HYP. Its locus has 126 variants between the LG/J and SM/J backgrounds, including seven non-synonymous SNPs in the coding region (six = LG/J genome, one = SM/J genome). *Apob* produces the main component of lipoproteins, which transport lipids (including cholesterol) in the blood (Olofsson and Borèn 2005). Maternal-specific associations between *APOB* variants and adiposity traits have been found in humans (Hochner et al. 2015). *Apob* expression levels also differ between high and low fructose diets in mice livers, suggesting its function is susceptible to dietary environment (Sud et al. 2017). *Slc22a3* (solute carrier family 22 member 3) had significant diet and diet-by-sex effects in LIV, reflected in strong maternal biases in the HF and HM contexts. *Slc22a3* was biallelically expressed in the remaining contexts in LIV and all contexts in WAT and HYP. Its locus has 285 variants between the LG/J and SM/J backgrounds, but none are predicted to be functional. *Slc22a3* is a transporter that eliminates organic cations from cells, such as monoamine neurotransmitters, cationic drugs, and xenobiotics (Kekuda et al. 1998). It has been reported to be maternally expressed in liver and extraembryonic tissues, but biallelically expressed elsewhere (Babak et al. 2015). *Slc22a3* is also significantly differentially expressed in kidneys between high fat diet- and chow-fed mice (Gai et al. 2016).

### Sequence dependent ASE arises from haplotype-specific genetic variation

Next, we similarly characterized the expression profiles of the 2,673 sequence biased genes across our 27 tissue-by-context analyses (3 tissues x 9 diet-by-sex contexts) to evaluate whether sequence dependent ASE is also influenced by tissue or environmental context. For each analysis, we quantified the direction and magnitude of each gene’s sequence expression bias by calculating an Allelic Genotype Effect (**AGE**) score from the mean allelic bias of each F_1_ reciprocal cross. AGE scores range from -1 (completely SM/J expressed) to +1 (completely LG/J expressed); a score of 0 indicates biallelic expression (see Methods). In each analysis, a gene could be expressed in one of three ways: significant sequence/allelic genotype bias, biallelic (expressed but with no allele-specific bias), or not expressed. We sorted the 2,673 genes with significant sequence dependent ASE into three expression profiles: tissue-independent, tissue-dependent, and context-dependent (**Figure 4A-4D**).

**Figure 4:**
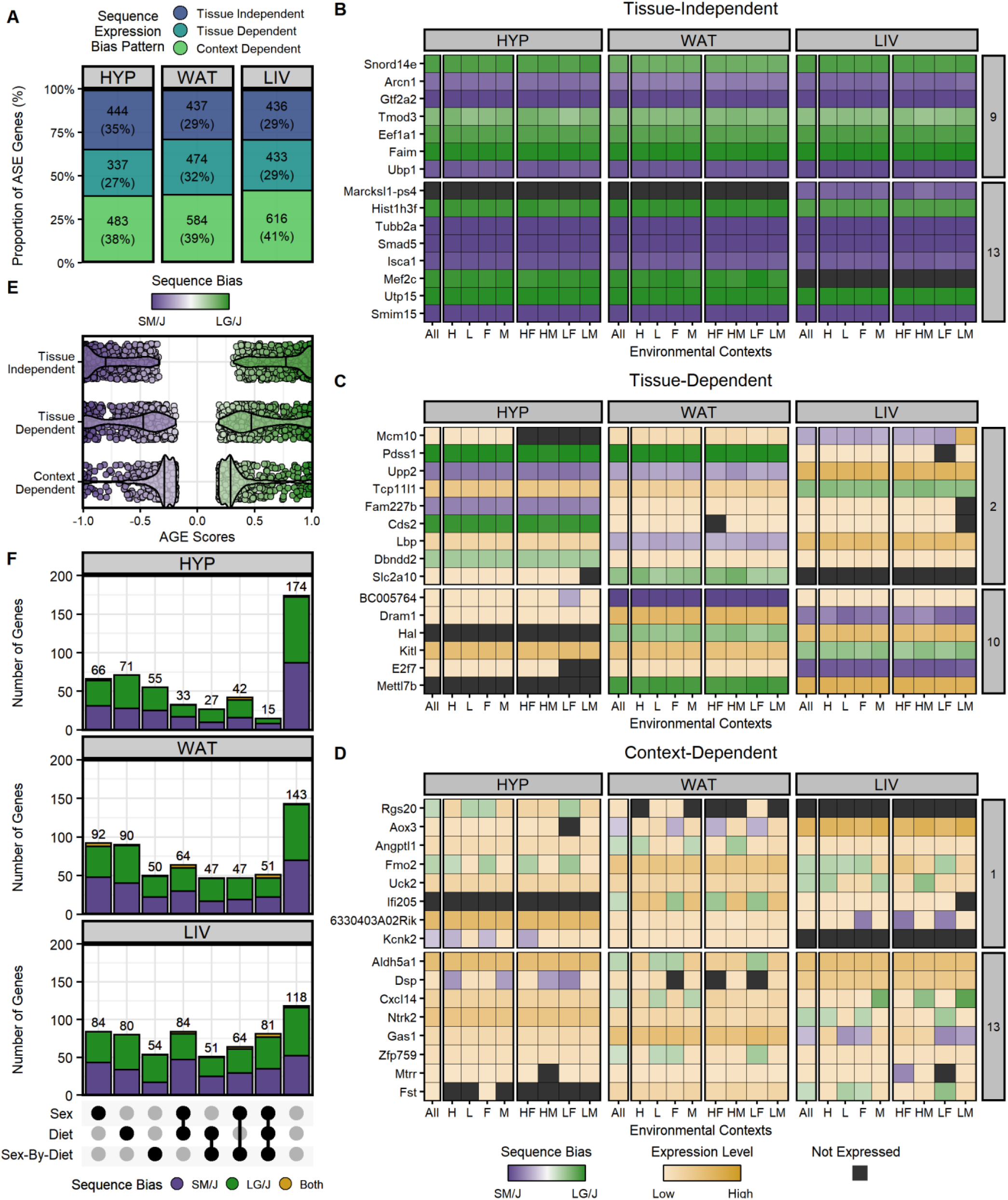
Atlas of sequence dependent ASE patterns across tissues and environmental contexts. **(A)** Number and proportion of ASE genes per tissue with each sequence expression bias profile: tissue-independent (dark blue), tissue-dependent (teal), and context-dependent (green). Heatmaps of ASE profiles across tissues and contexts: **(B)** tissue-independent (sequence bias wherever expressed), **(C)** tissue-dependent (sequence bias in some tissues, no bias in others), and **(D)** context-dependent (sequence bias only in certain diet-by-sex contexts). A subset of the 2,673 genes with significant sequence dependent ASE are shown, including those validated with pyrosequencing. Genes are color-coded by their expression pattern in each tissue-by-context analysis. Shades of purple and green indicate their degree of SM/J or LG/J allelic bias, respectively (AGE scores). Where genes are not significantly biased, shades of yellow indicate their biallelic gene expression levels (log-transformed total counts). Black indicates genes are not expressed in that context/tissue. The y-axis is grouped and sorted by chromosomal position. Each supercolumn denotes a tissue: hypothalamus (HYP), white adipose (WAT), and liver (LIV). Each subcolumn denotes an environmental context: all contexts collapsed (All), high fat diet (H), low fat diet (L), females (F), males (M), high fat-fed females (HF), high fat-fed males (HM), low fat-fed females (LF), and low fat-fed males (LM). **(E)** Allelic Genotype Effect (AGE) score distributions for each sequence expression bias profile. Vertical lines indicate that profile’s mean AGE score. Dots represent individual ASE genes. **(F)** UpSet plots for each tissue summarizing the set intersections of context-dependent genes with significant sex, diet, and/or sex-by-diet effects. Bar height and color indicate the number of genes with each sequence bias direction: SM/J biased (purple), LG/J biased (green), and those that switch expression bias direction depending on the cohort (yellow).

We identified 605 genes (23%) as tissue-independent, meaning they had a consistent sequence bias in every tissue they were expressed. Within a given tissue, they had a sequence bias in most environmental contexts (**Figure 4B**). These tissue-independent genes were strongly biased towards one strain’s allele; their mean AGE scores were -0.79 and 0.78 for SM/J and LG/J biased genes, respectively (**Figure 4E**). They also comprised a similar proportion of the total sequence dependent ASE genes in all three tissues (29-35%, **Figure 4A**). These patterns likely reflect the vast genetic variation that has accumulated between the LG/J and SM/J backgrounds over the decades, whereby a *cis*-acting variant impacts one strain’s allelic function wherever that gene is expressed, regardless of tissue type.

We validated the expression profiles of two tissue-independent genes (*Eef1a1* and *Tubb2a*) by pyrosequencing (**Figure 4B, Supplemental Figure S11**). *Eef1a1* (eukaryotic translation elongation factor 1 alpha 1) had a significant LG/J allelic bias in all three tissues, regardless of context. Its locus has 30 variants between the LG/J and SM/J backgrounds, including two non-synonymous SNPs in the coding region of the SM/J genome. *Eef1a1* delivers aminoacylated transfer RNAs to the elongating ribosome during protein synthesis and has crucial roles in protein degradation, RNA virus replication, and other cellular processes. It is abundantly and ubiquitously expressed in most tissues (Mateyak and Kinzy 2010; Li et al. 2013). *Tubb2a* (tubulin beta-2A Class IIa) had a significant SM/J allelic bias in all three tissues, regardless of context. Mutations in this gene are rare and often non-viable; however, its locus has one SNP in the LG/J genome. *Tubb2a* is a tubulin isoform that binds GTP to create microtubules, which are critical for the cytoskeleton organization, intracellular trafficking, and mitotic cell division. It is also ubiquitously expressed in most tissues, especially the brain (Hammond et al. 2008; Rice et al. 2008).

### Sequence dependent ASE can be mediated by tissue-specific features

Next, we identified 684 genes (25%) as tissue-dependent. These genes showed a sequence bias across most contexts in one or two tissues, but were biallelically expressed (no bias) across contexts in the other tissue(s) (**Figure 4C**). Here, we again distinguish between tissue-dependent ASE (biallelic expression in some tissues) and tissue-specific gene expression (simply not expressed in some tissues). These tissue-dependent genes were moderately biased towards one strain’s allele; their mean AGE scores were -0.57 and 0.55 for SM/J and LG/J biased genes, respectively (**Figure 4E**). They also comprised a similar proportion of the total sequence dependent ASE genes in all three tissues (27-32%, **Figure 4A**). These patterns demonstrate that sequence dependent ASE is not solely due to genetic variation; here, tissue-specific epigenetic factors likely interact with *cis*-acting variants to influence allelic frequency.

We validated the expression profiles of two tissue-dependent genes (*Upp2* and *Mettl7b*) by pyrosequencing (**Figure 4C, Supplemental Figure S12**). *Upp2* (uridine phosphorylase 2) had a strong SM/J allelic bias across contexts in HYP, a weaker SM/J bias in WAT, and biallelic expression in LIV. Its locus has 1,043 variants between the LG/J and SM/J backgrounds, including five non-synonymous SNPs in the coding region of the LG/J genome. *Upp2* catalyzes the phosphorolysis of uridine into uracil and ribose-1-phosphate, which are used as carbon and energy sources during catabolic metabolism. In mice, it is predominantly expressed in liver and weakly expressed in brain (Johansson 2003; Roosild et al. 2011). *Mettl7b* (methyltransferase-like 7b) had a LG/J allelic bias across contexts in WAT, biallelic expression in LIV, and was not expressed in HYP. Its locus has 77 variants between the LG/J and SM/J backgrounds, including one non-synonymous SNP in the coding region of the SM/J genome. *Mettl7b* is an akyl thiol methyltransferase that is implicated in several cancers. It is highly expressed in lipid droplets, particularly in liver and adipose tissue (Maldonato et al. 2021; Turró et al. 2006).

### Sequence dependent ASE is sensitive to sex and dietary environments

Finally, we classified 1,384 genes (52%) as context-dependent. These genes had a sequence bias only in certain environmental contexts within a tissue, but biallelic expression (no bias) in other contexts and/or tissues (**Figure 4D**). These context-dependent genes had more subtle allelic biases than the other two profiles; their mean AGE scores were -0.32 and 0.34 for SM/J and LG/J biased genes, respectively (**Figure 4E**). LIV had the most context-dependent genes (n = 616) and HYP had the fewest (n = 483), but overall they comprised a similar proportion of the total ASE genes in all three tissues (38-41%, **Figure 4A**). These patterns suggest that environmental factors can interact with genetic variation to influence the final allelic composition of a gene’s product, resulting in sequence dependent ASE.

To further explore how sex and/or dietary fat can alter sequence/allelic genotype ASE biases, we calculated individualized AGE scores for each context-dependent gene and modeled how they vary across diet, sex, and diet-by-sex contexts (**Supplemental Tables S5 and S6**). When we intersected these gene lists, we found that each tissue showed a similar pattern of context-dependent sequence ASE biases (**Figure 4F, Supplemental Figure S13**). Significant sex effects were the most prevalent in each tissue, ranging from 156 genes in HYP (32%), to 254 genes in WAT (43%), and 313 genes in LIV (51%). Diet effects were slightly less common but still widespread: 146 genes in HYP (30%), 252 genes in WAT (43%), and 296 genes in LIV (48%). Finally, diet-by-sex effects were also pervasive in each tissue, comprising 139 genes in HYP (29%), 195 genes in WAT (33%), and 250 genes in LIV (41%). These categories were not mutually exclusive; most context-dependent genes were significant for more than one effect but in different combinations across the three tissues.

We validated the expression profiles of two context-dependent genes (*Ifi205* and *Gas1*) by pyrosequencing (**Figure 4D, Supplemental Figure S14**). *Ifi205* (interferon activated gene 205) had significant sex and diet-by-sex effects in WAT, reflected in strong LG/J allelic biases in the three female-related (HF, LF, F) and “All” contexts. *Ifi205* was biallelically expressed in the remaining contexts in WAT and all contexts in LIV. Its locus has 254 variants between the LG/J and SM/J backgrounds, including ten non-synonymous SNPs in the coding region of the SM/J genome. *Ifi205* binds DNA in response to interferon signaling, a cytokine family that activates the immune system. Significant sex differences have been reported in mouse *Ifi205* expression levels, where females have higher total expression levels than males (Albrecht et al. 2005; Cao et al. 2018). *Gas1* (growth arrest specific 1) had significant diet and sex effects in LIV, reflected in strong SM/J biases in the three low fat diet-related (LF, LM, L), female, and “All” contexts. *Gas1* was biallelically expressed in the remaining high fat diet and male contexts in LIV and all contexts in HYP and WAT. Its locus has 57 variants between the LG/J and SM/J backgrounds, though none are predicted to have functional impacts. *Gas1* encodes a membrane glycoprotein that binds and regulates sonic hedgehog during development. *Gas1* has been found to be differentially expressed between high and low selenium diets in mice ovaries, suggesting its expression may be sensitive to dietary environment (Lee et al. 2001; Qazi et al. 2021).

### Sequence dependent ASE genes can switch the direction of their allelic biases

Surprisingly, we found 45 sequence dependent ASE genes with inconsistent patterns of allelic biases. These genes showed significant ASE in opposite directions among the tissues and/or environmental contexts (**Figure 5**). For 44 of the 45 genes (98%), such direction-switching occurred at the tissue level: for example, a gene may have a LG/J bias in one tissue, a SM/J bias in another tissue, and sometimes even no bias (biallelic) in the third tissue. Four genes had tissue-independent ASE, or a sequence bias across contexts in every expressed tissue (albeit in different directions). 19 genes had context-dependent ASE, or a sequence bias only in certain diet and/or sex contexts that switched direction across tissues. One of these genes (*Cidec*) had a diet- and sex-dependent switch in ASE direction within the same tissue, discussed further below. The remaining 22 genes had a combination of context- and tissue-dependent ASE patterns: such genes had a sequence bias in one direction across most contexts in one or two tissues, but a context-dependent sequence bias in the opposite direction in another tissue. Overall, these genes had moderate allelic biases, reflecting the various expression profiles herein; their mean AGE scores were -0.38 and 0.40 for SM/J and LG/J biased genes. These dynamic direction-switching patterns confirm that sequence dependent ASE is not solely due to genetic variation. Otherwise, the same variant causing biased expression in one tissue would also be present in the cells of any other tissue expressing that gene. These patterns hint at epigenetic regulatory elements or post-transcriptional modifications that interact with genetic variation in a tissue-specific manner to influence the final allelic frequency.

**Figure 5:**
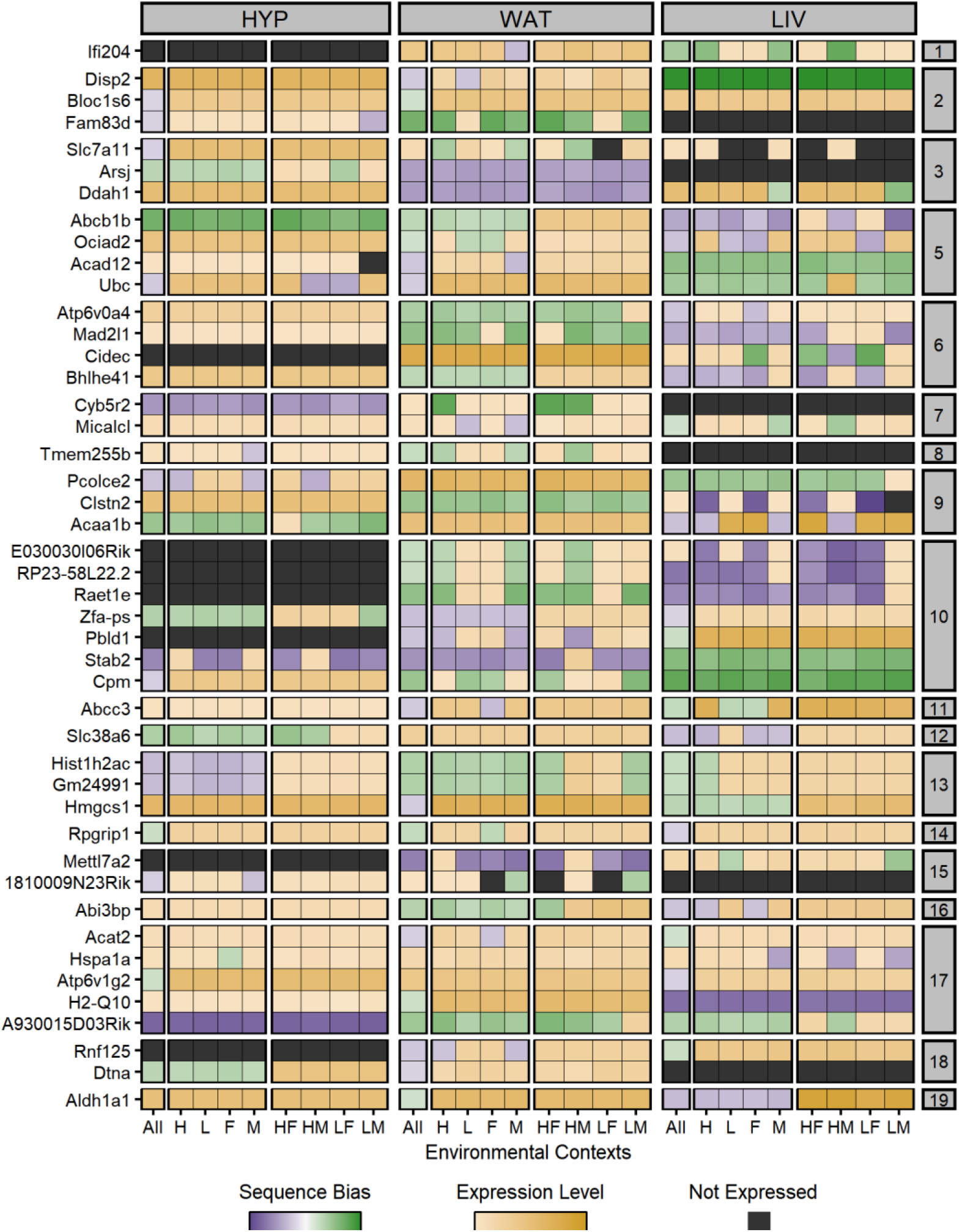
45 sequence dependent ASE genes switch their expression bias directions across tissues or environmental contexts. Heatmap of ASE profiles for the 45 genes with significant sequence biases in opposite directions across tissues and/or environmental contexts, including those validated with pyrosequencing. Genes are color-coded by their expression pattern in each tissue-by-context analysis. Shades of purple and green indicate their degree of SM/J or LG/J allelic bias, respectively (AGE scores). Where genes are not significantly biased, shades of yellow indicate their biallelic gene expression levels (log-transformed total counts). Black indicates genes are not expressed in that context/tissue. The y-axis is grouped and sorted by chromosomal position. Each supercolumn denotes a tissue: hypothalamus (HYP), white adipose (WAT), and liver (LIV). Each subcolumn denotes an environmental context: all contexts collapsed (All), high fat diet (H), low fat diet (L), females (F), males (M), high fat-fed females (HF), high fat-fed males (HM), low fat-fed females (LF), and low fat-fed males (LM).

We validated the expression profiles of two direction-switching genes (*Stab2* and *Ociad2*) with pyrosequencing (**Figure 5, Supplemental Figure S15**). *Stab2* (stabilin 2) had a significant SM/J bias across most contexts in HYP and WAT, but a LG/J bias in all contexts in LIV. Its locus has 1,019 variants between the LG/J and SM/J backgrounds, including 14 non-synonymous SNPs in the coding region of the SM/J genome. *Stab2* binds to hyaluronic acid and mediates its transportation inside the cell (Zhou et al. 2002). It is predominantly expressed in two isoforms in the liver and spleen, while weakly expressed in the brain and adipose tissue (Falkowski et al. 2003). Together with the non-synonymous variants in the SM/J allele, this may explain *Stab2*’s LG/J bias in the LIV but SM/J bias in the other two tissues. *Ociad2* (ovarian cancer immunoreactive antigen domain-containing protein 2) had significant sex and sex-by-diet effects in WAT and LIV. These are reflected in strong LG/J biases in the L, F, and “All” contexts in WAT, but strong SM/J biases in the LF, L, F, and “All” contexts in LIV. *Ociad2* was biallelically expressed in the remaining contexts of both tissues and in HYP. Its locus has 118 variants between the LG/J and SM/J backgrounds, though none are predicted to be functional. *Ociad2* activates STAT3, regulates cell migration, and is implicated in several cancers. It is moderately expressed in several non-cancerous tissues, including the brain and liver (Sinha et al. 2018). Sex or diet differences in expression levels have not been explored in the literature, but *Ociad2* is highly expressed in female ovarian tumors (Nagata et al. 2012). Low fat-fed female WAT and LIV tissues have strong sequence biases in *Ociad2* (but in different directions), suggesting a tissue- and sex-specific expression pattern that could have functional consequences.

### ASE genes are enriched in QTLs for metabolic and anatomical traits

Mapping studies in the LG/J x SM/J advanced intercross line (AIL) population have identified hundreds of QTLs associated with a constellation of complex traits. Such approaches are agnostic to tissue type and often implicate regions with several genes, making it difficult to pinpoint which genes are phenotypically relevant. In contrast, ASE patterns are gene-specific and vary across tissues and environments. If there is a functional difference between two alleles, then an expression imbalance in the right conditions can have phenotypic consequences. We hypothesized that significant ASE genes are over-represented in QTLs and thus could be prioritized as the candidate gene(s) manifesting that signal. To assess this, we integrated ASE and QTL data from populations at various degrees of intercrossing the same founder strains (**Figure 6A, Supplemental Table S7**).

**Figure 6:**
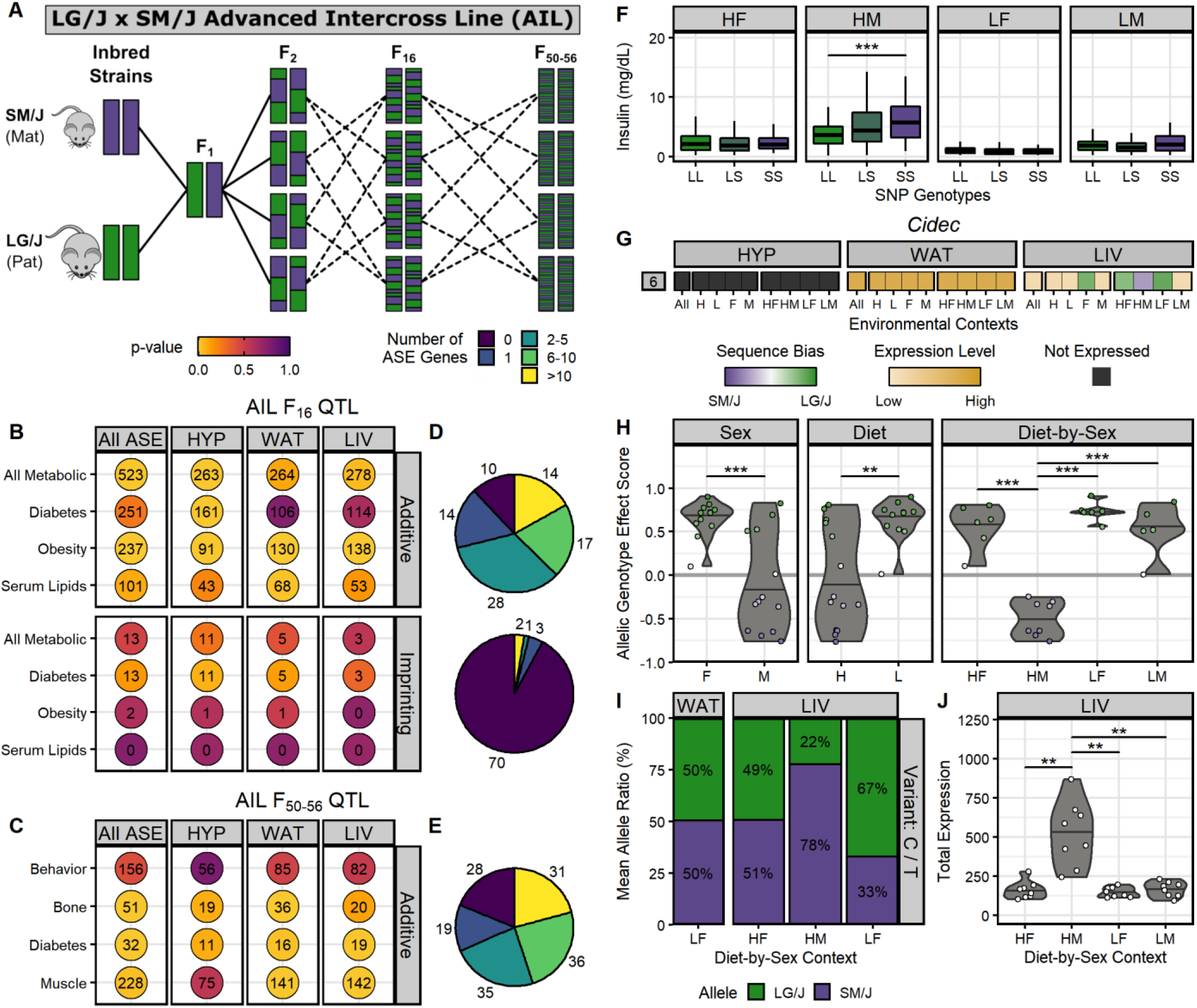
Incorporating ASE patterns with QTL mapping data can help prioritize candidate genes. **(A)** Breeding scheme for the LG/J x SM/J Advanced Intercross Line, produced by intercrossing mice beyond the F_2_ generation. We calculated enrichment of tissue-specific ASE gene sets (x-axis) in trait category-specific QTL sets (y-axis) among: **(B)** sequence dependent ASE genes in additive AIL F_16_ QTL (top) and parent-of-origin dependent ASE genes in imprinting AIL F_16_ QTL (bottom), and **(C)** sequence dependent ASE genes in additive AIL F_50-56_ QTL. Dot color corresponds to the enrichment p-value. Numbers indicate the total overlapping ASE genes in each QTL set. **(D & E)** Summary of how many AIL QTLs contain each range of ASE genes: 0 (purple) to more than 10 (yellow). Pie charts match their adjacent ASE gene and QTL set intersections. **(F)** The AIL F_16_ QTL *Ddiab6d* showed context-dependent additive effects. Boxplots of serum insulin levels across SNP rs6393943 genotypes in F_16_ mice (LL = LG/J homozygous, LS = heterozygous for LG/J and SM/J alleles, SS = SM/J homozygous). The x-axis is grouped by diet-by-sex context (HF = high fat-fed females, HM = high fat-fed males, LF = low fat-fed females, LM = low fat-fed males). Horizontal bars denote the mean phenotype of each cohort. *** p ≤ 0.001; assessed by Student’s t-test. **(G)** *Cidec* is located within the *Ddiab6d* locus and had a diet- and sex-dependent switch in sequence bias direction within LIV. Expression profile heatmap of *Cidec* across tissues and environmental contexts (see Figures 4 and 5 for full description). **(H)** *Cidec* ASE biases had significant sex, diet, and diet-by-sex effects. Violin plots display individual AGE scores for *Cidec* (y-axis) across environmental contexts (x-axis). Horizontal bars denote the mean AGE scores. Individual points are color-coded by their sequence bias. ** p ≤ 0.01, *** p ≤ 0.001; assessed by ANOVA (sex, diet) or Tukey’s post-hoc tests (diet-by-sex). **(I)** *Cidec* expression profile in LIV and WAT was validated by pyrosequencing. Bar graphs denote the mean allelic ratios (y-axis) in select tissue-by-context cohorts (x-axis) and are color-coded by allele (LG/J = green, SM/J = purple). **(J)** *Cidec* had significantly higher expression in the HM context. Violin plots of total expression levels in LIV (y-axis) for each diet-by-sex context (x-axis). Horizontal bars denote the mean expression. ** p ≤ 0.01; assessed by Student’s t-test.

Previously, 127 QTLs influencing obesity, diabetes, and serum lipid traits were mapped in the AIL F_16_ generation (Cheverud et al. 2011; Lawson et al. 2010, 2011b). These studies incorporated the same environmental contexts (sex and dietary fat) used here, traced the parental origin of each marker allele, and characterized the context-dependent additive and imprinting effects that influence metabolic traits. Overall, sequence dependent ASE genes were significantly enriched in the 83 F_16_ QTLs showing additive effects (p = 0.01, **Figure 6B**). We compared the QTL sets for each metabolic trait category with the full ASE gene sets for each tissue. QTLs for obesity-related traits were enriched for HYP, WAT, and LIV genes (all p < 0.01). QTLs for diabetes-related traits were enriched for HYP genes (p = 0.01), but not WAT or LIV genes (p = 0.76 and p = 0.62, respectively). QTLs for serum lipid-related traits were enriched for WAT genes (p = 0.02), but not HYP or LIV (p = 0.21 and p = 0.16, respectively) (**Figure 6B, Supplemental Figure S16**). These findings reflect the discrete roles of each tissue in metabolism physiology and illustrate that tissue-specific ASE patterns can inform which tissues are phenotypically relevant. Conversely, parent-of-origin dependent ASE genes (excluding the 171 snoRNA cluster in the PWS/AS domain) were not enriched in the 76 F_16_ QTLs showing imprinting effects (p = 0.47, **Figure 6B**). Tissue-specific ASE gene sets were also not enriched in the QTL sets for any metabolic trait category (p > 0.05, **Supplemental Figure S16**), supporting previous work that shows parental expression biases are often not the direct source of imprinting QTL effects (Macias-Velasco et al. 2021).

Additionally, 149 QTLs influencing behavioral, musculoskeletal, and metabolic traits were mapped in later generations (F_50-56_) of the same LG/J x SM/J AIL population (Gonzales et al. 2018; Cordero et al. 2019). These studies did not capture environmental contexts or parental origin in their analyses, but still explored how additive genetic effects impact complex traits in a context-independent manner. Overall, sequence dependent ASE genes were significantly enriched in additive F_50-56_ QTLs for muscle weights (p = 0.03), bone sizes (p = 0.01), and diabetes-related traits (p = 0.01, **Figure 6C**). These enrichment results are consistent with the history of the LG/J and SM/J founder strains: they were independently selectively bred for large and small body sizes, so it is not surprising that strain-specific genetic variation can induce ASE biases in this F_1_ model that often occur in/near regions relevant for morphometric traits. Interestingly, sequence dependent ASE genes were not enriched in additive F_50-56_ QTLs for behavioral traits related to locomotor activity and prepulse inhibition (p = 0.42, **Figure 6C**). Tissue-specific ASE gene sets were also not enriched in behavioral QTLs, even for the brain region HYP (p = 0.79, **Supplemental Figure S17**), suggesting the metabolic tissues profiled here for ASE are not relevant for these behavioral phenotypes.

Although sequence dependent ASE genes are located within additive AIL QTL more often than expected, individual QTLs typically contained few ASE genes. For example, the 83 additive AIL F_16_ QTLs overlap a mean of 75 genes each, yet 51% of those QTLs (n = 42) contained only 1-5 sequence dependent ASE genes. 37% of those QTLs (n = 31) contained more than 6 ASE genes while merely 12% (n = 10) had no ASE genes (**Figure 6D**). Similarly, the 149 additive AIL F_50-56_ QTLs overlap a mean of 61 genes each, but 36% of those QTLs (n = 54) contained only 1-5 sequence dependent ASE genes. 45% of those QTLs (n = 67) contained more than 6 ASE genes and 19% (n = 28) had no ASE genes (**Figure 6E**). In contrast, the 76 imprinting AIL F_16_ QTLs overlap a mean of 62 genes each, but 92% of them (n = 70) do not contain any parent-of-origin dependent ASE genes (**Figure 6D**). Only two imprinting F_16_ QTLs contain more than 10 ASE genes and they both overlap the 171 snoRNA cluster in the PWS/AS domain on chromosome 7.

### Cidec has a context-dependent switch in ASE bias direction and is a candidate gene for insulin levels

Since genes with significant ASE often comprise a small portion of all genes within a QTL, we propose that incorporating ASE patterns can help dissect QTL underpinnings and prioritize candidate genes in relevant tissues for functional validation. Here, we present an example of the utility of this orthogonal approach. *Ddiab6d* was an AIL F_16_ QTL on chromosome 6 associated with serum insulin, basal glucose, and glucose tolerance levels that showed additive genetic effects in a context-dependent manner (Lawson et al. 2011b). In the high fat-fed male context, AIL F_16_ mice that were homozygous for the SM/J allele had significantly higher insulin levels than the homozygous LG/J mice (p < 0.001). There were no significant differences in insulin levels between homozygote genotypes in the high fat-fed females, low fat-fed females, or low fat-fed male cohorts (**Figure 6F**). The *Ddiab6d* locus spanned 14.7Mb and contained 100 genes, complicating efforts to pinpoint the causal mechanisms of this additive effect.

However, only four sequence dependent ASE genes were located within the *Ddiab6d* locus: *Brpf1, Cidec, Irak2*, and *Syn2. Cidec* (cell death-inducing DNA fragmentation effector C) had a diet- and sex-dependent switch in ASE bias direction within the same tissue, which was a notable exception to the tissue-level ASE direction switches reported above. In LIV, *Cidec* had a strong SM/J bias in the high fat-fed male context, yet a LG/J bias in the high fat-fed females, low fat-fed females, and female context (**Figure 6G**). It was biallelically expressed in the remaining LIV contexts and across all WAT contexts, but was not expressed in HYP. *Cidec* ASE biases also had significant sex, diet, and diet-by-sex effects in LIV (p < 0.001, **Figure 6H**). We validated the context-dependent switch in ASE bias direction with pyrosequencing (**Figure 6I**). *Cidec* was also expressed in LIV at a significantly higher level in the high fat-fed males compared to the other diet-by-sex contexts (p < 0.01, **Figure 6J**).

*Cidec* promotes and regulates lipid storage in liver and adipose tissue (Xu et al. 2012). *Cidec* expression is sensitive to diet composition and sex hormones; for example, male mice had higher total hepatic expression levels than females when fed a Western diet (i.e. high cholesterol and saturated fats) (Herrera-Marcos et al. 2020). In another study, mice fed a high fat diet had significantly increased *Cidec* expression levels in liver that were also associated with improved insulin sensitivity (Iv et al. 2015). The liver plays an important role in regulating homeostatic insulin levels and removing it from the bloodstream (Tokarz et al. 2018). Hepatic insulin clearance is an emerging component of type 2 diabetes and metabolic syndrome, as impaired hepatic insulin clearance is correlated with increased waist circumference, blood pressure, fasting glucose, triglycerides, and insulin secretion (Najjar and Perdomo 2019; Pivovarova et al. 2013). Taken together, these data implicate *Cidec* as a putative candidate gene underlying *Ddiab6d* whose differential allelic composition in the liver is sensitive to metabolically-relevant environmental factors and can influence insulin levels.

## DISCUSSION

Allele-specific expression imbalances due to genetic and epigenetic variation have widespread functional consequences on complex traits, but how environmental signals contribute to this crosstalk remains understudied. Here, we explored how allelic genotype, parent-of-origin, tissue type, sex, and dietary fat simultaneously impact ASE biases in an adult mouse F_1_ reciprocal cross. We present a comprehensive genome-wide map of both parent-of-origin and sequence dependent ASE patterns across three metabolically-relevant tissues and nine environmental contexts. The granularity of our analyses revealed that both types of ASE are highly dependent on tissue type and environmental context. We identified 2,853 genes with significant parental and/or sequence biases and sorted them into three major expression profiles: tissue-independent, tissue-dependent, and context-dependent. Interestingly, we also found 45 genes with inconsistent ASE biases that switched direction across tissues and/or contexts (e.g. SM/J biased in one cohort, LG/J biased in another). Although the breadth of these patterns precludes a detailed discussion of each gene, we have validated examples of each expression profile to show how these allelic imbalances could manifest and lead to potential functional consequences.

“Tissue-independent” genes are strongly biased wherever they are expressed. Most parent-of-origin dependent ASE genes (67%) have this expression profile – all of which are related to genomic imprinting, a well-characterized epigenetic phenomenon that results in uniparental expression and is often conserved across tissues (Barlow and Bartolomei 2014). In contrast, only 23% of sequence dependent ASE genes have this expression profile, likely due to strain-specific *cis*-acting genetic variants in coding regions that severely affect one allele’s function whenever that gene is expressed. Both patterns conform to the conventional mechanism for ASE whereby the genetic and epigenetic processes that influence the allelic composition will always do so wherever a gene is expressed, in a manner impervious to tissue type or environmental signals (Lo et al. 2003; Babak et al. 2015). However, we found this is not always the case.

“Tissue-dependent” genes are moderately biased in one direction in some tissues, but are biallelically expressed (no bias) or biased in the opposite direction in other tissues. Tissue-specific gene expression (simply not expressed in certain tissues) is not considered here, as we are specifically interested in cases where a gene has one allelic ratio in one tissue (e.g. 80% SM/J, 20% LG/J) and a different allelic ratio in another tissue (e.g. 50% SM/J, 50% LG/J). Merely 6% of parent-of-origin dependent ASE genes have this expression profile and all are located in known imprinted domains. 25% of sequence dependent ASE genes also have this expression profile, demonstrating that sequence biases are often not solely attributable to genetic variation in coding regions. Instead, both patterns may be explained by variation in tissue-specific regulatory features, such as enhancers or RNA binding proteins. Such tissue-specific factors can then interact with *cis*-acting variants to produce a tissue-dependent allelic imbalance. This is likely the case for genes with inconsistent biases between tissues, where the variation in one allele may be favorable in certain tissues’ regulatory landscape but the opposite allele is more favorable elsewhere (Andergassen et al. 2017; Leung et al. 2015).

Finally, “context-dependent” genes are subtly biased in one direction in certain environmental contexts, but are biallelically expressed or biased in the opposite direction in other contexts and tissues. We found this expression profile is more prevalent than expected in both classes of ASE. 27% of parent-of-origin dependent ASE genes have diet- and/or sex-specific biases in a less extreme manner than the other profiles. Most genes have no clear connection to imprinting, suggesting another mechanism for parental ASE outside of traditional imprinting that is sensitive to environmental perturbations (Wolf et al. 2008; Morcos et al. 2011; Macias-Velasco et al. 2021). Surprisingly, over half of sequence dependent ASE genes (52%) have diet- and/or sex-specific biases, indicating these intrinsic and extrinsic environmental factors interact with strain-specific genetic variation to cause a sequence bias. Both patterns suggest a model where environmental signals may interact more efficiently with one allele over the other, leading to shifting and inconsistent allelic proportions in response to environmental cues (Shao et al. 2019). Often, these genes are not significantly biased in our “all contexts” analysis, only in a specific diet and/or sex cohort. This highlights the necessity of studying gene-by-environment interactions, as such effects are obscured when multiple contexts are collapsed together.

It is important to note that we can only detect ASE in genes with strain-specific variants in transcribed regions, which is a fraction of the total set of expressed genes. In particular, imprinted genes that are crucial for development/survival may be under tight evolutionary control and not have variants, thus we are unable to assess their ASE status here. Although our exact findings are limited to this F_1_ reciprocal cross mouse model, the broad patterns nevertheless demonstrate the complexity of allele-specific gene regulation and its contribution to complex traits. Traditional mapping studies connect genotypes to phenotypes, but are agnostic to tissue and often do not consider environment. eQTL studies connect genotypes to total gene expression levels, but assume biallelic expression. Tissue- and context-dependent ASE of both classes (parent-of-origin and sequence dependent) can bridge these approaches and pinpoint what tissues and/or environments are relevant for a phenotype. A gene could be expressed at the same level in two cohorts, but its allelic composition could differ due to ASE. If there is a functional difference between the two alleles, then an expression imbalance in the right tissue and/or environment could lead to phenotypic consequences. We integrated ASE patterns and QTL data from populations derived from the same founder strains at various degrees of intercrossing. Tissue-specific ASE genes are often enriched in QTLs for metabolic and musculoskeletal traits, yet comprise a small portion of all genes within a QTL. Incorporating these dynamic ASE patterns with orthogonal evidence will help us decipher the genotype to phenotype map.

## METHODS

### F_1_ Reciprocal Cross Mouse Model

We obtained LG/J and SM/J founders from The Jackson Laboratory (Bar Harbor, ME) and generated F_1_ reciprocal crosses by mating LG/J mothers with SM/J fathers (**LxS**) and vice versa (**SxL**). F_1_ offspring were weaned into sex-specific cages at three weeks of age and randomly placed on either a high fat diet (42% kcal from fat; Teklad TD88137) or an isocaloric low fat diet (15% kcal from fat; Research Diets D12284). They were fed *ad libitum*. F_1_ mice were euthanized at 20 weeks of age with a sodium pentobarbital injection followed by cardiac perfusion with phosphate-buffered saline. We harvested hypothalamus (**HYP**), liver (**LIV**), and reproductive white adipose (**WAT**) tissue, which were flash frozen in liquid nitrogen and stored at -80°C until RNA extraction. All procedures were approved by the Institutional Animal Care and Use Committee at Washington University School of Medicine.

### RNA-Sequencing and Allele-Specific Mapping

We sequenced 32 samples per tissue, representing 4 mice from each sex (male or female), diet (high or low fat), and F_1_ reciprocal cross (LxS or SxL) cohort. We extracted total RNA from WAT and HYP using the RNeasy Lipid Tissue Mini Kit (QIAGEN) and from LIV using a standard TRIzol-chloroform procedure. Samples were selected based on sufficient NanoDrop RNA concentrations (ThermoFisher) and RNA integrity scores ≥8.0 (Agilent). We constructed RNA-Seq libraries with the RiboZero rRNA Removal Kit (Illumina), checked their quality with the BioAnalyzer DNA 1000 assay (Agilent), and sequenced them at 100bp, paired-end reads on an Illumina HiSeq 400. After sequencing, reads were de-multiplexed and assigned to individual samples.

Quantifying ASE in a F_1_ reciprocal cross is vulnerable to reference genome alignment bias. If one parental strain is more closely related to the reference genome, then their reads tend to have a higher mapping quality than reads from the other parental strain (Wang and Clark 2014; Degner et al. 2009). We mitigated this concern by aligning RNA-Seq reads to a custom LG/J x SM/J “merged genome”. Previously, LG/J and SM/J reference genomes were created by combining strain-specific SNVs and indels with the GRC38.72-mm10 reference template (Nikolskiy et al. 2015). Customized gene annotations were also created by adjusting Ensembl definitions (Mus_musculus.GRCm38.72) for indexing differences due to strain-specific indels. Here, those LG/J and SM/J genomes were combined into one pseudogenome so we could align reads to both parental strains simultaneously.

We mapped reads uniquely using the two-pass mapping strategy in STAR v.2.7.2b (Dobin et al. 2013). Briefly, splice junctions are collected during the first round and used to inform a second round of mapping, thus detecting more reads that span novel junctions. By not allowing multi-mapping, we only retained reads uniquely covering strain-specific variants so we could assign reads to their allelic origin; reads covering identical regions between the parental strains were discarded. STAR alignment summaries are provided in **Supplemental Table S8** and **Supplemental Figure S18**.

Next, we assigned each aligned read to a gene using bedtools v.2.27.1 (Quinlan and Hall 2010) and our strain-specific Ensembl annotations. We did not assess ASE at the exon-level because not all exons within a given gene contained an informative variant. Gene-level allele-specific read counts were then upper quartile normalized (**Supplemental Figure S19**) and filtered to remove lowly-expressed genes (total normalized read counts < 20). We retained a total of 9,171 genes with detectable allele-specific expression in HYP, 9,761 genes in WAT, and 8,082 genes in LIV.

### Library Complexity

Insufficient library complexity can also hamper detecting ASE in a F_1_ reciprocal cross. In lowly or moderately expressed genes, if a read fragment from only one of the two alleles is randomly subjected to a duplicating event (i.e. PCR jackpotting) during RNA-Seq library construction, then that gene may spuriously appear as monoallelically expressed, even though it is a false positive (Wang and Clark 2014). To check for this, we measured each library’s complexity by fitting the distribution of LG/J allele expression biases (the proportion of total allele-specific read counts with the LG/J haplotype) to a beta-binomial distribution using the VGAM package (Yee 2010). We estimated the shape parameters (α, β) of the beta-binomial distribution and calculated the overdispersion parameter (ρ) as ρ = 1 / (1 + α + β). Lower values of ρ (< 0.075) indicate a library is sufficiently complex. One WAT library (CCGGACC) and one LIV library (TGATTAC) were deemed to have poor complexity and were removed from further analyses (**Supplemental Figure S20**).

### Determining Biased Allele-Specific Expression

To explore how dietary fat and/or sex impacts ASE patterns, we analyzed nine environmental contexts per tissue: high fat diet (**H**), low fat diet (**L**), females (**F**), males (**M**), high fat-fed females (**HF**), high fat-fed males (**HM**), low fat-fed females (**LF**), low fat-fed males (**LM**), and all contexts collapsed (**All**). In total, we analyzed 27 “tissue-by-context” cohorts (3 tissues x 9 contexts) (**Figure 1**). For each “tissue-by-context” analysis, we required a gene to be expressed in ≥75% of biological replicates for each F_1_ reciprocal cross: 3 mice per cross for diet-by-sex specific contexts (N = 4 mice per cross), 6 mice per cross for diet- and sex-specific contexts (N = 8), and 12 mice per cross for the “All” contexts (N = 16). This conservative sample size threshold retains only genes consistently expressed in most biological replicates in a cohort, allowing us to confidently conclude their expression status in that tissue and/or environment. This approach intentionally removes genes that are sporadically expressed within a cohort (e.g. from technical error or intrinsic stochasticity), as they may result in false or transient allelic biases.

We adapted a previously-published regression model (Takada et al. 2017) for jointly estimating parent-of-origin (**PO**) and allelic genotype (**AG**) effects on ASE. In our dataset, each row corresponds to a specific allele; each individual mouse/RNA-Seq library is represented over two rows. First, we assigned two binary variables to each gene’s allele-specific counts based on their allelic origin. For the PO term, maternal alleles received a 0 and paternal alleles received a 1; for the AG term, LG/J alleles received a 0 and SM/J alleles received a 1 (**Supplemental Figure S21**). Next, we added a term representing the RNA-Seq library barcode IDs so that both allele-specific read counts (both rows) for each individual mouse are treated as a pair. This paired-sample design helps distinguish between cases where a gene is not expressed in a particular sample (i.e. both alleles’ read counts are zero) and an extreme expression bias (i.e. only one allele’s counts are zero), which results in a lower biological coefficient of variation and lower overall dispersion (**Supplemental Figure S21**). Our final regression model equation is as follows:

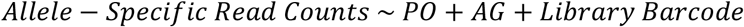

Finally, we fit a negative binomial generalized linear model (GLM) using this equation and conducted likelihood ratio tests in edgeR (Robinson et al. 2010; McCarthy et al. 2012) to estimate the parent-of-origin (PO term) and allelic genotype (AG term) effects on allele-specific gene expression levels.

Next, we quantified the direction and magnitude of each gene’s expression biases. For each sample, we calculated a gene’s allelic bias as the proportion of total read counts with the LG/J haplotype (L_bias_) or the SM/J haplotype (S_bias_). Using the mean allelic biases of each F_1_ reciprocal cross, we constructed Parent-of-Origin Effect (**POE**) and Allelic Genotype Effect (**AGE**) scores per gene as follows:

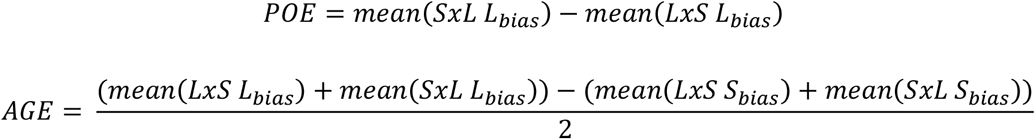

POE scores range from -1 (completely maternally expressed) to +1 (completely paternally expressed). Similarly, AGE scores range from -1 (completely SM/J expressed) to +1 (completely LG/J expressed). Scores of 0 for both indicate biallelic expression. Full GLM summary statistics and POE/AGE scores for each tissue-by-context analysis are provided in **Supplemental Tables S2 - S4**.

### Estimating Significance Thresholds

A crucial challenge for ASE analyses is how to best adjust for multiple tests. Allelic biases are often correlated for genes within and between imprinted domains (Edwards and Ferguson-Smith 2007) and for genes controlled by the same regulatory element with a functional variant (Cavalli et al. 2016), ensuring any tests performed are also correlated and breaking independence assumptions. To address this, we used a permutation approach to estimate our statistical and biological significance thresholds.

For each tissue-by-context analysis, we generated a permuted null distribution for both AG and PO terms. Over several iterations, we randomly shuffled the allele-specific read counts for all genes, reran our GLM analyses, and recalculated the POE/AGE scores. Instead of choosing an arbitrary number, we continued to add new iterations until the null model was “stable”. After every iteration, we added those permuted results to the overall null model and evaluated the difference in mean permuted likelihood ratios for both terms in the null model. We defined “stability” as when the difference in mean permuted likelihood ratios for both AG and PO terms deviated/fluctuated by < |0.001| for 10 consecutive iterations (**Supplemental Figure S22**). In other words, the null model was considered “stable” once incorporating a new iteration did not shift the overall null distribution of test statistics (likelihood ratios). Across the 27 tissue-by-context analyses, the permuted datasets included a mean of 51 iterations. Comparisons between the real and permuted datasets are provided for p-values, POE/AGE scores, and likelihood ratios (**Supplemental Figures S23 – S25**).

Next, we built an empirical cumulative distribution function (ECDF) from each term’s permuted p-values for each tissue-by-context analysis (**Supplemental Figures S26**). We fit each term’s raw p-values to its respective ECDF to compute the adjusted p-values, i.e. the proportion of tests from the permuted null model that are more extreme (smaller p-values) than the test from the real model. We also calculated the 5^th^ and 95^th^ quantiles of the permuted POE/AGE scores for each term in each tissue-by-context analysis. We set our critical threshold as the more extreme value. We deemed an adjusted p-value ≤ 0.05 as statistically significant and real POE/AGE scores beyond the critical threshold as biologically significant.

Genes with significant PO term p-values and POE scores were considered to have parent-of-origin dependent ASE. Similarly, genes with significant AG term p-values and AGE scores were considered to have sequence dependent ASE (**Supplemental Figures S27, Supplemental Table S1**).

### Total Expression Mapping

Quantifying total expression in F_1_ heterozygotes is challenging because both parental genomes should be included to minimize reference genome alignment bias. However, incorporating both genomes doubles the amount of genomic space, so typical RNA-Seq alignment parameters that allow for multi-mapping are too stringent. During our previous allele-specific mapping step, reads were not aligned if they covered homozygous regions between the parental strains. Here, we took those unmapped reads and aligned them separately to the LG/J and SM/J reference genomes using a standard multi-mapping approach in STAR v.2.7.2b. We assigned each aligned read to a gene using bedtools v.2.27.1 and our strain-specific Ensembl annotations. Next, we averaged together the gene-level read counts for the LG/J and SM/J genomes; since these reads covered identical regions between the strains, they aligned to each strain’s genome similarly. We then combined these averaged “multi-mapped” read counts with the raw “allele-specific” read counts from the previous mapping step, resulting in raw “total expression” read counts. Finally, gene-level read counts were upper quartile normalized and filtered to remove lowly-expressed genes (total read counts < 10). We retained 25,926 genes with adequate total expression levels in HYP, 26,450 genes in WAT, and 24,076 genes in LIV.

### Characterizing ASE Profiles: Tissue-Independent, Tissue-Dependent, and Context-Dependent

To examine how ASE is influenced by tissue type and/or environmental context, we characterized the expression profiles of the significant ASE genes across our 27 tissue-by-context analyses (3 tissues x 9 diet-by-sex contexts). In each analysis, a gene could be expressed in one of three ways: significantly biased, expressed with no allele-specific bias (biallelic), or simply not expressed. We sorted the significant ASE genes of both classes into three expression profiles based on the following criteria.

Tissue-independent ASE genes have a significant bias in every tissue where they are expressed. Within a tissue, the gene must have a significant bias in ≥5 of the 9 diet-by-sex contexts (i.e. most, but not all, environmental contexts). This flexibility allows for genes that may have true ASE but were excluded due to our stringent minimum sample size requirements; these genes could appear as biallelically expressed since they still pass our minimum read depth requirements. Tissue-dependent ASE genes have a significant bias in some tissues and no bias in others. The gene must have a significant bias in >5 of the 9 contexts in one or two tissues, but biallelic expression in >5 of the 9 contexts in the other tissue(s). Both profiles allowed for genes to not be expressed in some tissues, as we wanted to deliberately distinguish between tissue-specific gene expression (not expressed in certain tissues) and tissue-dependent ASE (biased in certain tissues). Finally, context-dependent ASE genes have a significant bias only in certain environmental contexts and no bias elsewhere. The gene must have a significant bias in ≤5 of the 9 contexts within a tissue, but biallelic expression in the other contexts and/or tissues. All significant ASE genes of both classes fit into one of these expression profiles; no genes were unclassified.

### Evaluating Environmental Context-Dependency

Once we identified context-dependent genes, we evaluated how sex and/or dietary environment shape their ASE patterns. For each context-dependent gene, we calculated each sample’s allelic bias as the proportion of total read counts with the LG/J haplotype (L_bias_) or the SM/J haplotype (S_bias_). We constructed individualized Parent-of-Origin Effect (POE) or Allelic Genotype Effect (AGE) scores per sample per gene by modifying the above equations as follows:

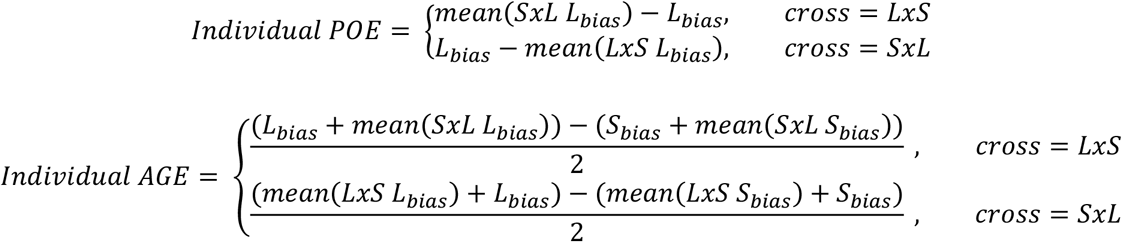

Individualized POE scores range from -1 (maternally expressed) to +1 (paternally expressed). Similarly, individualized AGE scores range from -1 (SM/J expressed) to +1 (LG/J expressed). Scores of 0 for both scores indicate biallelic expression or that the gene was not expressed in that sample.

Next, we used ANOVA models to test whether a gene’s allelic biases (individualized POE/AGE scores) were influenced by sex, diet, and/or their interaction. We considered FDR-corrected p-values ≤ 0.1 to be significant (**Supplemental Table S5**). For genes with significant diet-by-sex interactions, we conducted Tukey’s post-hoc tests to identify significant differences among diet-by-sex cohorts (adjusted p ≤ 0.05) (**Supplemental Table S6**).

### Chromosomal Location Enrichment

We performed over-representation analysis for chromosomal locations with the WEB-based GEne SeT AnaLysis Toolkit v2019 (Liao et al. 2019). For each tissue-by-context analysis, we analyzed the lists of genes with significant sequence or parent-of-origin biases against the background of all unique genes with detectable ASE in that analysis (**Supplemental Tables S2 – S4**). A Benjamini-Hochberg FDR-corrected p-value ≤ 0.1 was considered significant.

### Imprinted Gene List

We defined a gene as “canonically imprinted” if it appeared in the GeneImprint mouse database (https://www.geneimprint.com, as of May 2020) and/or a manual PubMed search of the gene name (or synonyms) with imprinting-related terms.

### Pyrosequencing

We validated examples of each allele-specific expression profile: tissue-independent, tissue-dependent, context-dependent, switching bias direction. We sorted genes in each ASE profile by total expression level. To minimize noise during the pyrosequencing reaction, we prioritized genes that were highly expressed and statistically significant (for context-dependent examples), but excluded those with high expression pattern variance between biological replicates. We randomly selected two or three genes fitting these criteria for each ASE profile (total validated = 13 genes). For each gene, we identified the strain-specific SNPs within exons and designed primer sets to flank these variants using Geneious Prime 2020.0.4 (https://www.geneious.com) (Kearse et al. 2012). Wherever possible, target regions were 150-200bp long and spanned an exon-exon junction to avoid genomic DNA contamination. We verified the specificity of each primer set *in silico* with Geneious and *in vitro* with PCR and Sanger sequencing. All primer sequences are provided in **Supplemental Table S9**.

We extracted total RNA from the HYP, WAT, and LIV of one mouse per F_1_ reciprocal cross (LxS and SxL) in each diet-by-sex cohort using the RNeasy Lipid Tissue Mini Kit (QIAGEN). Next, cDNA of each gene target was reverse-transcribed and PCR amplified with the PyroMark OneStep RT-PCR kit (QIAGEN) using one biotinylated (reverse) and one non-biotinylated (forward) primer. The biotinylated single-stranded PCR products were then purified with Streptavidin Sepharose High Performance beads (Cytiva) and hybridized to sequencing primers (same as forward) on the PyroMark vacuum prep workstation (QIAGEN). Finally, we performed pyrosequencing with the Allele Quantification program on the PyroMark Q24 system (QIAGEN). The pyrosequencing reaction emits a light signal as it builds the DNA fragment, which appears on the pyrogram output as a peak whose height is proportional to how many nucleotides were incorporated at that base. From these peaks, the PyroMark Q24 software quantified the allelic ratio of the variable position(s) in each gene’s assay. We calculated the mean allelic ratios of each variant in each tissue-by-context cohort.

### Integrating ASE and QTL Data

The genotypes, phenotypes, and statistical methods used to map QTL in the F_16_ and F_50-56_ generations of the LG/J x SM/J advanced intercross line (**AIL**) are described elsewhere (Lawson et al. 2010, 2011b; Cheverud et al. 2011; Gonzales et al. 2018; Cordero et al. 2019). The F_16_ QTL were mapped to the NCBI37/mm9 mouse reference genome, so we converted those QTL intervals to their GRCm38/mm10 coordinates with the UCSC Genome Browser’s liftOver tool (https://genome.ucsc.edu/cgi-bin/hgLiftOver) (Lee et al. 2022). We also extracted the positions of significant ASE genes from the GRCm38/mm10 assembly annotation. The F_50-56_ QTL were originally mapped to the GRCm38/mm10 reference.

We analyzed the two ASE classes separately; we intersected sequence dependent ASE genes with F_16_ and F_50-56_ QTL showing additive effects as well as parent-of-origin dependent ASE genes with F_16_ QTL showing imprinting effects (**Supplemental Table S7**). Within each analysis, we counted how many ASE genes overlapped with each AIL QTL using bedtools v.2.27.1. We also sorted each AIL generation’s QTL based on their associated trait category (e.g. related to diabetes, obesity, serum lipids, behavior, muscle, or bone) and compared those trait-specific QTL sets with the tissue-specific ASE gene sets. Next, we conducted enrichment analyses to determine whether ASE genes are overrepresented in AIL QTLs (i.e. located within QTL more often than expected). We randomly shuffled the QTL windows around the genome and tallied how many ASE genes overlap with regions of those lengths by chance. These permutations were repeated over 10,000 iterations to establish a null model. We calculated a z-score based on the null model and the real ASE/QTL overlap, then sampled a Normal distribution to obtain a p-value. We considered p-values ≤ 0.05 to be significantly enriched.

## DATA ACCESS

All raw RNA-Sequencing data generated in this study have been submitted to the NCBI BioProject database (https://www.ncbi.nlm.nih.gov/bioproject/) under accession number PRJNA753198.

## ACKNOWLEDGEMENTS

This work was supported by the Washington University Department of Genetics, the Diabetes Research Center at Washington University (CFS: P30DK020579), the National Science Foundation Graduate Research Fellowship (CLS: DGE1745038, DGE2139839), as well as the National Institute of Diabetes and Digestive and Kidney Diseases (HAL: K01DK095003), the National Institute of Environmental Health Sciences (HAL: U24ES026699), and the National Human Genome Research Institute (JFMV: T32GM007067) of the National Institutes of Health. The authors declare no conflicts of interest.

