## Supplementary material for "Genetic, epigenetic, and environmental mechanisms govern allele-specific gene expression": StPierre2021_SupplementalMaterials.pdf

**Table S1: Summary of ASE analyses in all tissues and environmental contexts.**
ASE summaries for all 27 tissue-by-environmental context analyses (3 tissues x 9 contexts). This includes the number of samples per analysis, the input ("raw") number of genes in each tissue's aligned RNA-Seq dataset, the final number of genes remaining after filtering for read count and sample size minimums, and the number of permutation iterations. For both sequence/allelic genotype ("AG") and parent-of-origin ("PO") effects, the table contains the effect score significance thresholds and the number of genes with significant ASE biases.

**Table S2: Allele-specific expression analyses for all contexts in HYP.**
Full summary statistics for all environmental context analyses in HYP. Each gene's allelic genotype (AG) and parent-of-origin (PO) effects on ASE were jointly estimated with a generalized linear model and the magnitude of its ASE biases were quantified (see Methods). For both AG and PO terms, the table includes the $log_2$ fold-changes, $log_2$ counts-per-million, likelihood ratios, raw p-values, effect scores (AGE/POE), and ECDF-adjusted p-values of all genes. Each sheet contains a separate diet-by-sex context analysis. Genes with adjusted p-values ≤ 0.05 and AGE/POE scores below their threshold (Table S1) were considered to have significantly biased ASE.

**Table S3: Allele-specific expression analyses for all contexts in LIV.**
Full summary statistics for all environmental context analyses in LIV. Each gene's allelic genotype (AG) and parent-of-origin (PO) effects on ASE were jointly estimated with a generalized linear model and the magnitude of its ASE biases were quantified (see Methods). For both AG and PO terms, the table includes the $log_2$ fold-changes, $log_2$ counts-per-million, likelihood ratios, raw p-values, effect scores (AGE/POE), and ECDF-adjusted p-values of all genes. Each sheet contains a separate diet-by-sex context analysis. Genes with adjusted p-values ≤ 0.05 and AGE/POE scores below their threshold (Table S1) were considered to have significantly biased ASE.

**Table S4: Allele-specific expression analyses for all contexts in WAT.**
Full summary statistics for all environmental context analyses in WAT. Each gene's allelic genotype (AG) and parent-of-origin (PO) effects on ASE were jointly estimated with a generalized linear model and the magnitude of its ASE biases were quantified (see Methods). For both AG and PO terms, the table includes the $log_2$ fold-changes, $log_2$ counts-per-million, likelihood ratios, raw p-values, effect scores (AGE/POE), and ECDF-adjusted p-values of all genes. Each sheet contains a separate diet-by-sex context analysis. Genes with adjusted p-values ≤ 0.05 and AGE/POE scores below their threshold (Table S1) were considered to have significantly biased ASE.

**Table S5: Context-dependent ANOVA results for all tissues.**
Full summary statistics for the context-dependent ANOVA analyses in each tissue and ASE class. For each context-dependent gene, individualized POE/AGE scores were calculated and tested for significant diet, sex, and diet-by-sex interaction effects (see Methods). The table includes the degrees of freedom, sums of squares, mean sums of squares, F statistics, raw p-values, and FDR-adjusted p-values for each ANOVA model term. FDR-corrected p-values ≤ 0.05 were considered significant.

**Table S6: Context-dependent post-hoc results for all tissues.**
Full post-hoc summary statistics in each tissue and ASE class. Tukey's Honest Significant Differences tests were conducted on the individualized POE/AGE scores of genes with significant diet-by-sex interaction effects. The table includes the mean differences between groups, the lower and upper end points of the interval, and the adjusted p-value for every diet-by-sex context comparison. Adjusted p-values ≤ 0.05 were considered significant.

**Table S7: Intersection results for all ASE gene and AIL QTL comparisons.**
Intersection results for ASE genes and QTL mapped in the LG/J x SM/J Advanced Intercross Line. Sequence dependent ASE gene positions were intersected with additive AIL $F_{16}$ and $F_{50\text{-}56}$ QTL windows. Parent-of-origin dependent ASE gene positions were intersected with imprinting AIL $F_{16}$ QTL windows. For each QTL, the table includes the AIL generation, effect type, chromosomal position, associated phenotypes, and tally of all genes located within each QTL. The table also lists the number and names of overlapping significant ASE genes (total ASE genes and each tissue-specific ASE gene set).

**Table S8: STAR alignment summaries of RNA-Seq libraries.**
Sample information and STAR alignment rates for each RNA-Seq library. Sample information includes the tissue, library barcode, mouse ID, diet, sex, and $F_1$ reciprocal cross. Libraries were aligned uniquely to the LG/J x SM/J "merged genome" so RNA-Seq reads could be mapped to both parental genomes simultaneously (see Methods).

The STAR alignment rates include the total number of input reads as well as the number and proportion of reads that mapped uniquely, reads mapped to “too many” loci, and reads not mapped (too short or another reason).

**Table S9: Primer sequences used in pyrosequencing.**
Forward and reverse primer sequences for pyrosequencing of our selected genes. The length, chromosomal location in each strain’s genome, and number of SNPs captured in each amplified PCR product are also included. All reverse primers are biotinylated at the 5’ end (*). All forward primers were used as the sequencing primer.

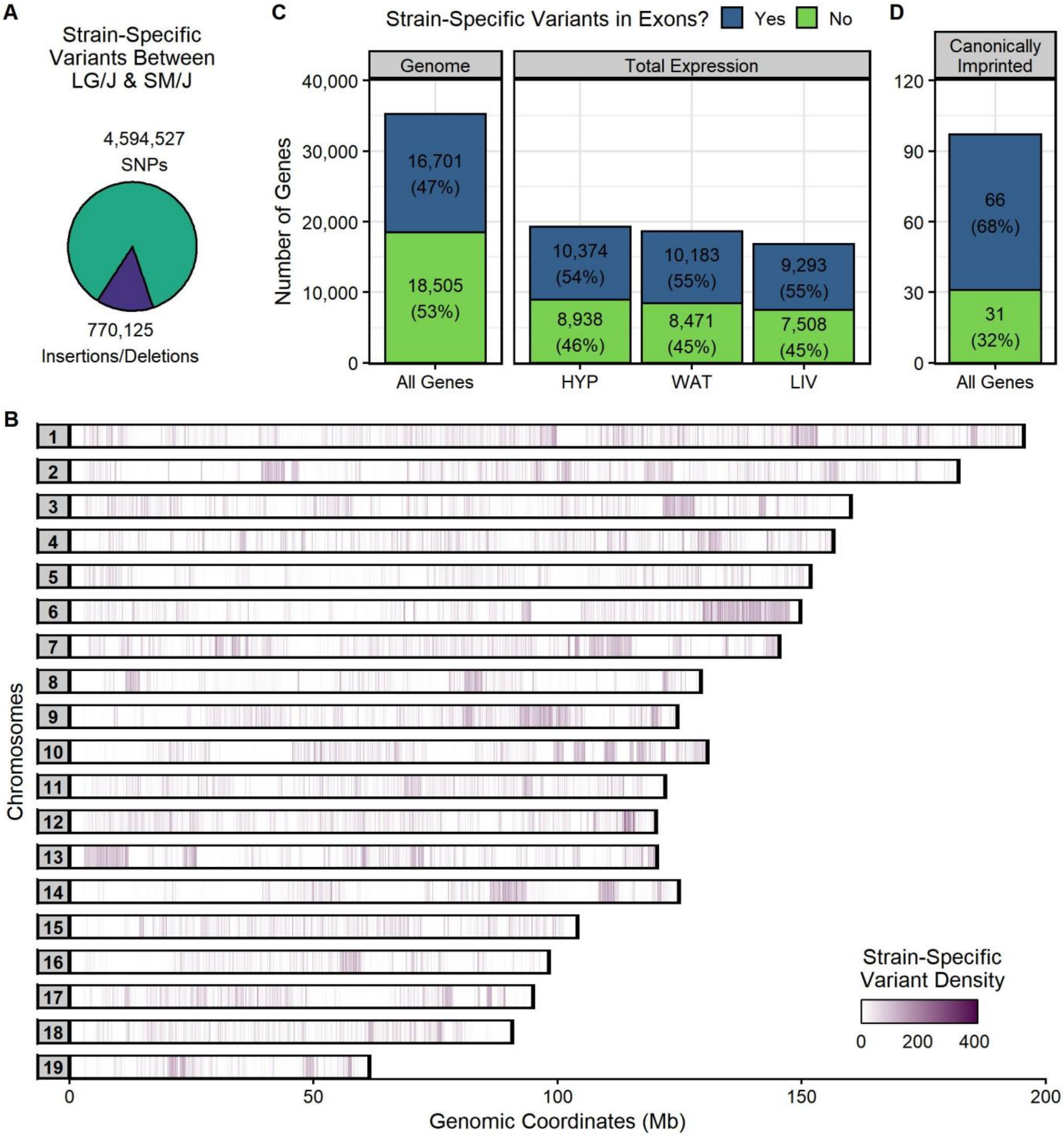

**Figure S1: Informative variants between the LG/J and SM/J strains are necessary to assess ASE.** Our ability to probe ASE in this $F_1$ reciprocal cross model depends on the frequency and distribution of strain-specific genetic variants between the parental inbred strains. **(A)** The LG/J and SM/J genomes differ by ~5.36 million single nucleotide polymorphisms (SNPs) and short (<50 bp) insertions/deletions. **(B)** These variants are densely clustered in certain regions and sparse in others. We are unable to evaluate ASE in homozygous regions between these two strains. On this genomic map, the x-axis denotes genomic coordinates (in Megabases) and the y-axis is grouped by chromosome. 10 kb genomic windows are color-coded by the number of variants within them. **(C)** To be informative for ASE, a gene must have at least one strain-specific variant in its annotated exon regions: this criteria applies to 47% of all genes in the mm10 reference genome (left) and 54-55% of total expressed genes in our sequenced tissues (right). Total expression is based on a standard multi-mapping approach that does not consider allelic origin (see Methods). **(D)** Known imprinted genes are often crucial for development/survival; if they are under tight evolutionary control and do not contain variants, then we cannot assess their ASE. However, 68% of “canonically imprinted” mouse genes do contain a strain-specific variant in their annotated exon regions (definition based on the GeneImprint.com database, see Methods).

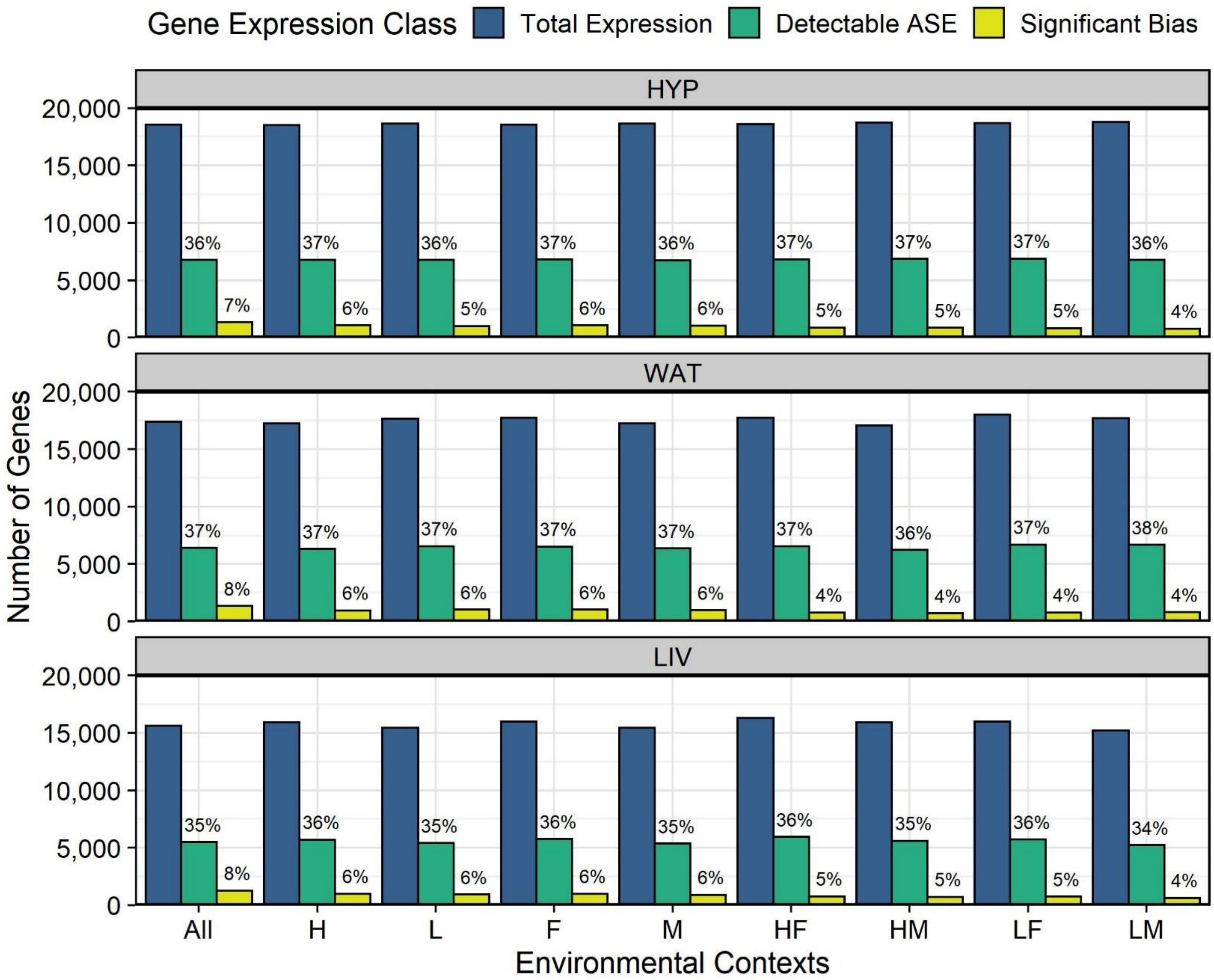

**Figure S2: ~37% of expressed genes can be probed in an allele-specific manner between LG/J and SM/J.** Across our tissue-by-context analyses, 15,500 – 18,800 genes were expressed when RNA-Seq reads are aligned with a standard multi-mapping approach that does not consider allelic origin ("total expression", dark blue bars, see Methods). 5,200 – 6,900 of these genes had detectable allele-specific expression that passed our stringent minimum read count and sample size thresholds ("detectable ASE", green bars). Thus, ~37% of all expressed genes in these tissues can be probed in an allele-specific manner between the LG/J and SM/J strains. This proportion is different from **Figure S1C** because ASE mapping is restricted to a subset of exons that contain informative variants; some genes are removed from the ASE dataset due to low read counts but are retained in the total expression dataset. Finally, 600 – 1400 (~5%) of genes had a significant allelic bias in a sequence or parent-of-origin dependent manner ("significant bias", yellow bars). Analyses are grouped by tissue: hypothalamus (HYP), white adipose (WAT), and liver (LIV). They are then ordered on the x-axis by environmental context: all contexts collapsed (All), high fat diet (H), low fat diet (L), females (F), males (M), high fat-fed females (HF), high fat-fed males (HM), low fat-fed females (LF), and low fat-fed males (LM). The y-axis denotes the number of genes in each expression class. Proportions shown are relative to the number of total expressed genes for that tissue-by-context cohort.

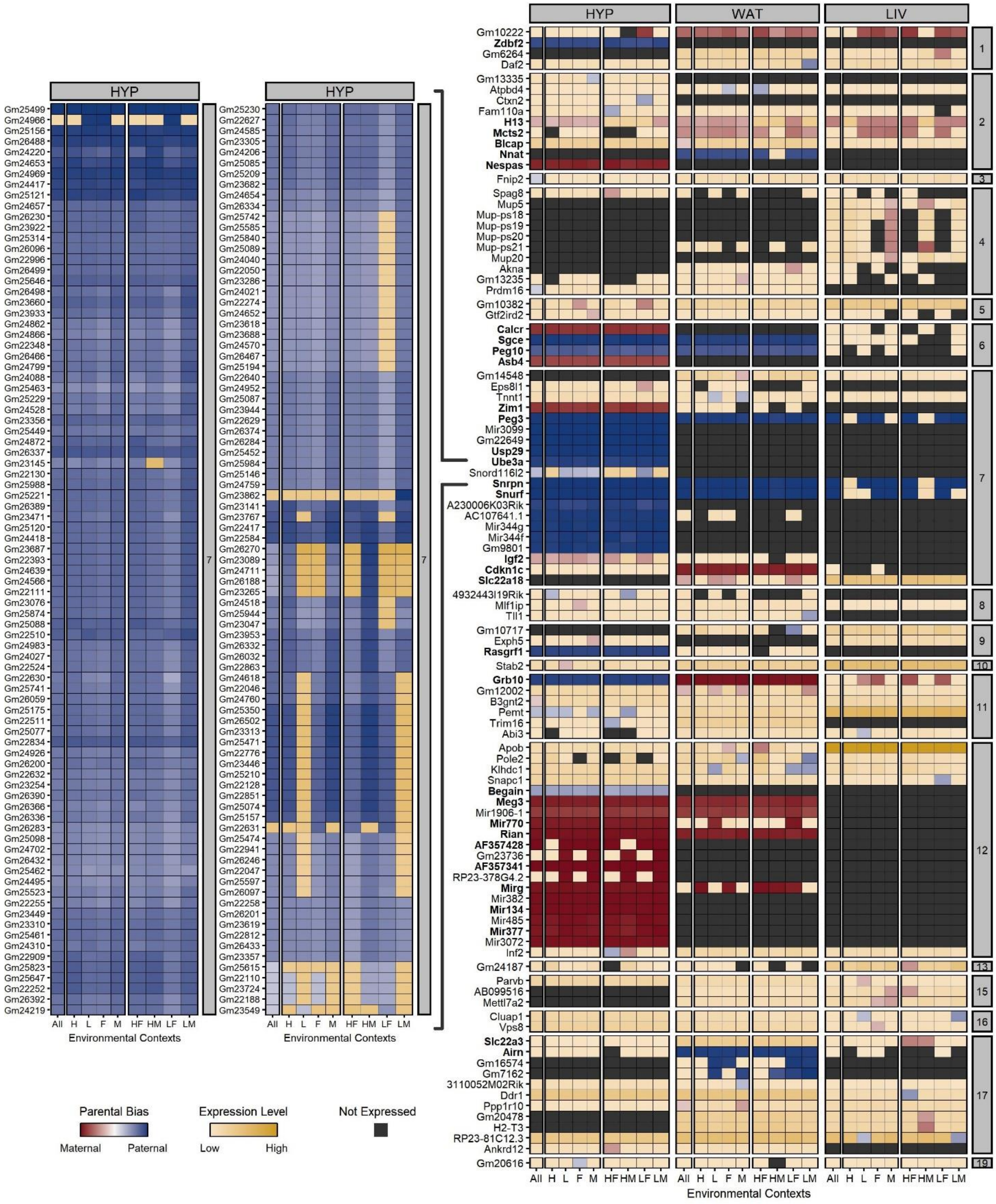

**Figure S3: Full heatmap of parent-of-origin dependent ASE profiles across tissues and contexts.** Heatmap of all 271 genes with significant parental biases, including the 672 kb cluster of small nucleolar RNAs located in the Prader-Willi/Angelman syndrome domain on chr7 which contains canonically imprinted genes (e.g. *Ube3a* and *Snrpn*). All 170 snoRNAs are paternally biased and only expressed in HYP. Genes are color-coded by their expression pattern in each tissue-by-context analysis. Shades of red and blue indicate their degree of maternal or paternal bias, respectively (POE scores). Where genes are not significantly biased, shades of yellow indicate their biallelic expression levels (log-transformed total counts). Black indicates genes are not expressed in that analysis. Bolded genes are canonically imprinted. The y-axis is sorted by chromosomal position. Each supercolumn denotes a tissue: hypothalamus (HYP), white adipose (WAT), and liver (LIV). Each subcolumn denotes an environmental context: all contexts (All), high fat diet (H), low fat diet (L), females (F), males (M), high fat-fed females (HF), high fat-fed males (HM), low fat-fed females (LF), and low fat-fed males (LM).

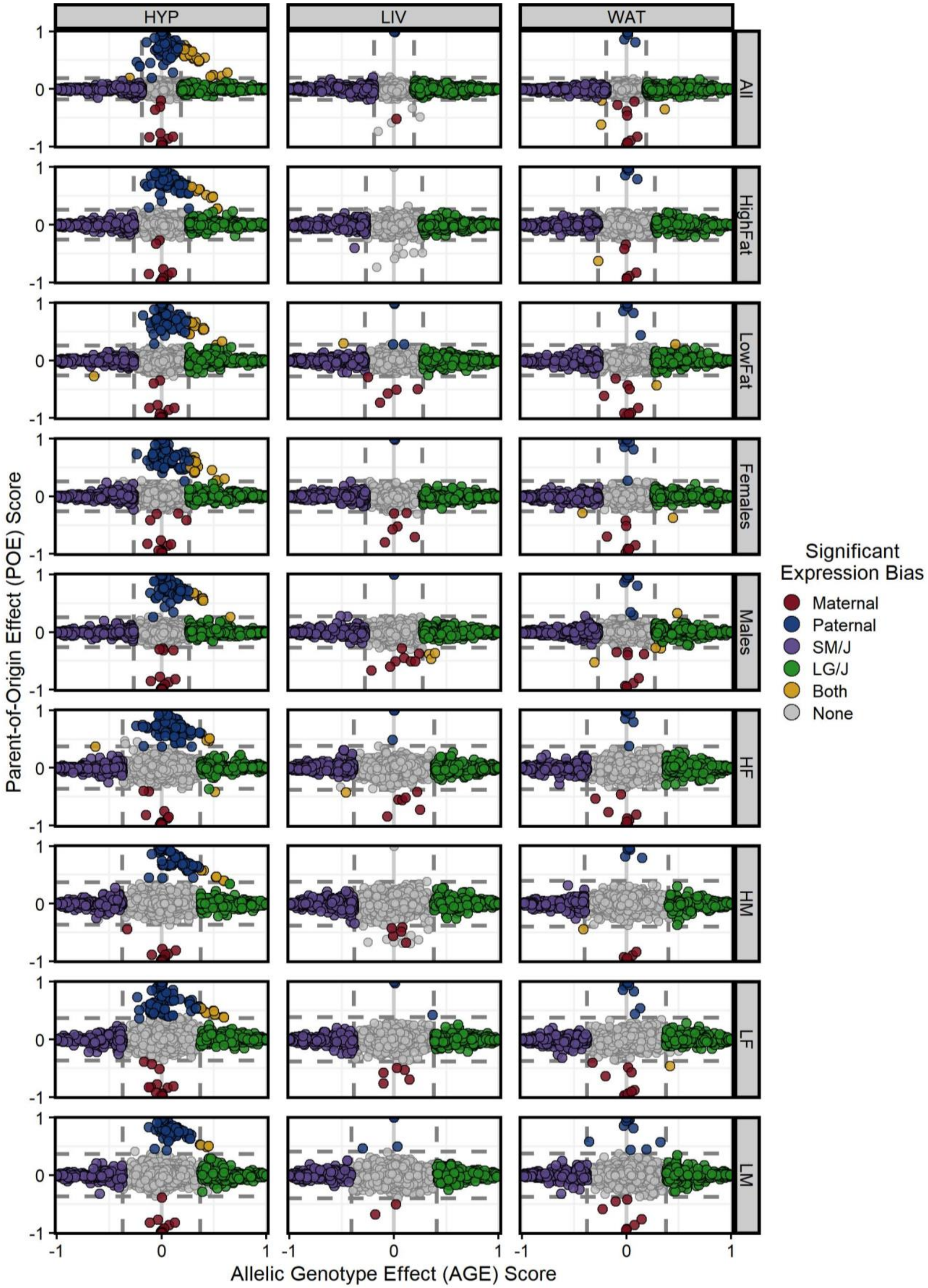

**Figure S4: Comparison of allelic bias magnitudes across each tissue-by-context analysis.** Parent-of-Origin Effect (POE, y-axis) versus Allelic Genotype Effect (AGE, x-axis) scores in each tissue-by-context analysis. Genes with significant parental biases (red/blue) have extreme POE scores but weak AGE scores. Genes with significant sequence biases (purple/green) have extreme AGE scores but weak POE scores. Some genes have both parental and sequence biases (yellow). Most genes have no ASE bias (gray). Columns are grouped by tissue and rows are grouped by environmental context. Dashed lines indicate score thresholds.

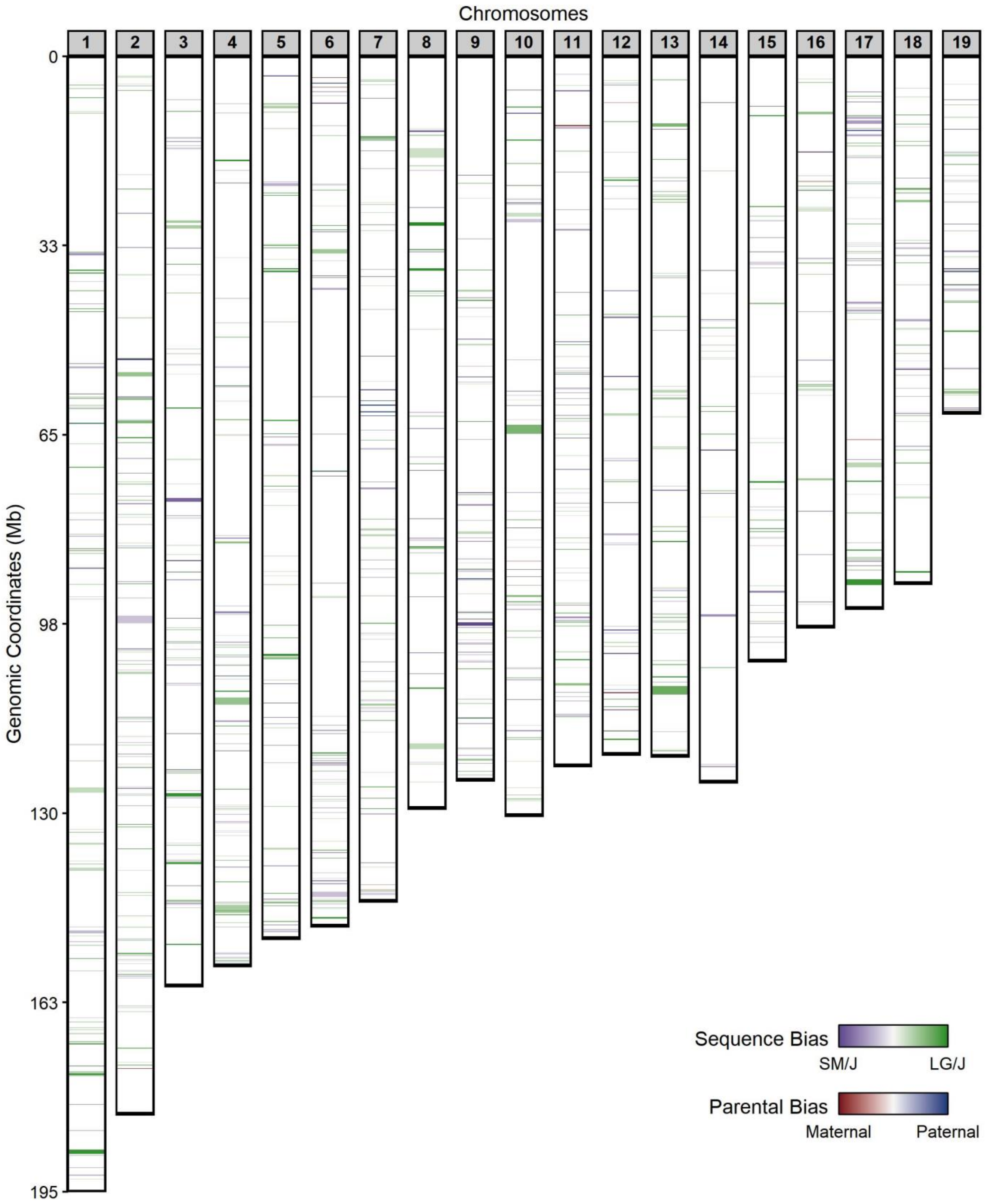

**Figure S5: Chromosome map of all significant ASE genes.**
Both parent-of-origin and sequence dependent ASE patterns occurred genome-wide. Parentally biased genes tended to cluster in known imprinted domains, such as the *Ube3a/Snrpn* (chr 7), *Meg3/Gtl2* (chr 12), *Peg3/Usp29* (chr 7), and *H13/Mcts2* (chr 2) domains. Sequence biased genes were spread more diffusely across the genome, but occasionally clustered in distinct regions. The y-axis denotes genomic coordinates (in Megabases) and the x-axis is grouped by chromosome. Gene windows for each significant ASE gene are color-coded by the degree of their parental (red to blue) or sequence (purple to green) bias. For each gene, the most extreme Parent-of-Origin Effect or Allelic Genotype Effect score of the 27 tissue-by-context analyses was plotted as representative.

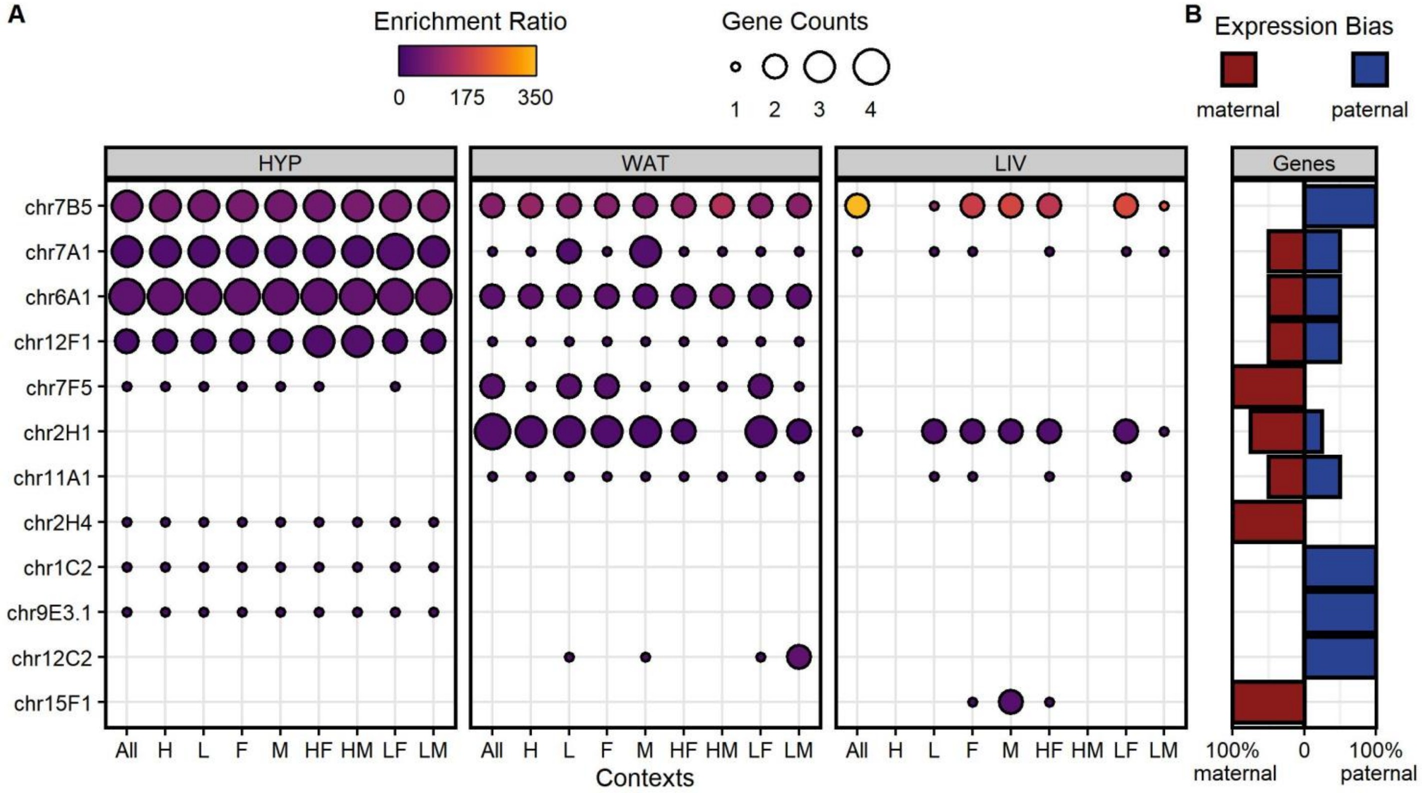

**Figure S6: Chromosomal domains enriched in parent-of-origin dependent ASE genes.**
We performed over-representation analysis for chromosomal locations with the WEB-based GEne SeT AnaLysis Toolkit. Genes with parental biases often cluster in canonical imprinted domains. **(A)** Chromosome domains enriched among the significant parent-of-origin dependent ASE genes in each tissue-by-context analysis. Domains are sorted by significance on the y-axis (mean p-value across all analyses, low to high). Dot color is proportional to the enrichment ratio (the number of observed/expected genes in that domain) and dot size is proportional to the number of significant genes within each domain. Analyses are grouped by tissue on the x-axis, then ordered by environmental context. No dot is plotted if a domain is not significantly enriched ($p > 0.1$) among that analysis' gene list. **(B)** Expression bias directions of the significant parent-of-origin dependent ASE genes within each chromosomal domain (all tissue-by-context analyses collapsed). Divergent bars indicate the percentage of maternally (red) and paternally (blue) biased genes in each domain.

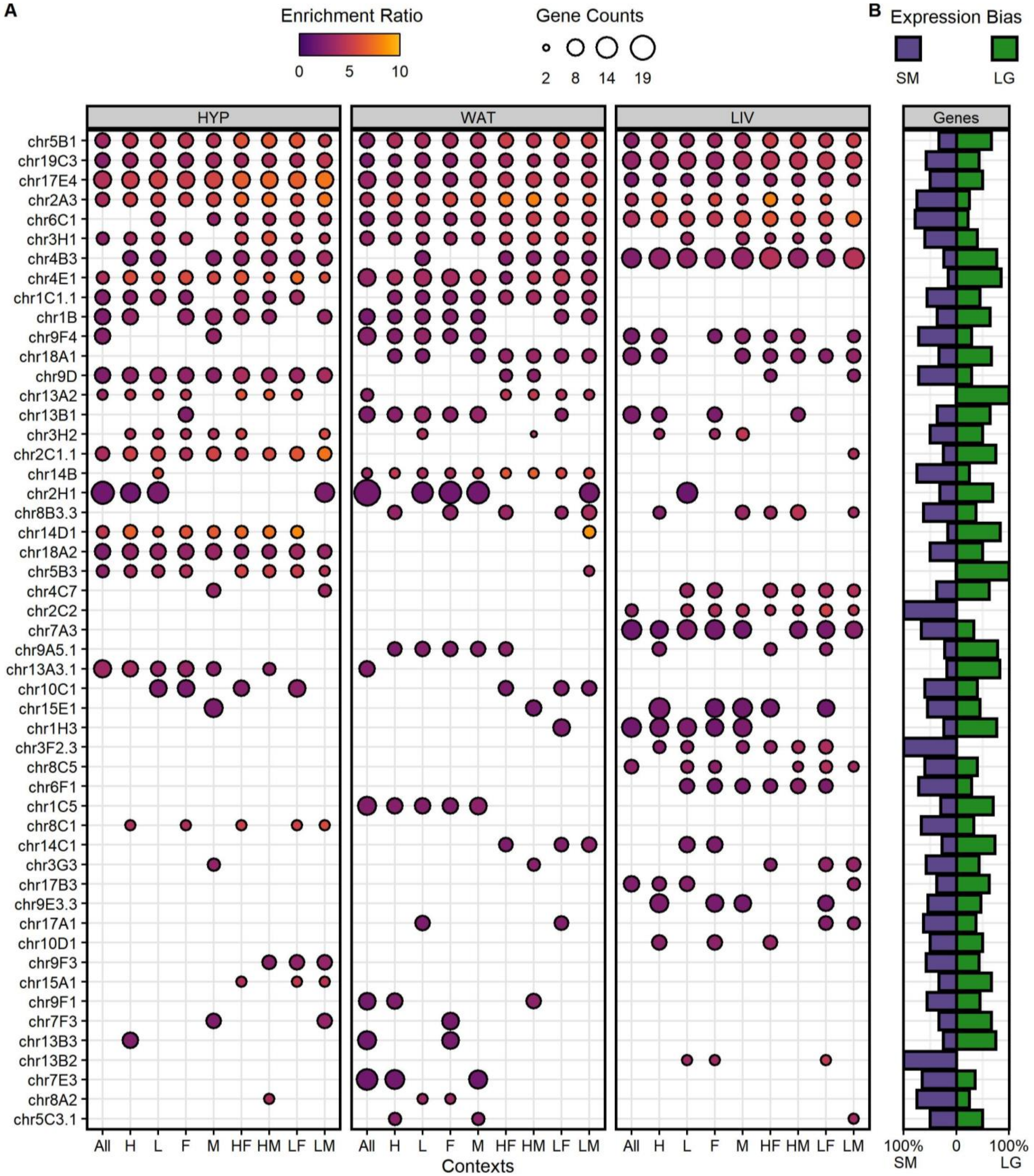

**Figure S7: Chromosomal domains enriched in sequence dependent ASE genes.**
We performed over-representation analysis for chromosomal locations with the WEB-based GEne SeT AnaLysis Toolkit. Genes with sequence biases cluster in regions potentially controlled by the same regulatory element. **(A)** Chromosome domains enriched among the significant sequence dependent ASE genes in each tissue-by-context analysis. Domains are sorted by significance on the y-axis (mean p-value across all analyses, low to high). Dot color is proportional to the enrichment ratio (the number of observed/expected genes in that domain) and dot size is proportional to the number of significant genes within each domain. Analyses are grouped by tissue on the x-axis, then ordered by environmental context. No dot is plotted if a domain is not significantly enriched ($p > 0.1$) among that analysis' gene list. **(B)** Expression bias directions of the significant sequence dependent ASE genes within each chromosomal domain (all tissue-by-context analyses collapsed). Divergent bars indicate the percentage of SM/J (purple) and LG/J (green) biased genes in each domain.

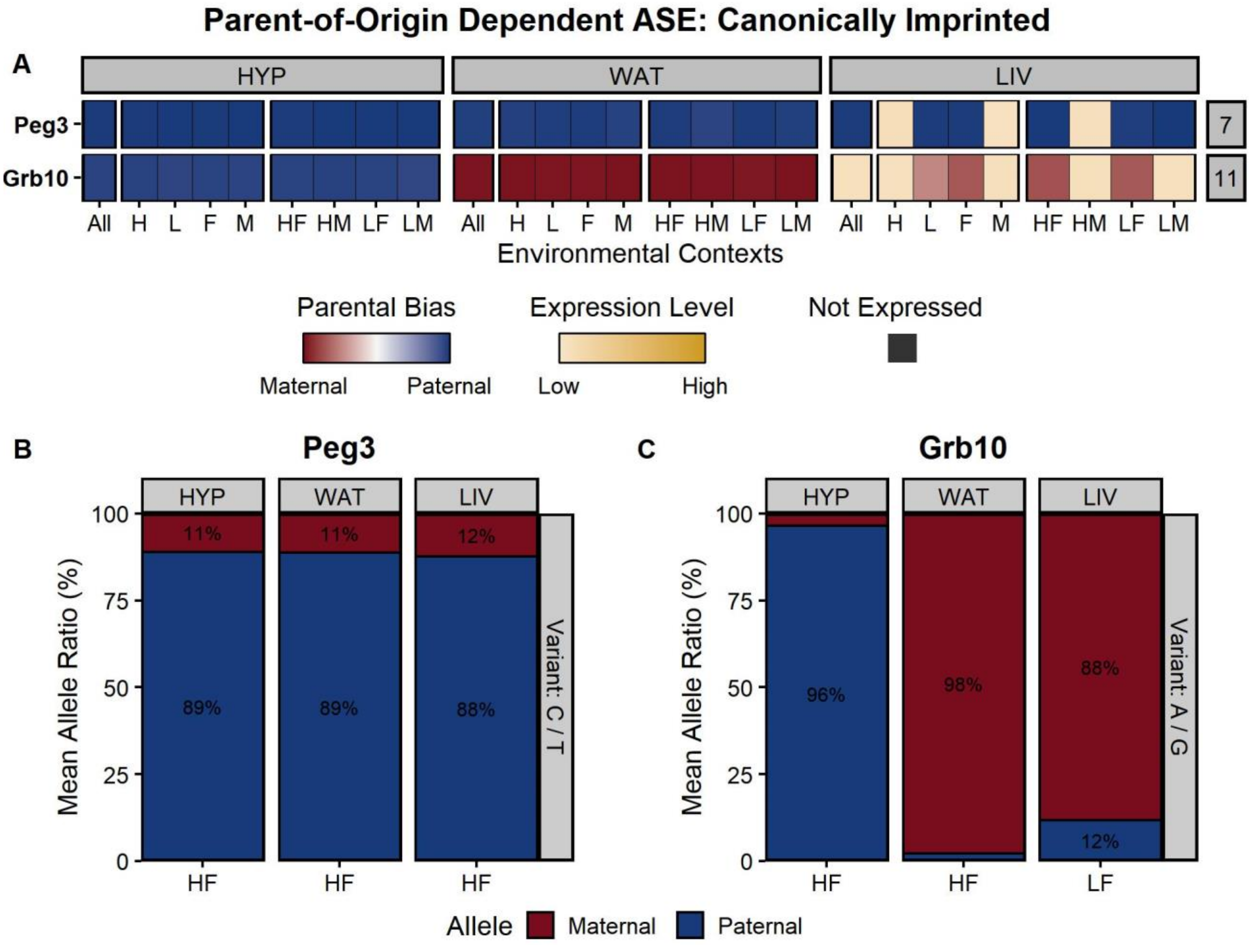

**Figure S8: Pyrosequencing validation of two canonically imprinted genes with parental biases.** *Peg3* is a tissue-independent imprinted gene with a paternal bias wherever it's expressed, regardless of environment. *Grb10* is a tissue-dependent imprinted gene with a paternal bias in HYP but a maternal bias in WAT and LIV. **(A)** Heatmap of the *Peg3* and *Grb10* expression profiles based on RNA-Seq data. Genes are color-coded by their expression pattern in each tissue-by-context analysis. Shades of red and blue indicate their degree of maternal or paternal bias, respectively (POE scores). Where genes are not significantly biased, shades of yellow indicate their biallelic expression levels (log-transformed total counts). Black indicates genes are not expressed in that analysis. Bolded genes are canonically imprinted. The y-axis is sorted by chromosomal position. Each supercolumn denotes a tissue: hypothalamus (HYP), white adipose (WAT), and liver (LIV). Each subcolumn denotes an environmental context: all contexts collapsed (All), high fat diet (H), low fat diet (L), females (F), males (M), high fat-fed females (HF), high fat-fed males (HM), low fat-fed females (LF), and low fat-fed males (LM). Pyrosequencing results for **(B)** *Peg3* and **(C)** *Grb10*. We quantified the mean allelic ratios in select tissue-by-context cohorts to validate their RNA-Seq expression profiles: *Peg3* is paternally biased in all three tissues, and *Grb10* exhibits a tissue-dependent switch in bias direction (paternally biased in the brain, maternally biased elsewhere). The y-axis shows the mean proportions (%) of alleles across biological replicates. Alleles are color-coded by their parental inheritance (maternal = red, paternal = blue). The x-axis is grouped by tissue, then ordered by environmental context.

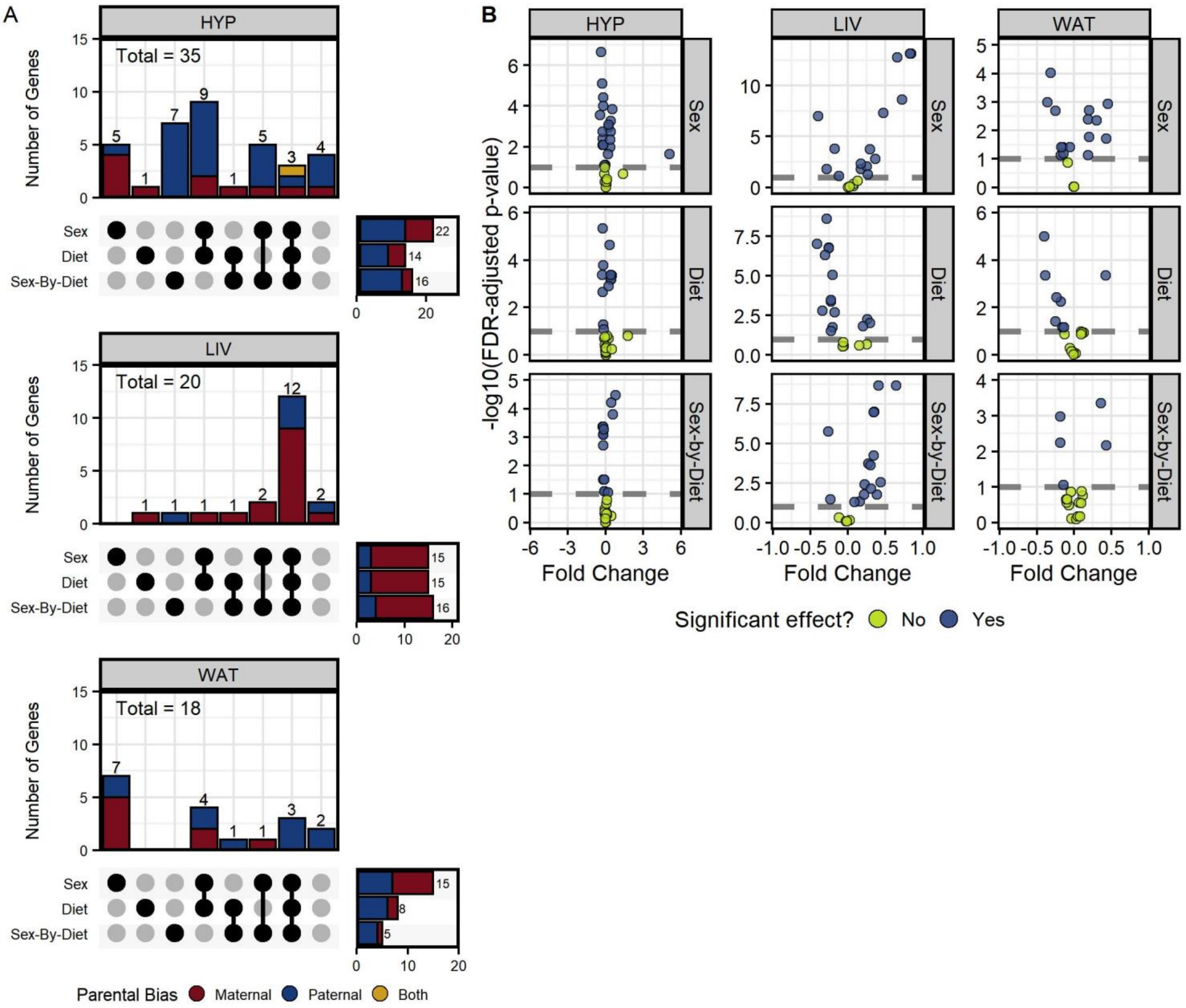

**Figure S9: Context-dependent genes with parental biases have significant sex and/or diet effects.**
For all context-dependent genes with parent-of-origin dependent ASE, we calculated individual POE scores and assessed how they vary across diet and/or sex contexts with ANOVA models. **(A)** UpSet plots for each tissue summarizing the set intersections of context-dependent genes with significant sex, diet, and/or sex-by-diet interaction effects (FDR-adjusted p-values ≤ 0.1). Most context-dependent genes were significant for more than one effect. The columns in the combination matrix (x-axis) correspond to the eight possible intersections of the three sets (sex, diet, and sex-by-diet effects), including where genes with context-dependent expression profiles do not have a significant effect. Stacked bars (y-axis) indicate the number of genes in each set intersection (also labeled above the bars) and are color-coded by the genes' expression bias directions (maternal = red, paternal = blue, both/direction-switching = yellow). The set menu on the right shows the total number of significant genes with each effect. The total number of context-dependent genes per tissue is labeled in the upper-left corner. **(B)** Volcano plots visualizing the ANOVA results for each model term/effect in each tissue's analysis. The y-axis shows the -$\log_{10}$(FDR-adjusted p-value) and the x-axis shows the fold change in mean individual POE scores between that context's cohorts (sex: females versus males, diet: high fat versus low fat, diet-by-sex: the cohorts with the greatest difference in mean POE scores). Dashed lines indicate significance thresholds. Genes are color-coded by whether they are significant for that effect (blue = yes, lime green = no). Columns are grouped by tissue and rows are grouped by model term/effect.

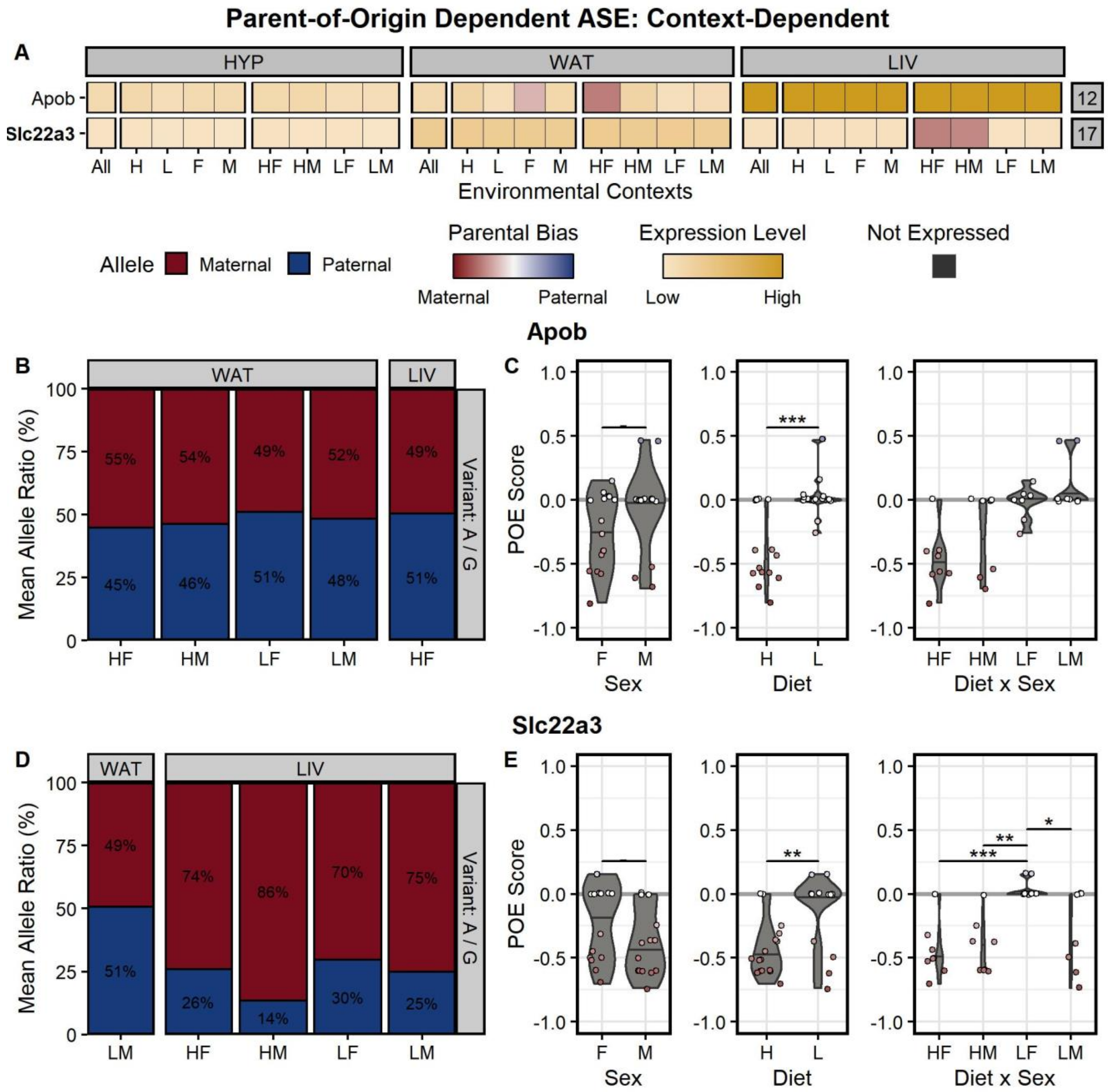

**Figure S10: Pyrosequencing validation of two context-dependent genes with parental biases.** *Apob* and *Slc22a3* are context-dependent genes with parental biases only in certain environmental contexts, but are biallelically expressed (no bias) elsewhere. **(A)** Heatmap of the *Apob* and *Slc22a3* expression profiles based on RNA-Seq data. Genes are color-coded by their expression pattern in each tissue-by-context analysis. Shades of red and blue indicate their degree of maternal or paternal bias, respectively (POE scores). Where genes are not significantly biased, shades of yellow indicate their biallelic expression levels (log-transformed total counts). Black indicates genes are not expressed in that analysis. Bolded genes are canonically imprinted. The y-axis is sorted by chromosomal position. The x-axis is grouped by tissue, then organized by environmental context (see **Figure S8**). **(B)** Pyrosequencing results and **(C)** ANOVA analyses confirm that *Apob* has a diet-specific maternal bias in the HF and HM contexts in WAT. **(D)** Pyrosequencing results and **(E)** ANOVA analyses confirm that *Slc22a3* has a diet- and diet-by-sex-specific maternal bias in the HF and HM contexts in LIV. **(Panels B, D)** We quantified the mean allelic ratios in select tissue-by-context cohorts with pyrosequencing to validate their RNA-Seq expression profiles. The y-axis shows the mean proportions (%) of alleles across biological replicates. Alleles are color-coded by their parental inheritance (maternal = red, paternal = blue). The x-axis is grouped by tissue, then ordered by environmental context. **(Panels C, E)** We modeled how individual POE scores (y-axis) vary across diet, sex, and diet-by-sex contexts (x-axis). Violin plots display the 50$^{th}$ quantiles (horizontal bars) of each context-dependent cohort. Individual data points are color-coded by their parental bias (POE scores). - p ≤ 0.1, * p ≤ 0.05, ** p ≤ 0.01, *** p ≤ 0.001; assessed by ANOVA (sex, diet) or Tukey's post-hoc tests (diet-by-sex).

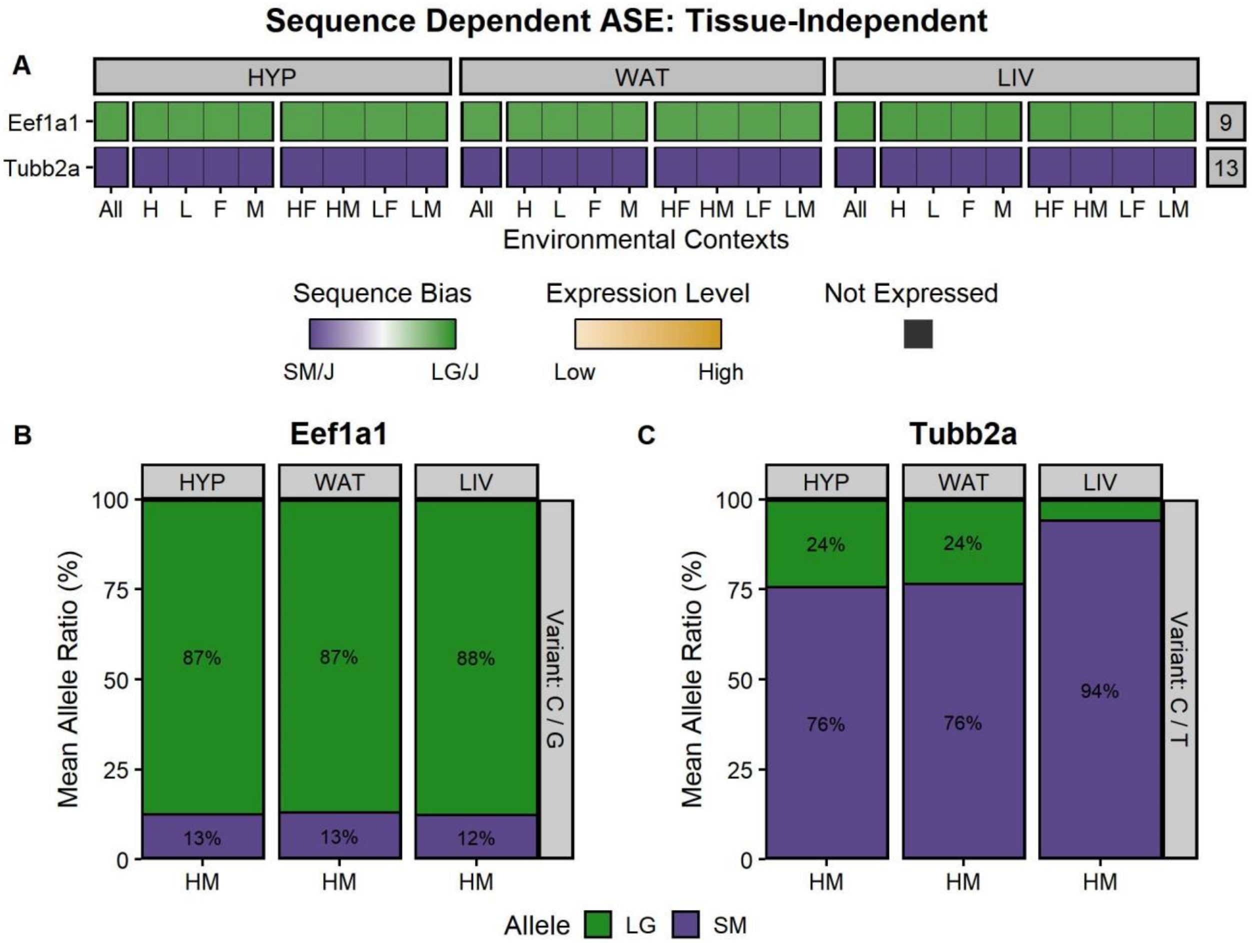

**Figure S11: Pyrosequencing validation of two tissue-independent genes with sequence biases.** *Eef1a1* and *Tubb2a* are tissue-independent genes with a LG/J and SM/J allelic bias (respectively) wherever they are expressed, regardless of environmental context. **(A)** Heatmap of the *Eef1a1* and *Tubb2a* expression profiles based on RNA-Seq data. Genes are color-coded by their expression pattern in each tissue-by-context analysis. Shades of purple and green indicate their degree of SM/J or LG/J allelic bias, respectively (AGE scores). Where genes are not significantly biased, shades of yellow indicate their biallelic expression levels (log-transformed total counts). Black indicates genes are not expressed in that analysis. The y-axis is sorted by chromosomal position. Each supercolumn denotes a tissue: hypothalamus (HYP), white adipose (WAT), and liver (LIV). Each subcolumn denotes an environmental context: all contexts collapsed (All), high fat diet (H), low fat diet (L), females (F), males (M), high fat-fed females (HF), high fat-fed males (HM), low fat-fed females (LF), and low fat-fed males (LM). Pyrosequencing results for **(B)** *Eef1a1* and **(C)** *Tubb2a*. We quantified the mean allelic ratios in select tissue-by-context cohorts to validate their RNA-Seq expression profiles: *Eef1a1* is LG/J biased in all three tissues and *Tubb2a* is SM/J biased in all three tissues. The y-axis shows the mean proportions (%) of alleles across biological replicates. Alleles are color-coded by their strain-specific haplotype (SM/J = purple, LG/J = green). The x-axis is grouped by tissue, then ordered by environmental context.

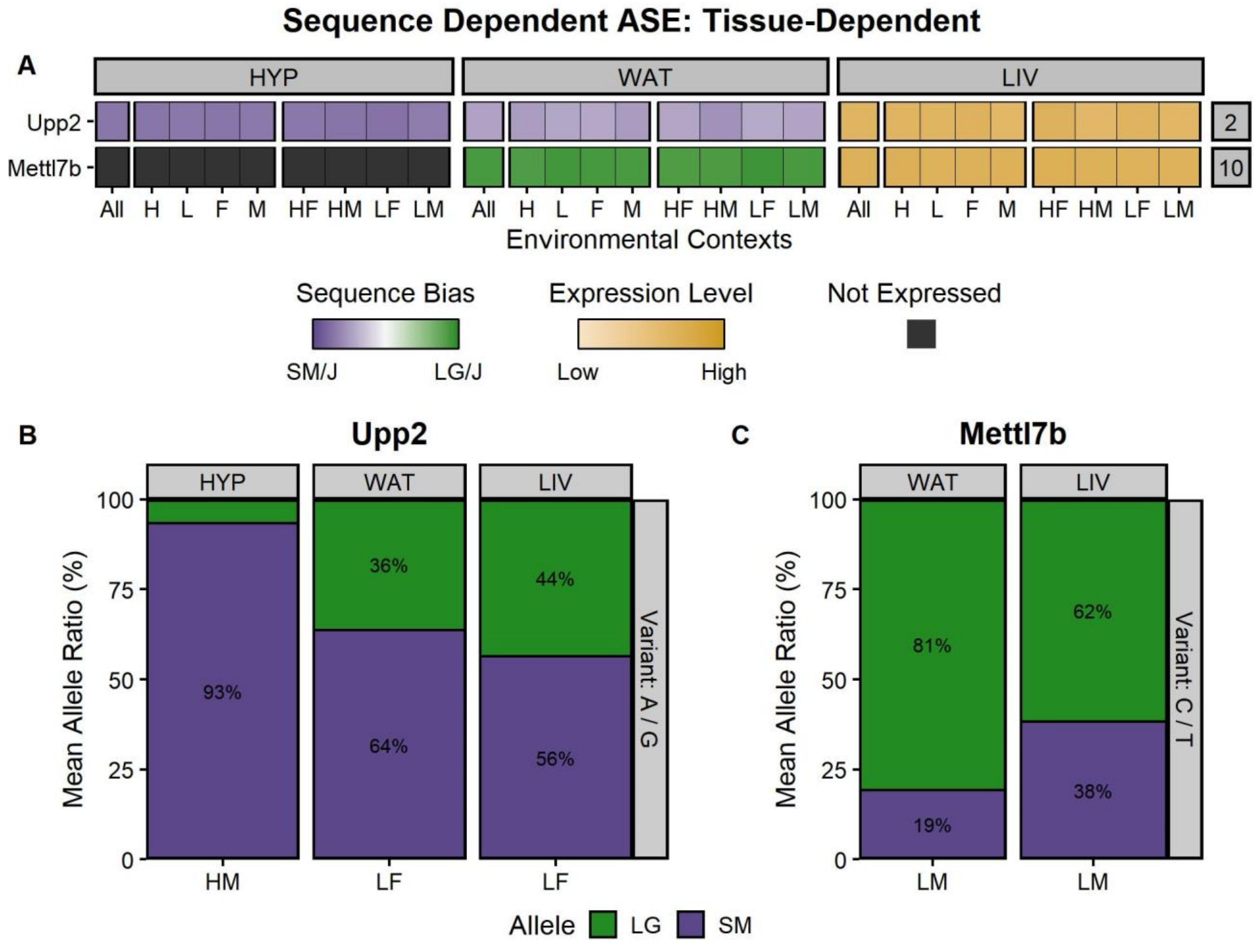

**Figure S12: Pyrosequencing validation of two tissue-dependent genes with sequence biases.** *Upp2* is a tissue-dependent gene with a strong SM/J bias in HYP, a moderate SM/J bias in WAT, and no bias in LIV. *Mettl7b* is a tissue-dependent gene with a LG/J bias in WAT, no bias in LIV, and is not expressed in HYP. **(A)** Heatmap of the *Upp2* and *Mettl7b* expression profiles based on RNA-Seq data. Genes are color-coded by their expression pattern in each tissue-by-context analysis. Shades of purple and green indicate their degree of SM/J or LG/J allelic bias, respectively (AGE scores). Where genes are not significantly biased, shades of yellow indicate their biallelic expression levels (log-transformed total counts). Black indicates genes are not expressed in that analysis. The y-axis is sorted by chromosomal position. Each supercolumn denotes a tissue: hypothalamus (HYP), white adipose (WAT), and liver (LIV). Each subcolumn denotes an environmental context: all contexts collapsed (All), high fat diet (H), low fat diet (L), females (F), males (M), high fat-fed females (HF), high fat-fed males (HM), low fat-fed females (LF), and low fat-fed males (LM). Pyrosequencing results for **(B)** *Upp2* and **(C)** *Mettl7b*. We quantified the mean allelic ratios in select tissue-by-context cohorts to validate their RNA-Seq expression profiles. *Upp2* has an extreme SM/J bias in HYP, a moderate SM/J bias in WAT, and a weak SM/J bias in LIV. *Mettl7b* has a strong LG/J bias in WAT and a weaker LG/J bias in LIV. The y-axis shows the mean proportions (%) of alleles across biological replicates. Alleles are color-coded by their strain-specific haplotype (SM/J = purple, LG/J = green). The x-axis is grouped by tissue, then ordered by environmental context.

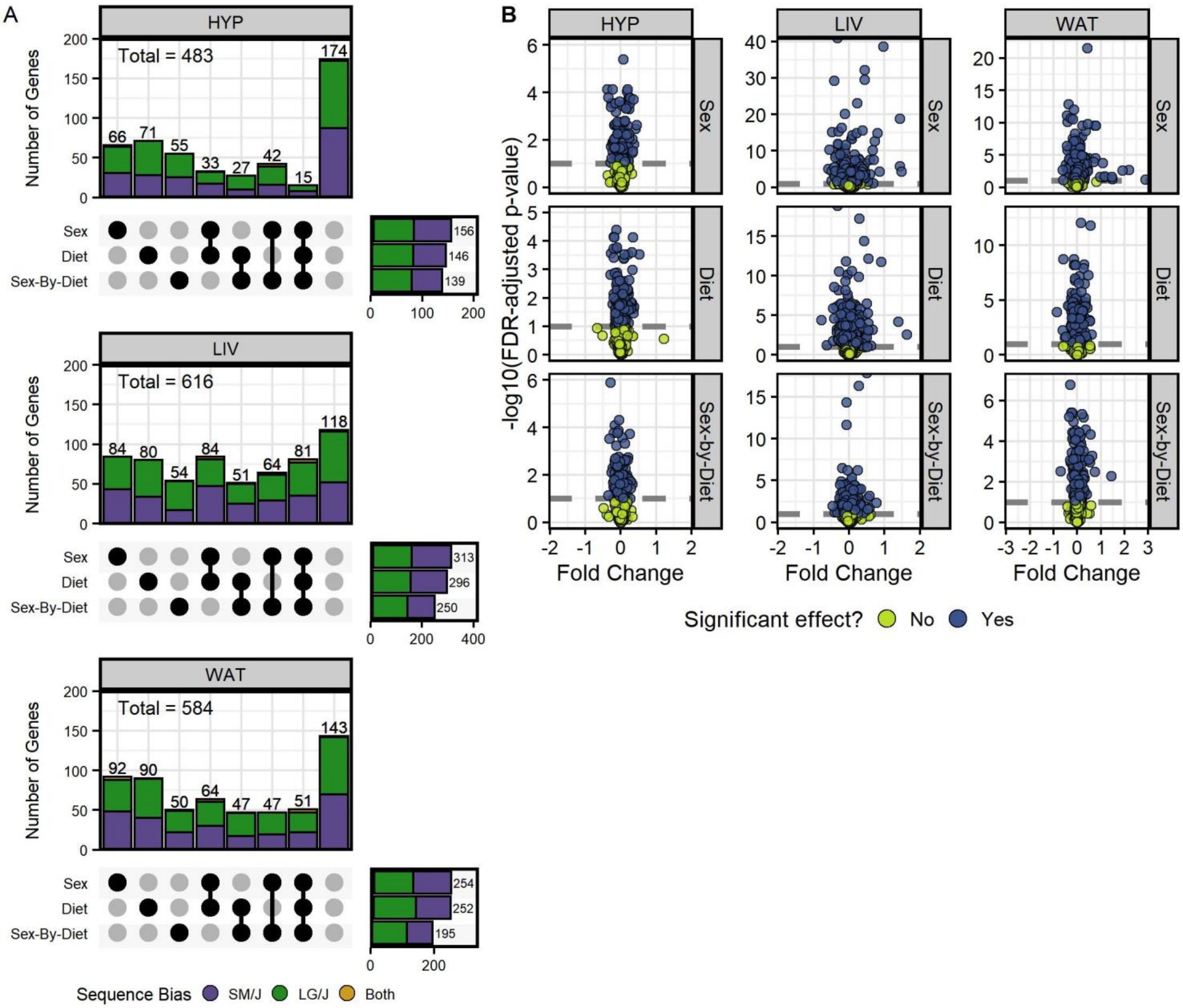

**Figure S13: Context-dependent genes with sequence biases have significant sex and/or diet effects.** For all context-dependent genes with sequence dependent ASE, we calculated individual AGE scores and assessed how they vary across diet and/or sex contexts with ANOVA models. **(A)** UpSet plots for each tissue summarizing the set intersections of context-dependent genes with significant sex, diet, and/or sex-by-diet interaction effects (FDR-adjusted p-values ≤ 0.1). Most context-dependent genes were significant for more than one effect. The columns in the combination matrix (x-axis) correspond to the eight possible intersections of the three sets (sex, diet, and sex-by-diet effects), including where genes with context-dependent expression profiles do not have a significant effect. Stacked bars (y-axis) indicate the number of genes in each set intersection (also labeled above the bars) and are color-coded by the genes' expression bias directions (SM/J = purple, LG/J = green, both/direction-switching = yellow). The set menu on the right shows the total number of significant genes with each effect. The total number of context-dependent genes per tissue is labeled in the upper-left corner. **(B)** Volcano plots visualizing the ANOVA results for each model term/effect in each tissue's analysis. The y-axis shows the -$\log_{10}$(FDR-adjusted p-value) and the x-axis shows the fold change in mean individual AGE scores between that context's cohorts (sex: females versus males, diet: high fat versus low fat, diet-by-sex: the cohorts with the greatest difference in mean AGE scores). Dashed lines indicate significance thresholds. Genes are color-coded by whether they are significant for that effect (blue = yes, lime green = no). Columns are grouped by tissue and rows are grouped by model term/effect.

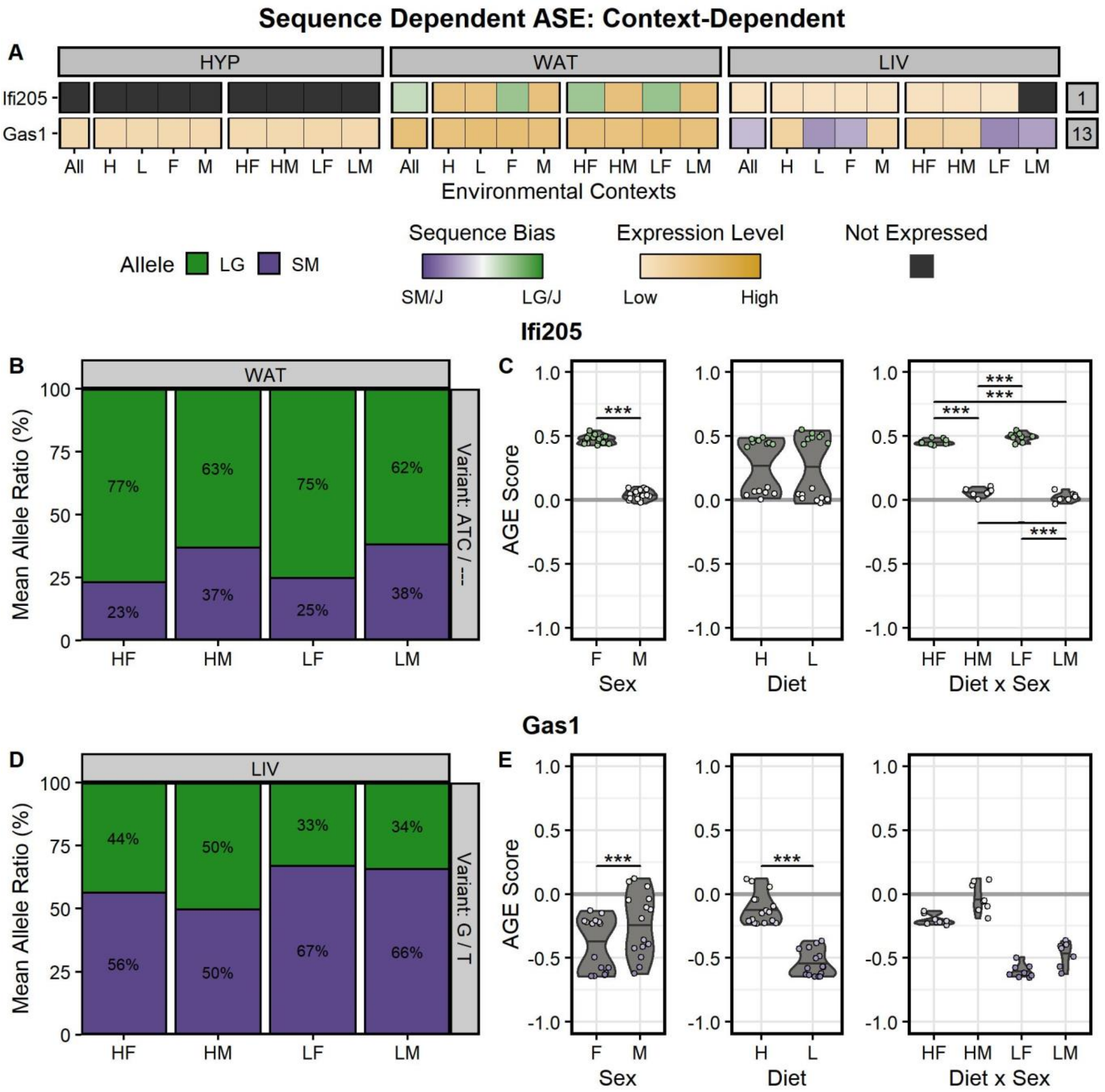

**Figure S14: Pyrosequencing validation of two context-dependent genes with sequence biases.** *Ifi205* and *Gas1* are context-dependent genes with sequence biases only in certain environmental contexts, but are biallelically expressed (no bias) elsewhere. **(A)** Heatmap of the *Ifi205* and *Gas1* expression profiles based on RNA-Seq data. Genes are color-coded by their expression pattern in each tissue-by-context analysis. Shades of purple and green indicate their degree of SM/J or LG/J allelic bias, respectively (AGE scores). Where genes are not significantly biased, shades of yellow indicate their biallelic expression levels (log-transformed total counts). Black indicates genes are not expressed in that analysis. The y-axis is sorted by chromosomal position. The x-axis is grouped by tissue, then organized by environmental context (see **Figure S12**). **(B)** Pyrosequencing results and **(C)** ANOVA analyses confirm that *Ifi205* has a sex- and diet-by-sex-specific LG/J bias in the HF and LF contexts in WAT. **(D)** Pyrosequencing results and **(E)** ANOVA analyses confirm that *Gas1* has a diet and sex-specific SM/J bias in the LF and LM contexts in LIV. **(Panels B, D)** We quantified the mean allelic ratios in select tissue-by-context cohorts with pyrosequencing to validate their RNA-Seq expression profiles. The y-axis shows the mean proportions (%) of alleles across biological replicates. Alleles are color-coded by their strain-specific haplotype (SM/J = purple, LG/J = green). The x-axis is grouped by tissue, then ordered by environmental context. **(Panels C, E)** We modeled how individual AGE scores (y-axis) vary across diet, sex, and diet-by-sex contexts (x-axis). Violin plots display the 50$^{th}$ quantiles (horizontal bars) of each context-dependent cohort. Individual data points are color-coded by their sequence bias (AGE scores). - $p \leq 0.1$, * $p \leq 0.05$, ** $p \leq 0.01$, *** $p \leq 0.001$; assessed by ANOVA (sex, diet) or Tukey's post-hoc tests (diet-by-sex).

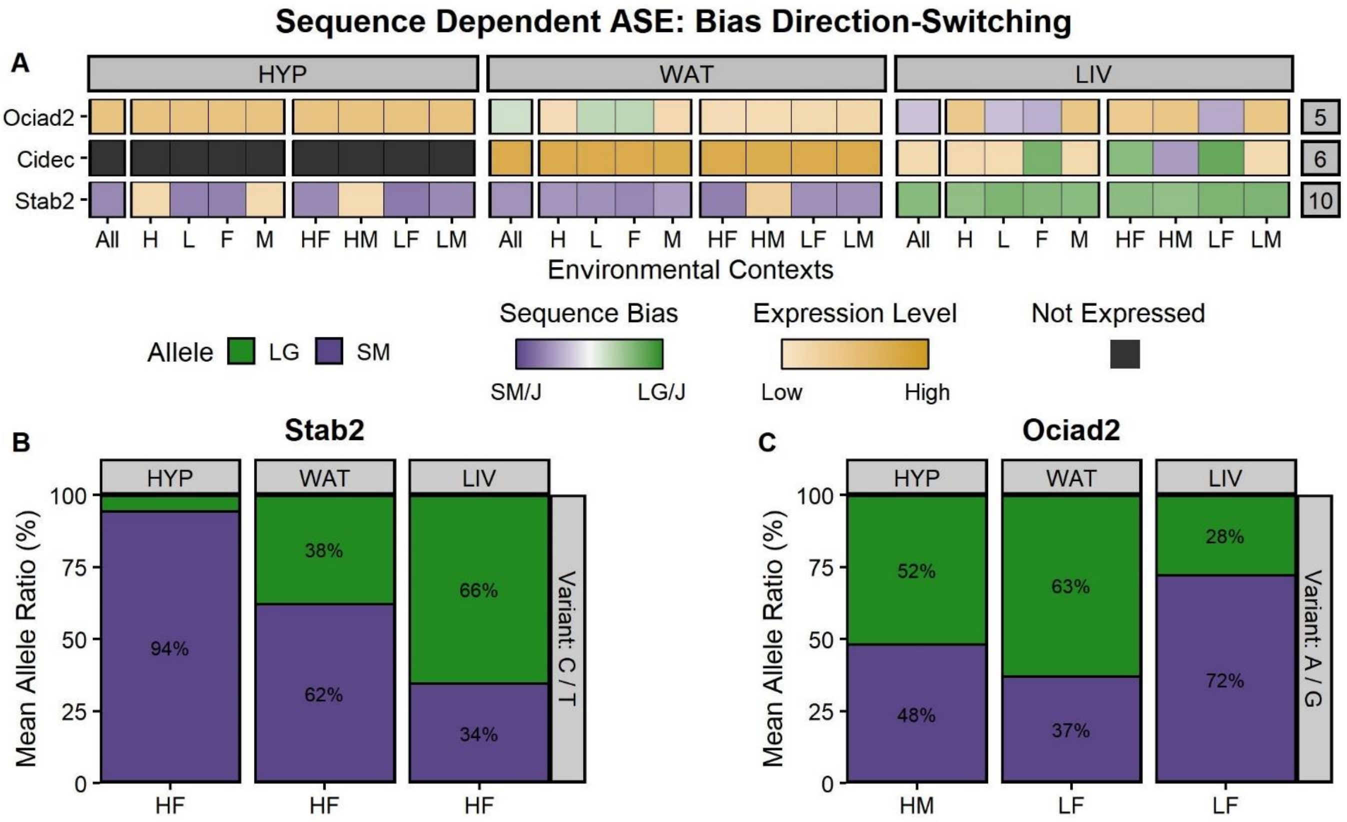

**Figure S15: Pyrosequencing validation of two genes with sequence bias that switch directions.** *Stab2* and *Ociad2* have sequence biases in opposite directions across their tissues-by-context analyses. This direction-switching occurred at the tissue level. **(A)** Heatmap of the *Stab2* and *Ociad2* expression profiles based on RNA-Seq data. Genes are color-coded by their expression pattern in each tissue-by-context analysis. Shades of purple and green indicate their degree of SM/J or LG/J allelic bias, respectively (AGE scores). Where genes are not significantly biased, shades of yellow indicate their biallelic expression levels (log-transformed total counts). Black indicates genes are not expressed in that analysis. The y-axis is sorted by chromosomal position. Each supercolumn denotes a tissue: hypothalamus (HYP), white adipose (WAT), and liver (LIV). Each subcolumn denotes an environmental context: all contexts collapsed (All), high fat diet (H), low fat diet (L), females (F), males (M), high fat-fed females (HF), high fat-fed males (HM), low fat-fed females (LF), and low fat-fed males (LM). Pyrosequencing results for **(B)** *Stab2* and **(C)** *Ociad2*. We quantified the mean allelic ratios in select tissue-by-context cohorts to validate their RNA-Seq expression profiles. *Stab2* has an extreme SM/J bias in HYP, a moderate SM/J bias in WAT, but a moderate LG/J bias in LIV. *Ociad2* has no bias (biallelic) in HYP, a moderate LG/J bias in certain contexts in WAT, and a strong SM/J bias in certain contexts in LIV. The y-axis shows the mean proportions (%) of alleles across biological replicates. Alleles are color-coded by their strain-specific haplotype (SM/J = purple, LG/J = green). The x-axis is grouped by tissue, then ordered by environmental context.

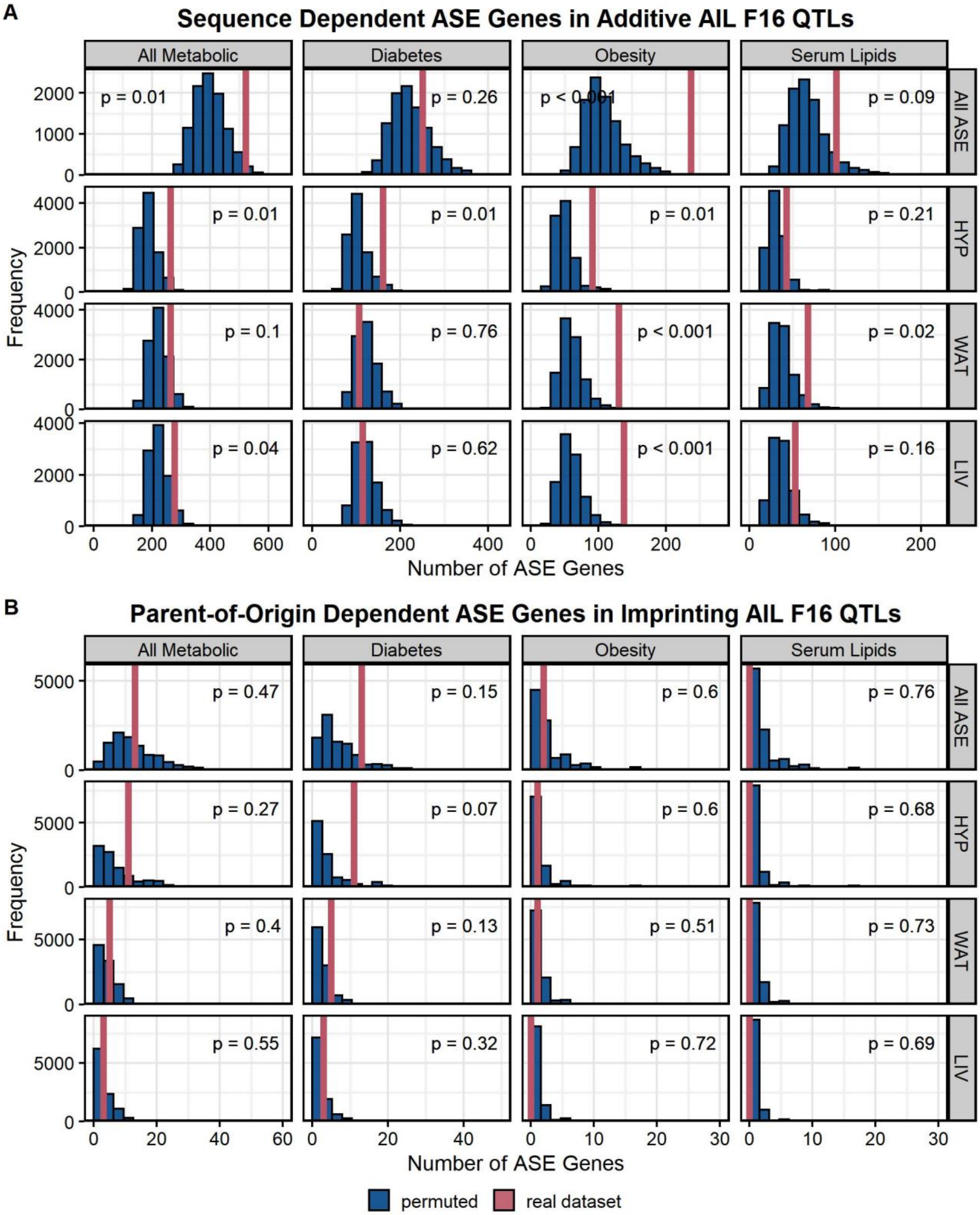

**Figure S16: Enrichment results between ASE genes and AIL $F_{16}$ QTLs.**
Histograms of enrichment results between **(A)** sequence dependent ASE genes in additive AIL $F_{16}$ QTL and **(B)** parent-of-origin dependent ASE genes in imprinting AIL $F_{16}$ QTL. Columns are grouped by trait-specific QTL sets and rows are grouped by tissue-specific ASE gene sets ("all ASE" = all tissues collapsed together). The x-axis denotes the number of ASE genes overlapping with all AIL QTL in that set. The y-axis denotes the number of permutation iterations with each overlap count. The null model distribution is colored in blue. The real overlap count for each ASE/QTL comparison is indicated by the pink line. The enrichment p-values are annotated for each analysis; $p \leq 0.05$ was considered significant.

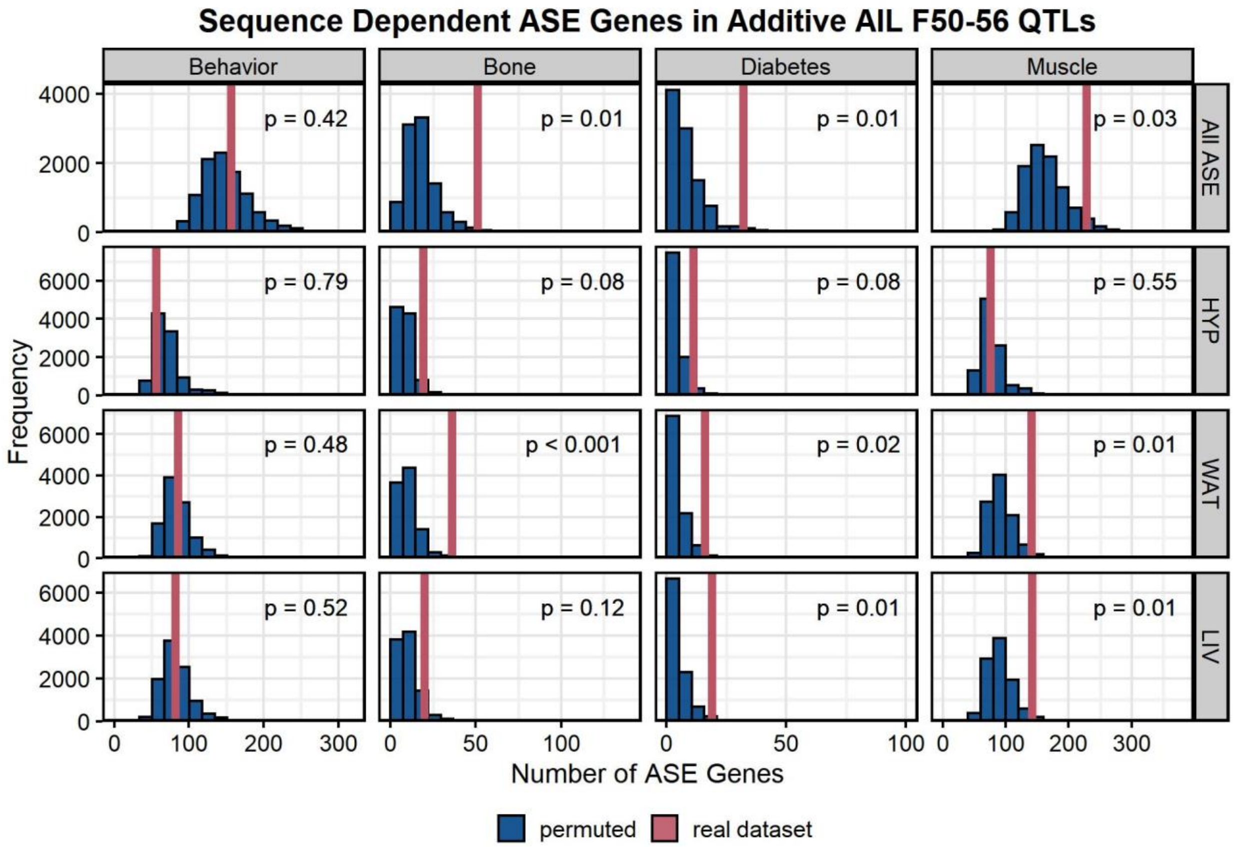

**Figure S17: Enrichment results between ASE genes and AIL $F_{50\text{-}56}$ QTLs.**
Histograms of enrichment results between sequence dependent ASE genes in additive AIL $F_{50\text{-}56}$ QTL. Columns are grouped by trait-specific QTL sets and rows are grouped by tissue-specific ASE gene sets (“all ASE” = all tissues collapsed together). The x-axis denotes the number of ASE genes overlapping with all AIL QTL in that set. The y-axis denotes the number of permutation iterations with each overlap count. The null model distribution is colored in blue. The real overlap count for each ASE/QTL comparison is indicated by the pink line. The enrichment p-values are annotated for each analysis; $p \leq 0.05$ was considered significant.

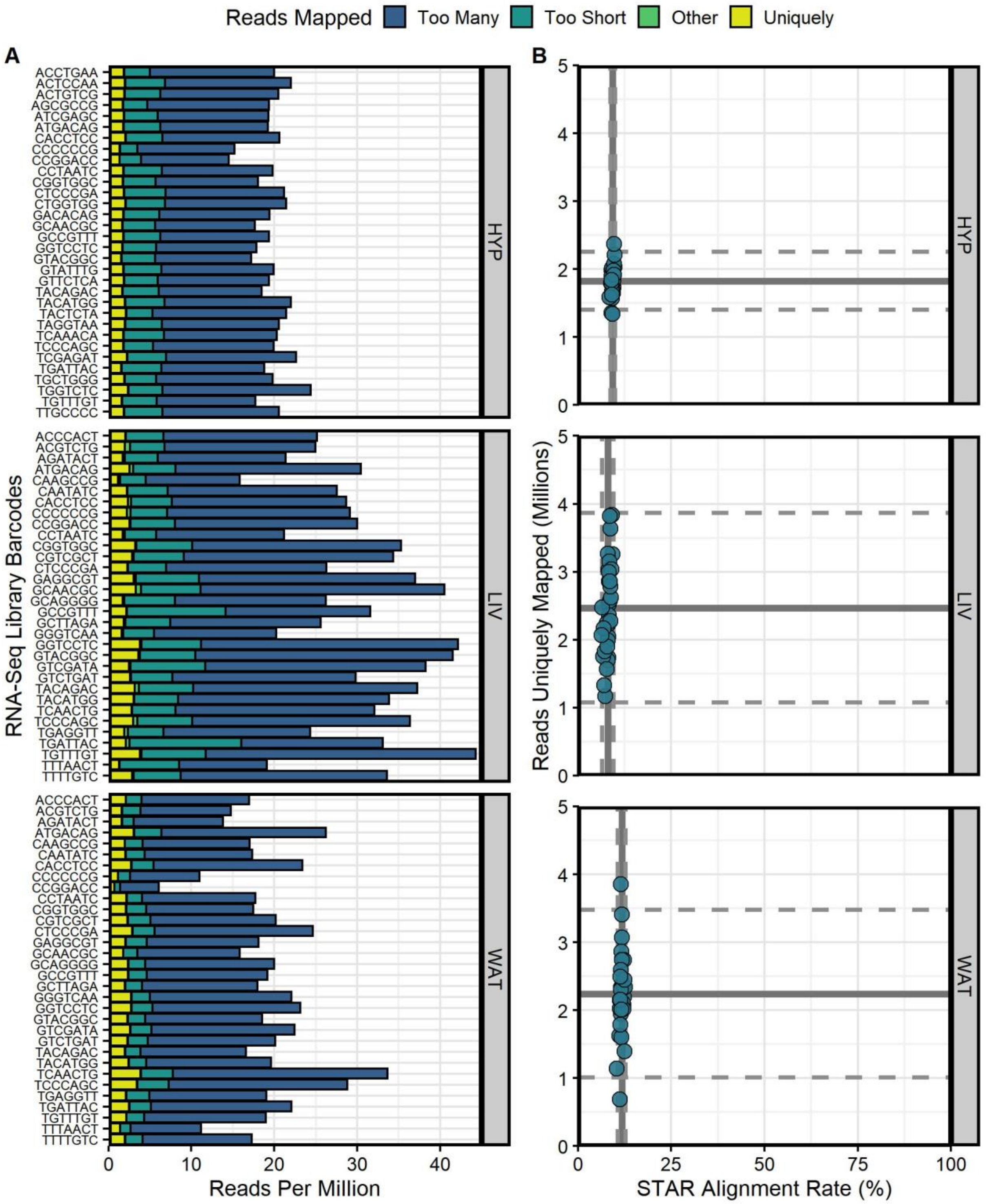

**Figure S18: STAR alignment summaries for RNA-Seq libraries in each tissue.**
We sequenced 32 mice per tissue on an Illumina HiSeq 4000 at 100bp PE reads. **(A)** Bar graph for RNA-Seq libraries in each tissue (by barcode, y-axis) showing the number of reads (per million, x-axis) that mapped to too many loci (dark blue), were too short to map (teal), were not mapped for another reason (green), and mapped uniquely (yellow). “Too many loci” is defined as n > 1 since multi-mapping was disallowed for ASE calling. **(B)** Scatterplot for each tissue comparing the number of RNA-Seq reads uniquely mapped (in millions of reads, y-axis) against the proportion of reads mapped from total input reads (%, x-axis). Mean alignment rates and mean number of reads mapped are shown as solid lines; their two standard deviations are shown as dashed lines.

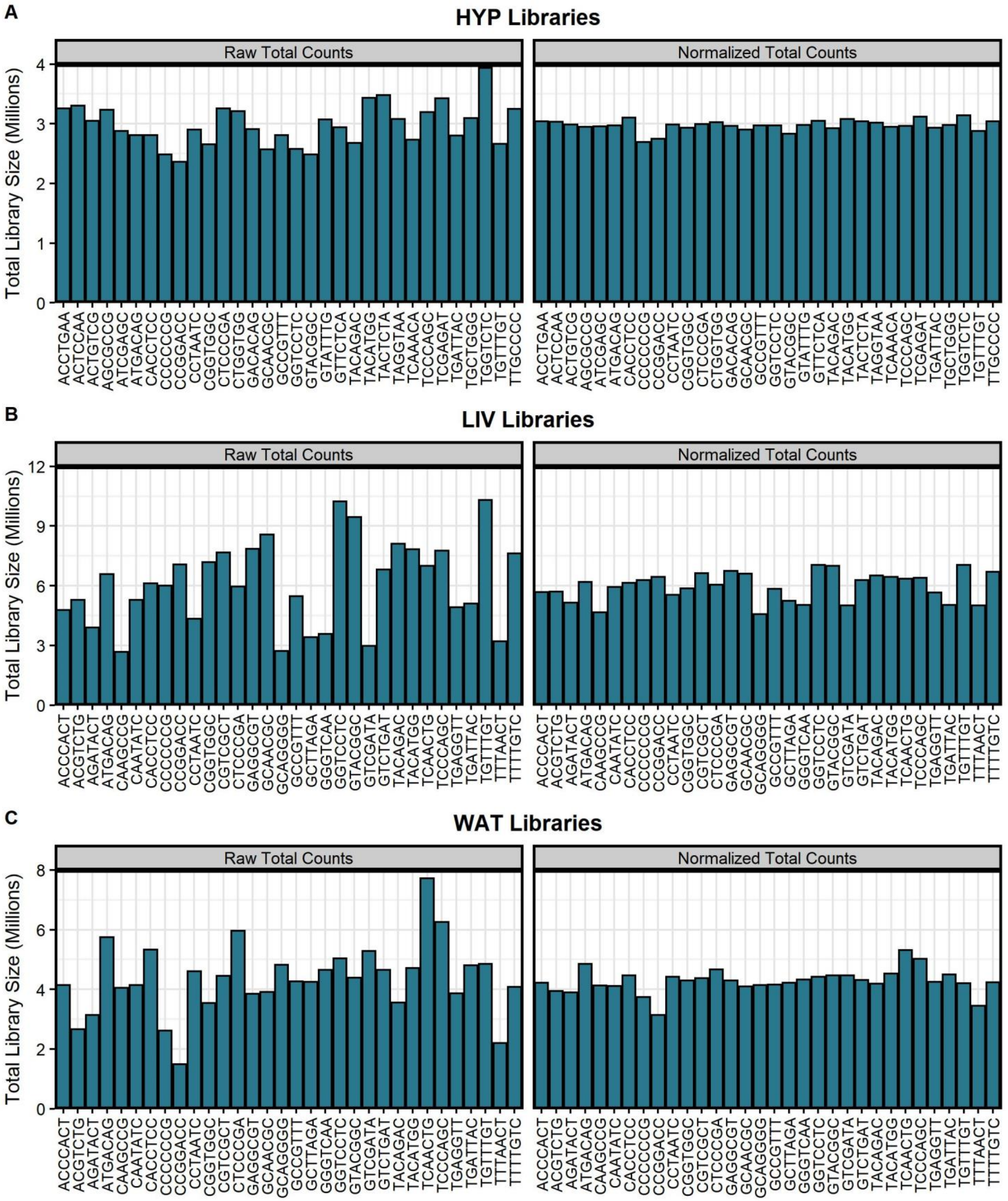

**Figure S19: Gene-level allele-specific read counts were upper quartile normalized.**
Total library sizes (in millions of reads) before and after upper quartile normalization for all libraries in **(A)** HYP, **(B)** LIV, and **(C)** WAT. Total library sizes were calculated by summing all gene-level allele-specific read counts. The y-axis shows the total library size for each RNA-Seq library (x-axis).

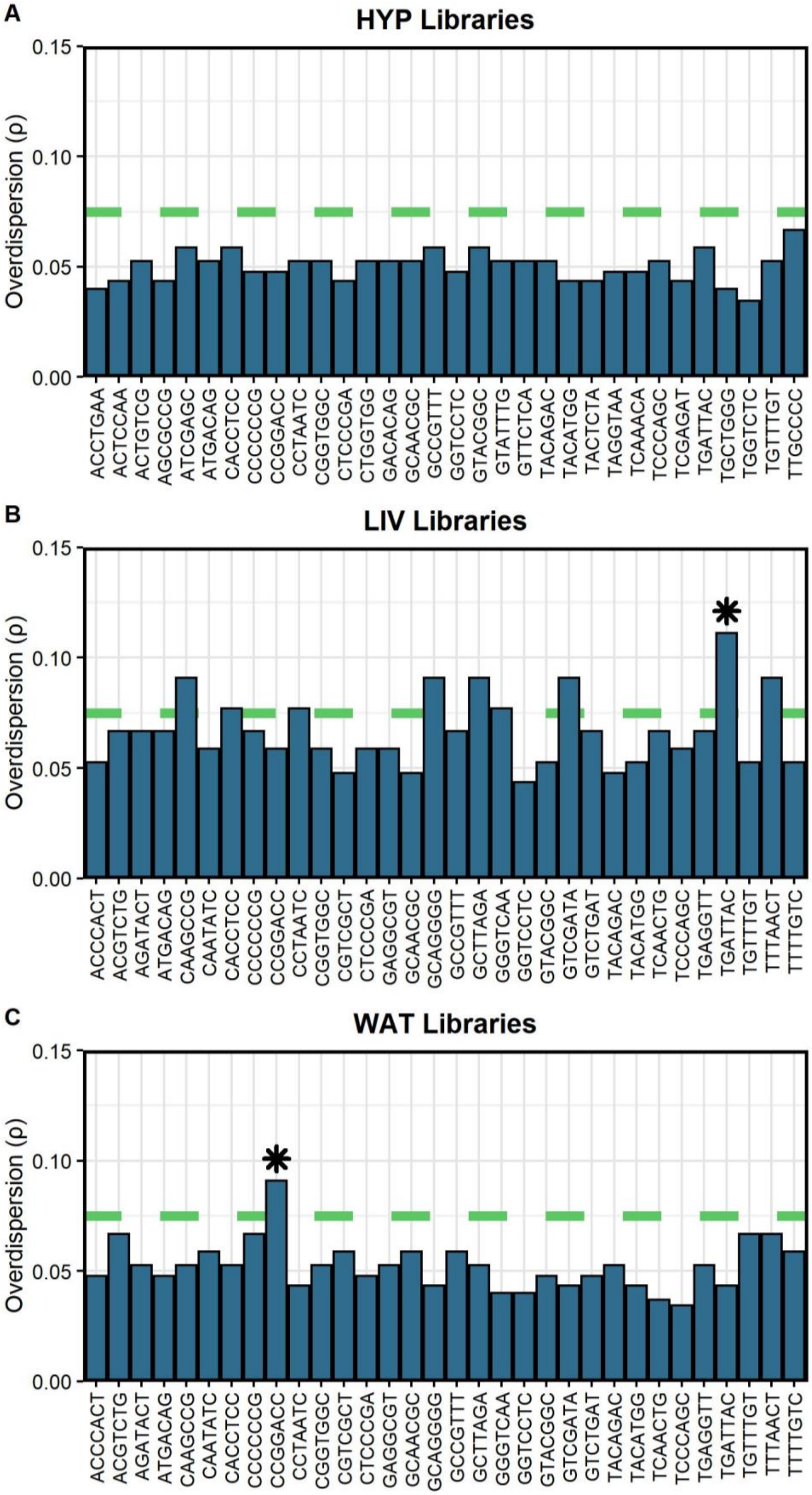

**Figure S20: RNA-Seq libraries in each tissue are sufficiently complex to detect ASE.**
Complexity results for all libraries in **(A)** HYP, **(B)** LIV, and **(C)** WAT. We measured each library's complexity by fitting its LG/J allele bias distribution to a beta-binomial distribution and calculating its overdispersion parameter (ρ). The y-axis shows the overdispersion values (ρ) for each RNA-Seq library (x-axis). Dashed green lines indicate complexity thresholds; lower values of ρ (< 0.075) signify a library is sufficiently complex. Two libraries were deemed to have poor complexity (flagged with a *) and were removed from further analyses

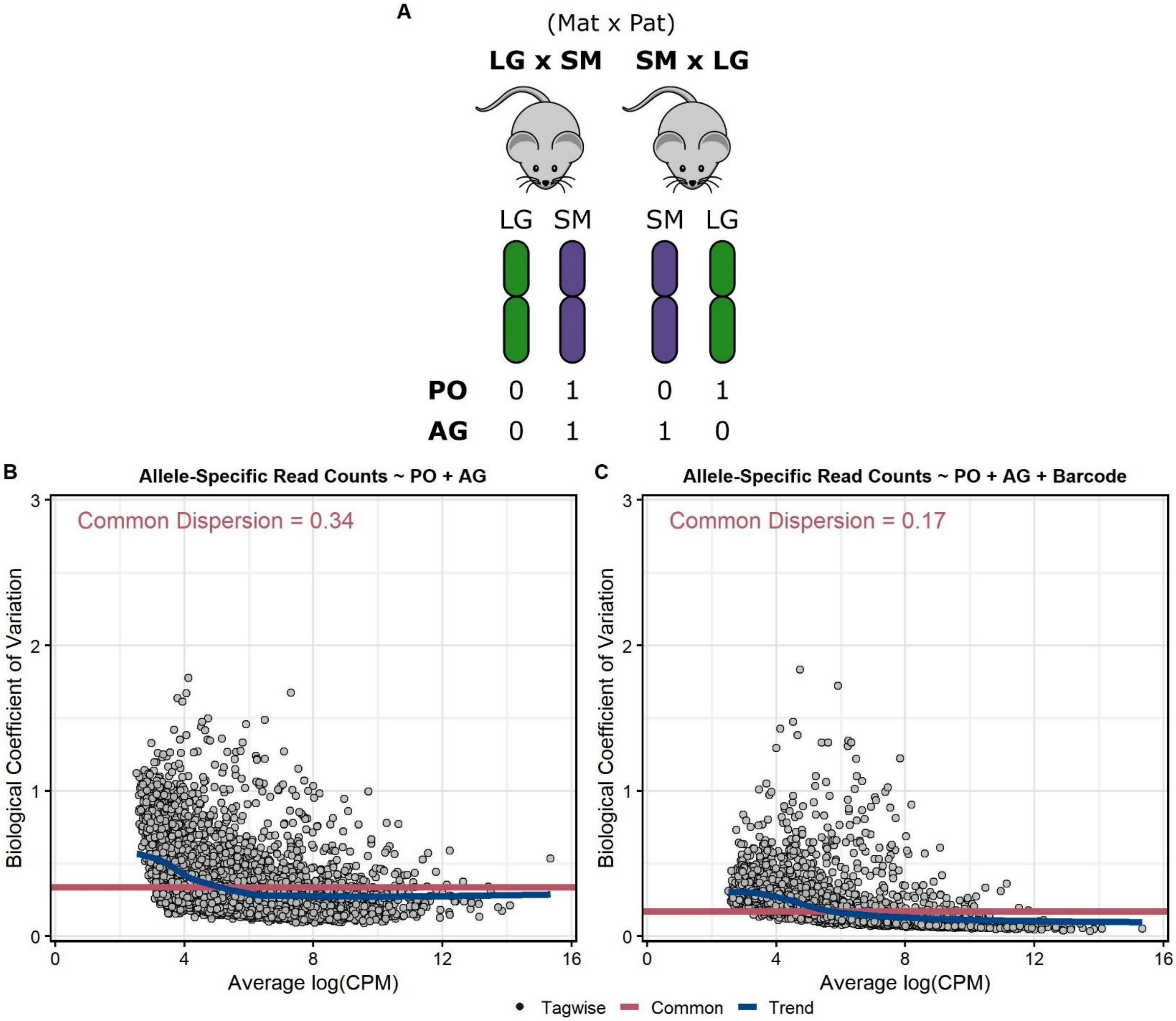

**Figure S21: Schematic of variables in the generalized linear model.**
**(A)** We quantified allele-specific expression in a $F_1$ reciprocal cross model of the LG/J and SM/J strains. Briefly, LG/J mothers were mated with SM/J fathers and vice versa, resulting in two crosses of $F_1$ offspring who are genetically equivalent but differ in the allelic direction of inheritance. We assigned two binary variables to the allele-specific read counts for the parent-of-origin (PO) and sequence/allelic genotype (AG) terms, denoting whether the allele was maternally or paternally derived (0 and 1 for PO term, respectively) and LG/J or SM/J haplotype derived (0 and 1 for AG term, respectively). Scatterplots measuring model fit for two potential negative binomial generalized linear models: **(B)** without the paired-sample design and **(C)** with the paired-sample design (i.e. an additional term representing the RNA-Seq library barcodes). The x-axis denotes the average gene abundance (in $\log_2$ counts per million). The y-axis denotes the gene-wise biological coefficient of variation, which is the square root of the tagwise dispersion. Dots represent individual genes. The blue line represents the dispersion trend. The red line represents the common dispersion across all genes. A lower common dispersion indicates a better model fit; the linear regression model implementing the paired-sample design has a better fit.

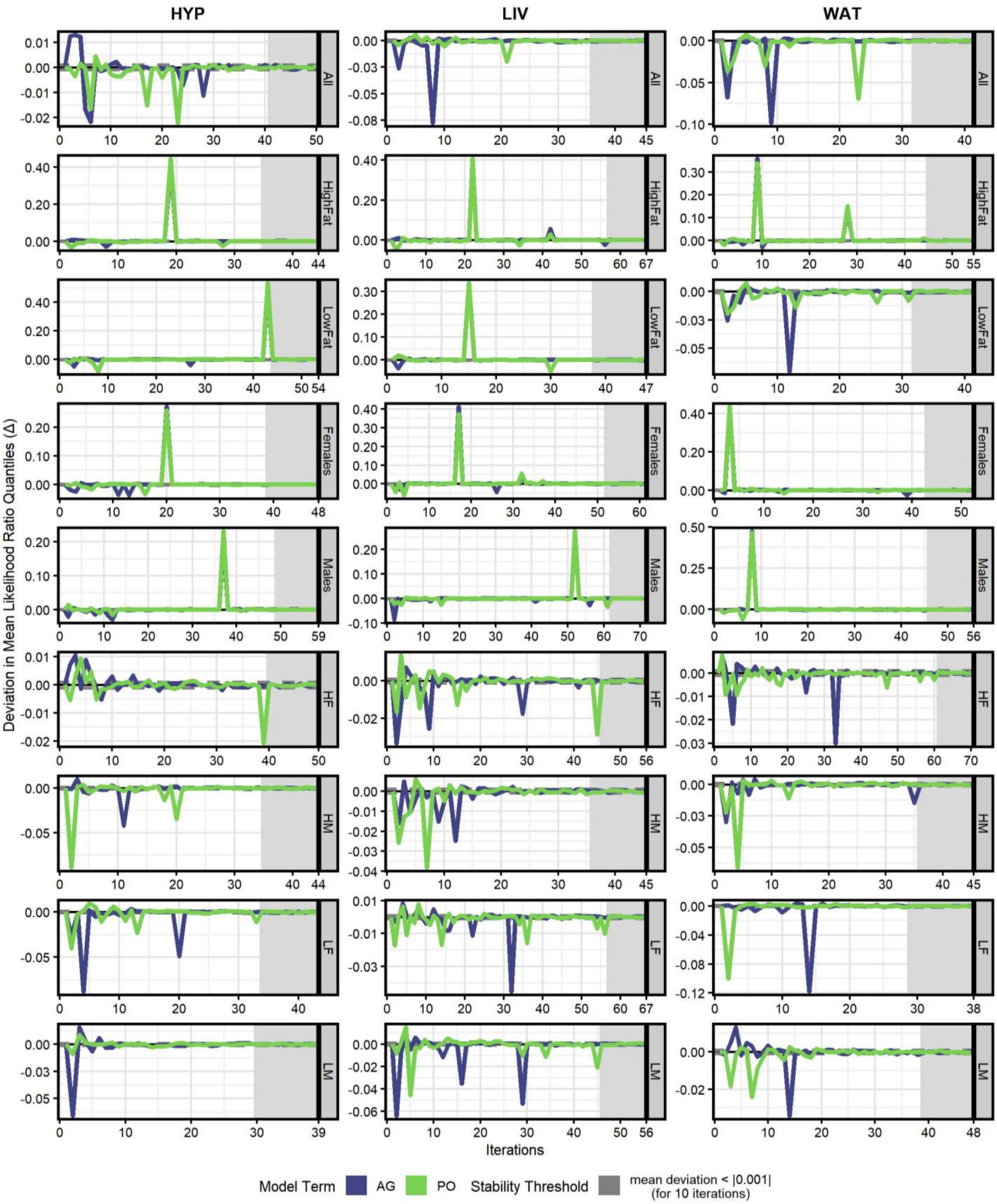

**Figure S22: Stable null distribution of likelihood ratios for both model terms generated by permutation.** In each tissue-by-context analysis, allele-specific read counts for all genes were randomly-shuffled and re-analyzed over several iterations to generate an appropriately null model. After each iteration, we calculated the change in mean likelihood ratio (LR) quantiles for both Allelic Genotype (AG, blue) and Parent-of-Origin (PO, green) terms in the full permuted dataset. We added new iterations until the null model was considered "stable": the mean LR quantiles for both terms fluctuated by <|0.001| (i.e. approached zero) for 10 consecutive iterations (shaded gray). Columns are grouped by tissue. Rows are grouped by environmental context. The y-axis shows the deviation (Δ) in mean LR quantiles; the x-axis shows the number of iterations (range = 38-71, mean = 51).

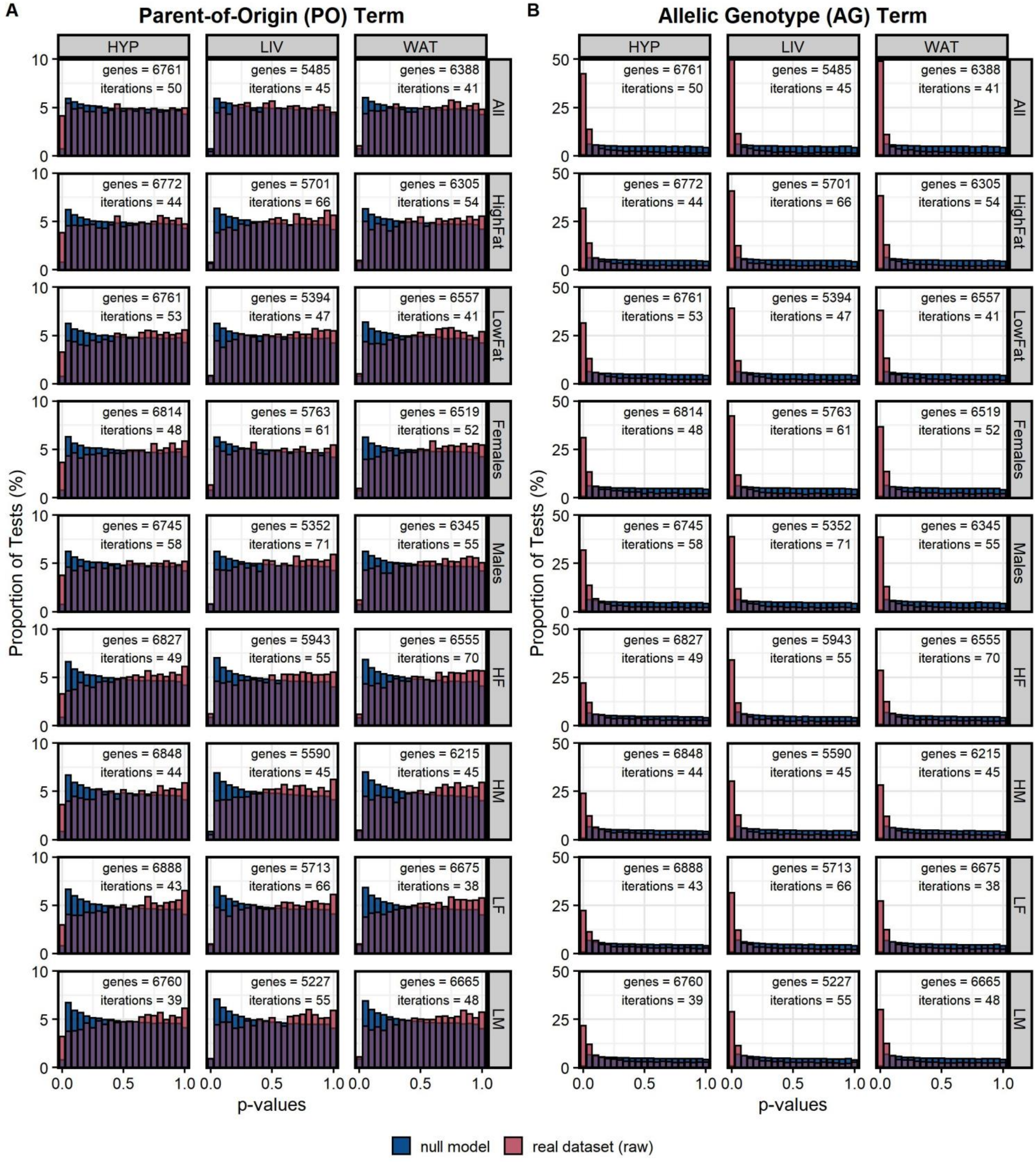

**Figure S23: P-value distributions between real and permuted datasets for both PO and AG terms.** Histograms of raw p-value distributions for the **(A)** Parent-of-Origin (PO) and **(B)** Allelic Genotype (AG) model terms in each tissue-by-context analysis. Distributions are overlaid to highlight how the permuted/null models (blue) differ from the real datasets (pink) before multiple tests correction: the null models have proportionally few tests (i.e. genes) with extremely small p-values compared to the real datasets. Purple bars indicate where the two distributions overlap. The x-axis shows the range of p-values (0-1) and the y-axis shows the proportion of tests (%) in each p-value bin. Columns are grouped by tissue and rows are grouped by environmental context. The number of genes and iterations are also denoted for each analysis.

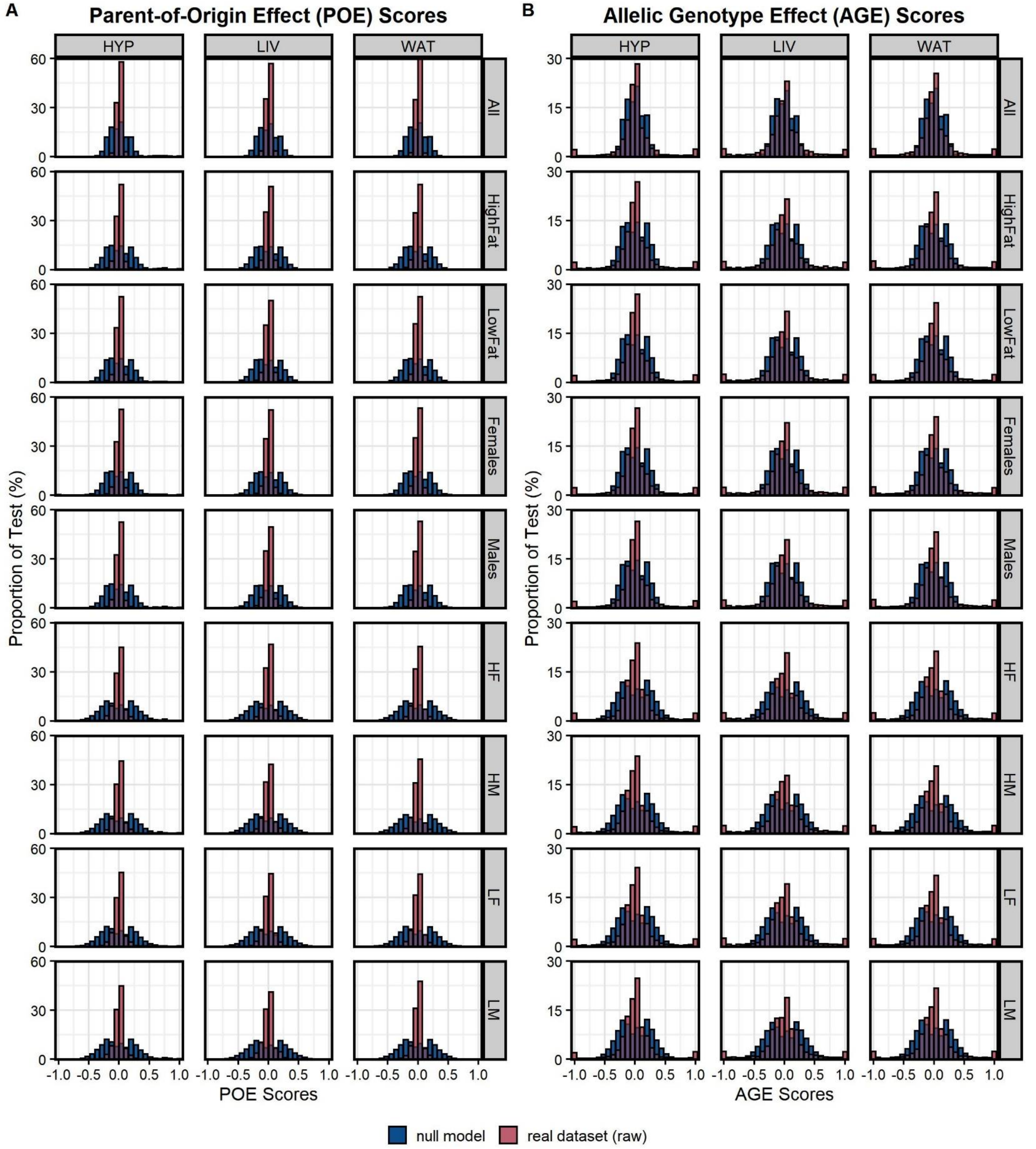

**Figure S24: POE and AGE score distributions between real and permuted datasets.**
Histograms of score distributions for the **(A)** Parent-of-Origin Effect (POE) and **(B)** Allelic Genotype Effect (AGE) scores in each tissue-by-context analysis. POE/AGE scores quantify the direction and magnitude of each gene's expression biases. Distributions are overlaid to highlight how the permuted/null models (blue) differ from the real datasets (pink): the null models have proportionally few tests (i.e. genes) with extreme scores near either tail. However, the real datasets have a high proportion of tests with scores near zero, indicating most genes are truly biallelic. The x-axis shows the range of scores (-1 to +1) and the y-axis shows the proportion of tests (%) in each bin. Columns are grouped by tissue and rows are grouped by environmental context.

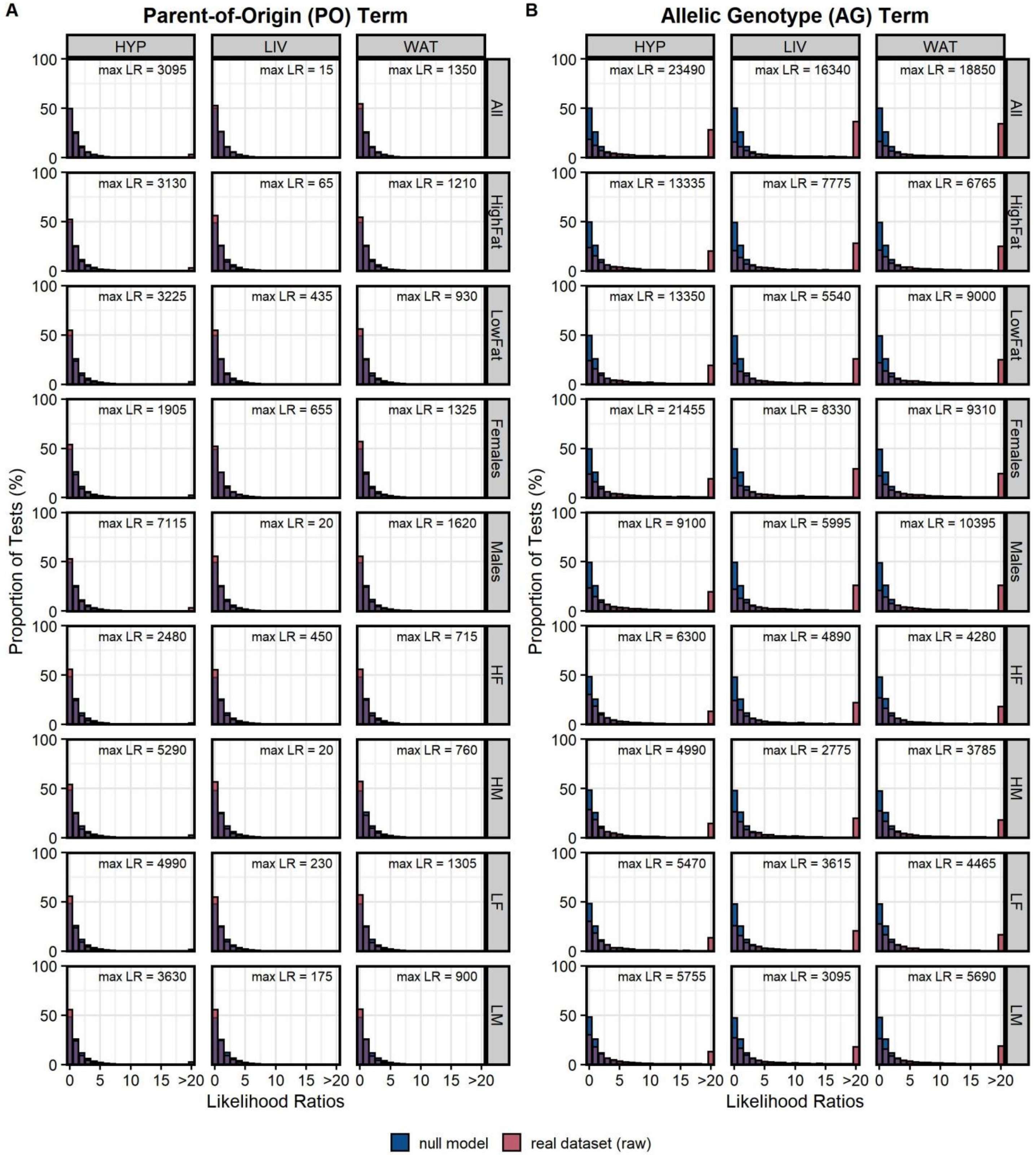

**Figure S25: Likelihood ratio distributions between real and permuted datasets for both terms.** Histograms of likelihood ratio (LR) distributions for the **(A)** Parent-of-Origin (PO) and **(B)** Allelic Genotype (AG) model terms in each tissue-by-context analysis. Distributions are overlaid to highlight how the permuted/null models (blue) differ from the real datasets (pink): the null models have a very low proportion of tests (i.e. genes) with large LRs (>20) as compared to the real datasets. Purple bars indicate where the two distributions overlap. The x-axis shows the range of LRs, where LRs >20 are collapsed into one bin for easier interpretability. The y-axis shows the proportion of tests (%) in each LR bin. Columns are grouped by tissue and rows are grouped by environmental context. The max LR is also denoted for each analysis.

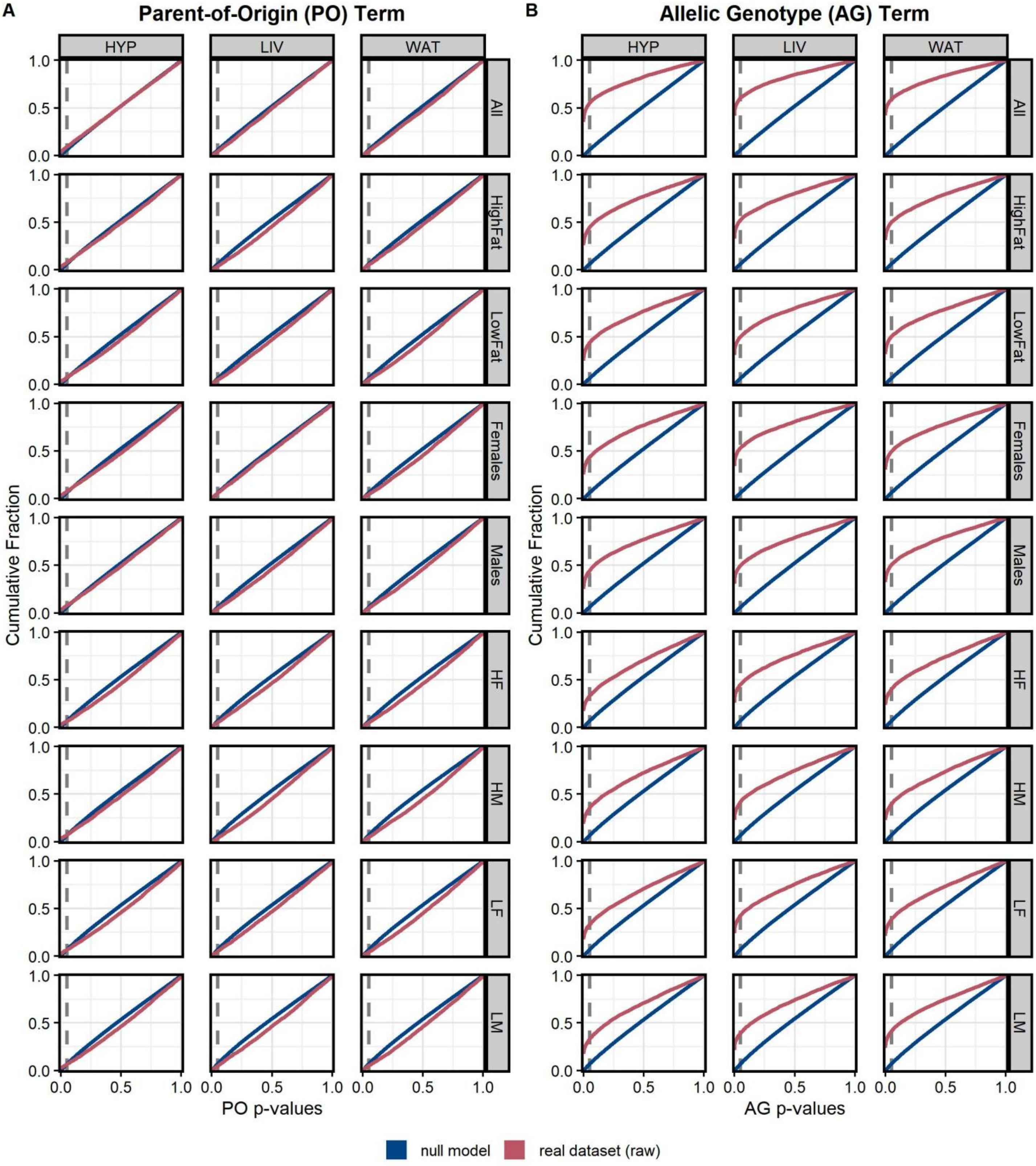

**Figure S26: Empirical cumulative distribution curves for the real and permuted datasets for both terms.** Density curves for the empirical cumulative distribution functions (ECDF) for the **(A)** Parent-of-Origin (PO) and **(B)** Allelic Genotype (AG) model terms in each tissue-by-context analysis. We corrected for multiple tests by fitting each term's raw p-values to its respective ECDF and computing adjusted p-values. Densities are overlaid to highlight how the permuted/null models (blue) differ from the real datasets (pink): the real datasets have a higher cumulative fraction of p-values below the significance threshold ($p \leq 0.05$) than the null models. The x-axis shows the range of p-values (0 to 1) and the y-axis shows the cumulative fraction of tests with p-values that are more extreme (smaller). Columns are grouped by tissue and rows are grouped by environmental context.

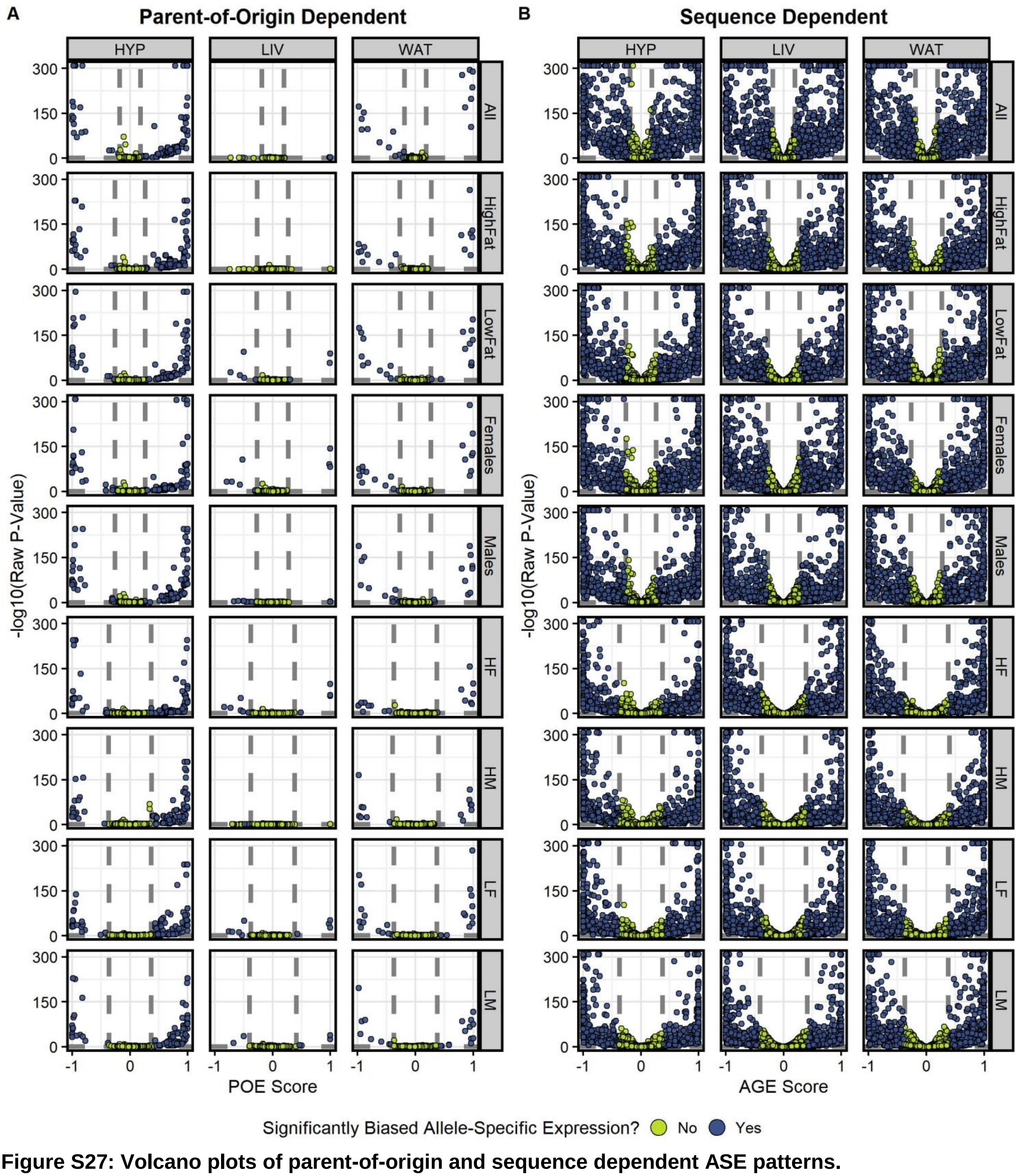

**Figure S27: Volcano plots of parent-of-origin and sequence dependent ASE patterns.**
Volcano plots summarizing the **(A)** parent-of-origin dependent and **(B)** sequence dependent ASE results in each tissue-by-context analysis. The x-axis shows the range of POE/AGE scores and the y-axis shows the -$\log_{10}$(raw p-value). Dashed lines indicate biological (effect scores) and statistical (p-values) significance thresholds. Genes exceeding both biological and statistical thresholds have significant ASE (blue). Most genes have no significant expression bias (green). Columns are grouped by tissue and rows are grouped by environmental context.
